# Structure of endogenous Pfs230:Pfs48/45 in complex with potent malaria transmission-blocking antibodies

**DOI:** 10.1101/2025.02.14.638310

**Authors:** Ezra T. Bekkering, Randy Yoo, Sophia Hailemariam, Fabian Heide, Danton Ivanochko, Matthew Jackman, Nicholas I. Proellochs, Rianne Stoter, Geert-Jan van Gemert, Ayana Maeda, Takaaki Yuguchi, Oscar T. Wanders, Renate C. van Daalen, Maartje R. Inklaar, Carolina M. Andrade, Pascal W.T.C. Jansen, Michiel Vermeulen, Teun Bousema, Eizo Takashima, John L. Rubinstein, Taco W.A. Kooij, Matthijs M. Jore, Jean-Philippe Julien

**Affiliations:** Department of Medical Microbiology, Radboud University Medical Center, The Netherlands; Program in Molecular Medicine, The Hospital for Sick Children Research Institute, Toronto, ON, Canada; Department of Biochemistry, University of Toronto, Toronto, ON, Canada; Division of Malaria Research, Proteo-Science Center, Ehime University, Matsuyama, Japan; Department of Molecular Biology, Faculty of Science, Oncode Institute, Radboud University Nijmegen, Nijmegen, The Netherlands; Division of Molecular Genetics, The Netherlands Cancer Institute, Amsterdam, The Netherlands; Department of Medical Biophysics, University of Toronto, Toronto, Ontario, Canada; Department of Immunology, University of Toronto, Toronto, Ontario, Canada

**Keywords:** Malaria, *Plasmodium falciparum*, Pfs230, Pfs48/45, cryo-electron microscopy, monoclonal antibodies, transmission-reducing activity, transmission-blocking vaccines

## Abstract

The Pfs230:Pfs48/45 complex forms the basis for leading malaria transmission-blocking vaccine candidates, yet li]le is known about its molecular assembly. Here, we used cryogenic electron microscopy to elucidate the structure of the endogenous Pfs230:Pfs48/45 complex bound to six potent transmission-blocking antibodies. Pfs230 consists of multiple domain clusters rigidified by interactions mediated through insertion domains. Membrane-anchored Pfs48/45 forms a disc-like structure and interacts with a short C-terminal peptide on Pfs230 that is critical for Pfs230 membrane-retention *in vivo*. Interestingly, membrane retention through this interaction is not essential for transmission to mosquitoes, suggesting that complex disruption is not a mode of action for transmission-blocking antibodies. Analyses of Pfs48/45-and Pfs230-targeted antibodies identify conserved epitopes on the Pfs230:Pfs48/45 complex and provides a structural paradigm for complement-dependent activity of Pfs230-targeting antibodies. Altogether, the antibody-bound Pfs230:Pfs48/45 structure presented improves our molecular understanding of this biological complex, informing the development of next-generation *Plasmodium falciparum* transmission-blocking interventions.

## INTRODUCTION

Nearly half of the human population is at risk for malaria infection, making malaria one of the largest public health concerns (1). The parasite that is responsible for the majority of fatal human cases, *Plasmodium falciparum*, is efficiently transmitted by *Anopheles* mosquitoes throughout the population (2). Human-to-mosquito transmission is mediated by mature male and female gametocytes that are taken up by an *Anopheles* mosquito during a bloodmeal (3). Inside the mosquito midgut, male and female gametocytes rapidly activate and egress from their red blood cells (RBCs) as micro-and macrogametes, respectively. Microgametes fuse with macrogametes creating a zygote that forms the basis for further parasite development in the mosquito, eventually resulting in an infectious mosquito that can further spread the parasite and its related disease (4).

Two sexual-stage surface proteins essential for *Plasmodium* transmission, Pfs230 (5) and Pfs48/45 (6), are expressed in both gametocytes and gametes. Pfs48/45 and Pfs230 knockout (KO) parasites showed markedly lower oocyst formation rates, demonstrating that these proteins play a critical role in establishing mosquito infection (7, 8). Pfs48/45 is predicted to have a GPI-anchor and localizes together with Pfs230 to the parasite plasma membrane (PPM) (9). Pfs230, without a recognizable GPI anchor or transmembrane motif, is thought to associate with the PPM by forming a stable heterodimeric complex with Pfs48/45, as shown in co-immunoprecipitation studies (10, 11). This is further supported by the observation that Pfs48/45 KO parasites lose Pfs230 surface retention (7). Both proteins are part of the 6-Cysteine (6-Cys) protein family, characterized by the presence of 6-Cys domains: Immunoglobulin-like folds that contain up to six cysteines that can form disulfide bonds (12, 13). Pfs230 contains fourteen 6-Cys domains, whereas Pfs48/45 contains three (**Figure 1A**). Structural information on the full Pfs230:Pfs48/45 structure has been limited; two full-length structures of recombinant Pfs48/45 have been reported, adopting two distinct conformations – a disc-like (14) and an extended (15) conformation – whereas the structural characterization of Pfs230 has been limited to just the first two domains of the protein (16–20). Additionally, there is a lack of structural understanding of what domains are involved in the Pfs230:Pfs48/45 interaction itself.

**Figure 1:**
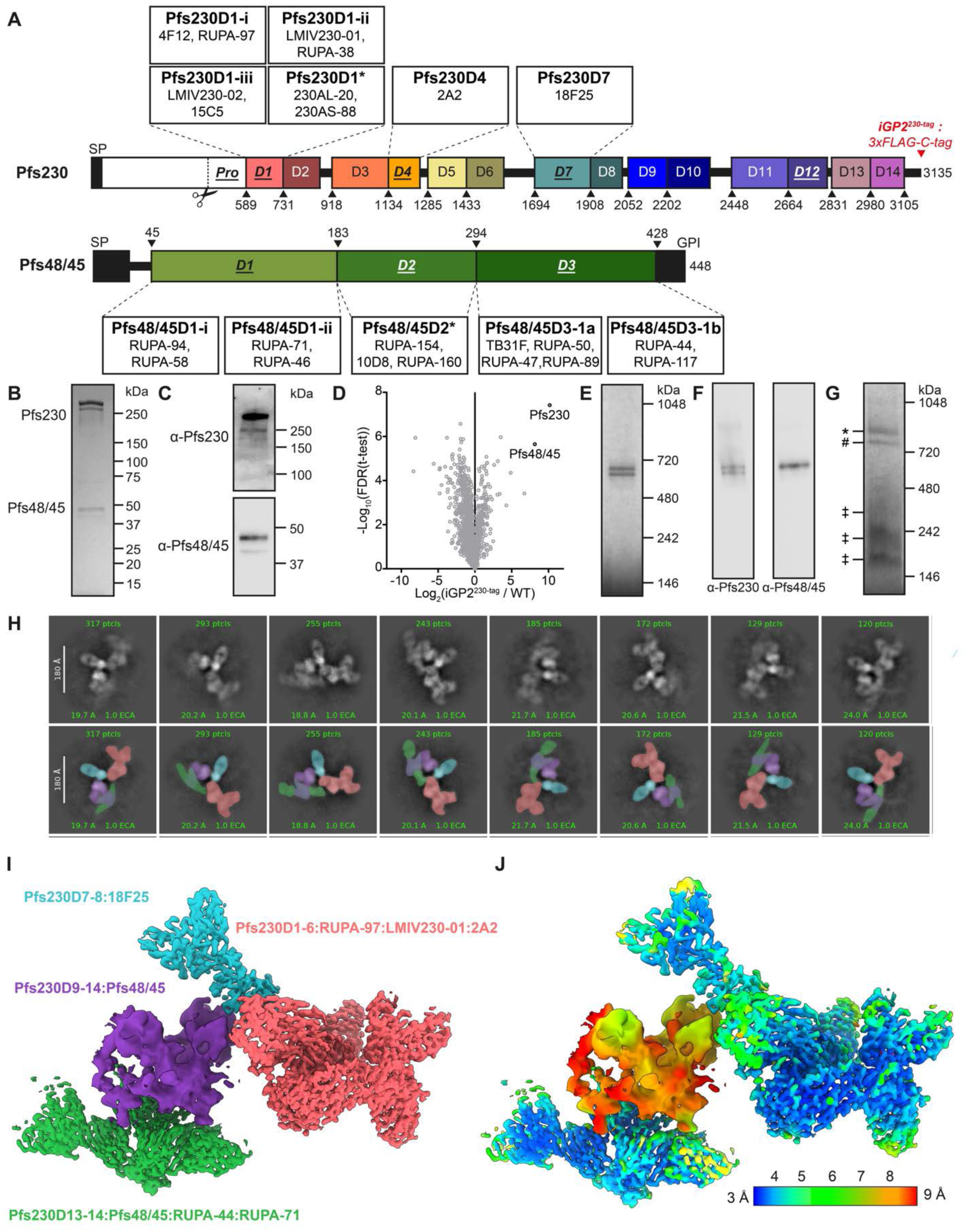
**Purification of Pfs230:Pfs48/45:6Fab complex from *P. falciparum* gametocytes**. **(A)** Schematic illustration of Pfs230 and Pfs48/45 domains (not to scale), epitopes found within these domains, and examples of mAbs targeting these epitopes (see also **Table S1**). Asterisks indicate multiple additional distinct epitopes at the indicated domain. Domains that contain known transmission-blocking epitopes are indicated by underlined italic. Arrows indicate amino acid number of the approximate start of the domain as determined by (*13*). SP = Signal peptide, GPI = Glycosylphosphatidylinositol anchor site. **(B-C)** Coomassie Blue (B) and immunoblot (C) staining of a denaturing SDS-PAGE gel of FLAG-tag purified Pfs230:Pfs48/45 complex. **(D)** Mass spectrometry analysis of FLAG-tag purified Pfs230:Pfs48/45 complex. The volcano plot shows the log_2_ fold change plotted against the log_10_ fold false discovery rate, comparing FLAG-tag co-immunoprecipitation of iGP2^230-tag^ versus NF54/WT parasites. **(E-G)** Blue-Native PAGE analysis, showing (E) Coomassie blue and (F) α-Pfs230 and α-Pfs48/45 immunoblot staining of neat purified Pfs230:Pfs48/45 complex, or (G) Coomassie blue staining of eluted complex incubated with molar excess of six Fabs (RUPA-97, LMIV230-01, 2A2, 18F25, RUPA-44, and RUPA-71). Symbols: * = Pfs230:Pfs48/45:6Fab; # = Pfs230:4Fab; ‡ = excess Fab fragment. **(H)** Eight representative 2D class averages generated from cryo-EM of HPLC-SEC purified Pfs230:Pfs48/45:6Fab complex. Structures are annotated in the lower row, with orange highlighting Pfs230D1-6:RUPA-97:LMIV230-01:2A2, blue Pfs230D7-8:18F25, purple Pfs230D-13-14:Pfs48/45, and green Pfs230D13-14:Pfs48/45:RUPA-44:RUPA-71. **(I)** Composite CryoEM map of the Pfs230:Pfs48/45 complex coloured according to the same colouring scheme. **(J)** Local resolution of each map coloured according to scale. Maps are contoured to 5 Å around built models. *See also Figure S1-3 and Table S1*.

The human-to-mosquito transmission stages form a developmental bottleneck for the malaria parasite and are therefore an attractive target for interventions seeking malaria elimination (21), such as transmission-blocking vaccines (TBVs). Indeed, modeling studies have suggested that TBVs could synergistically increase the effectiveness of vaccines targeting other parasite stages (22–24). Importantly, both Pfs230 and Pfs48/45 were recognized in the early 1980s as the targets for antibodies that had potent human-to-mosquito transmission-reducing activity (TRA) (9, 11, 25). Since then, multiple studies have shown that naturally infected individuals can acquire Pfs230 and Pfs48/45-targeted antibodies that are associated with high-level TRA in serum (*e.g.* (26–31)) and these antibodies can block transmission when purified (26). The structure, epitopes, and TRA of these naturally acquired antibodies have been partially characterized at the monoclonal antibody level (15, 17, 20, 32). Domain 1 of Pfs230 (Pfs230D1) and domain 3 of Pfs48/45 (Pfs48/45D3) are the targets of the most potent transmission-blocking monoclonal antibodies (mAbs) (*e.g.* (15, 17, 19, 20, 32–35)). Initial reports suggested that other domains of Pfs230 and Pfs48/45 were not able to elicit comparably potent transmission-blocking antibodies (35). Recently at least three other Pfs230 domains, and Pfs48/45D1, have been identified as targets for potent transmission-blocking antibodies, but structural insight into these epitopes is lacking in comparison to Pfs230D1 and Pfs48/45D3 (32, 36–39).

Regardless of the domain they target, the vast majority of the potent α-Pfs230 antibodies is complement-dependent (17, 20, 36, 40), although some antibodies retain TRA at very high concentrations in the absence of complement (16, 34). In contrast, α-Pfs48/45 antibodies can block transmission at low concentrations in a complement-independent manner (15, 18, 32, 33, 38, 41). The rationale behind what drives antibody potency for both α-Pfs230 and α-Pfs48/45 antibodies remains unclear. As both proteins are capable of eliciting highly potent antibodies, they have been targeted as lead antigens and are currently the most advanced and promising TBVs in development (42, 43). Indeed, immunogens based off these two proteins have been extensively evaluated in preclinical and phase I/II clinical trials (44–47). Although these clinical trials show promising results, induced TRA was incomplete and more efficacious vaccines may thus be needed. Structure-function relationships of how potent antibodies target the Pfs230:Pfs48/45 complex would not only enhance our understanding of *P. falciparum* transmission biology and antibody mechanism of actions, but also aid in the design and development of next-generation malaria vaccines that can induce more potent TRA to reach elimination goals.

Here, we report the molecular structure of the endogenous Pfs230:Pfs48/45 heterodimer in complex with six antibodies with potent TRA as determined by cryogenic electron microscopy (cryo-EM). Cumulatively, our data provide critical molecular insights into the Pfs230:Pfs48/45 heterodimer complex, moving towards deepening our understanding of *P. falciparum* transmission biology and opportunities to block this process through next-generation biomedical interventions.

## RESULTS

### Structure of the Pfs230:Pfs48/45 complex

We set out to determine the cryo-EM structure of the Pfs230:Pfs48/45 complex by purifying it from mature *P. falciparum* gametocytes. Using the inducible gametocyte producing parasite line NF54/iGP2 (48), we fused *pfs230* (Pf3D7_0209000) C-terminally with a 3xFLAG-C-tag coding sequence, yielding parasite line iGP2^230-tag^ (**Figure S1A-D**). iGP2^230-tag^ gametocytes showed normal Pfs230 and Pfs48/45 localization and retained normal *in vitro* exflagellation and mosquito infectivity in standard membrane feeding assays (SMFA; **Figure S1E-F**). The Pfs230:Pfs48/45 complex was purified using anti-FLAG resin. Pfs230 and Pfs48/45 were the highest enriched proteins from this purification as detected by LC-MS/MS (**Figure 1B-D**). Two high-molecular weight bands were observed in Blue-Native Polyacrylamide Gel Electrophoresis (BN-PAGE) **(Figure 1E)**. Both bands were stained by α-Pfs230 antibodies in immunoblotting assays, but only one band was stained by α-Pfs48/45 antibodies, attributing these two bands to the Pfs230:Pfs48/45 complex and Pfs230 alone, respectively (**Figure 1F**). BN-PAGE analysis of eluted protein incubated with a molar excess of different fragment of antigen-binding (Fab) fragments of transmission-blocking antibodies, targeting a range of conformational epitopes on Pfs48/45 and Pfs230, confirmed the native structure of the eluted proteins (**Figure S1G**). While α-Pfs230 Fabs shifted both protein species upwards, α-Pfs48/45 Fabs only affected the upper band, further ascertaining the top band as the Pfs230:Pfs48/45 complex.

Since we were interested in how potent transmission-blocking antibodies are targeting the native Pfs230:Pfs48/45 complex, we generated a size exclusion chromatography (SEC)-purified Pfs230:Pfs48/45 heterodimer in complex with six Fab fragments, using the structurally delineated RUPA-97 (20), LMIV230-01 (19), and RUPA-44 (32), and the yet-to-be-structurally delineated 2A2 (38, 40, 49), 18F25 (36, 49), and RUPA-71 (32) inhibitory antibodies (**Figure 1G, Figure S1H**). Cryo-EM data of this Pfs230:Pfs48/45:6Fab complex was collected at 0°, 35°, and 40° tilt. 2D class averages from this dataset clearly revealed the presence of all components of the complex: Pfs230 bound to four Fabs (LMIV230-01, RUPA-97, 2A2, 18F25) in complex with Pfs48/45 bound to two Fabs (RUPA-44 and RUPA-71) (**Figure 1H**).

The highest resolution structural information was derived by generating four locally refined maps corresponding to more rigid components of the complex: Pfs230D1-6 bound to RUPA-97, LMIV230-01, and 2A2 at a global resolution of 3.6 Å; Pfs230D7-8 bound to 18F25 at a global resolution of 3.8 Å; Pfs230D9-14 bound to Pfs48/45 at a global resolution of 4.7 Å; and Pfs230D13-14 bound to Pfs48/45, RUPA-44, and RUPA-71 at a global resolution of 3.4 Å (**Figure 1I-J, Figure S2-3, Table 1**).

**Table 1.**
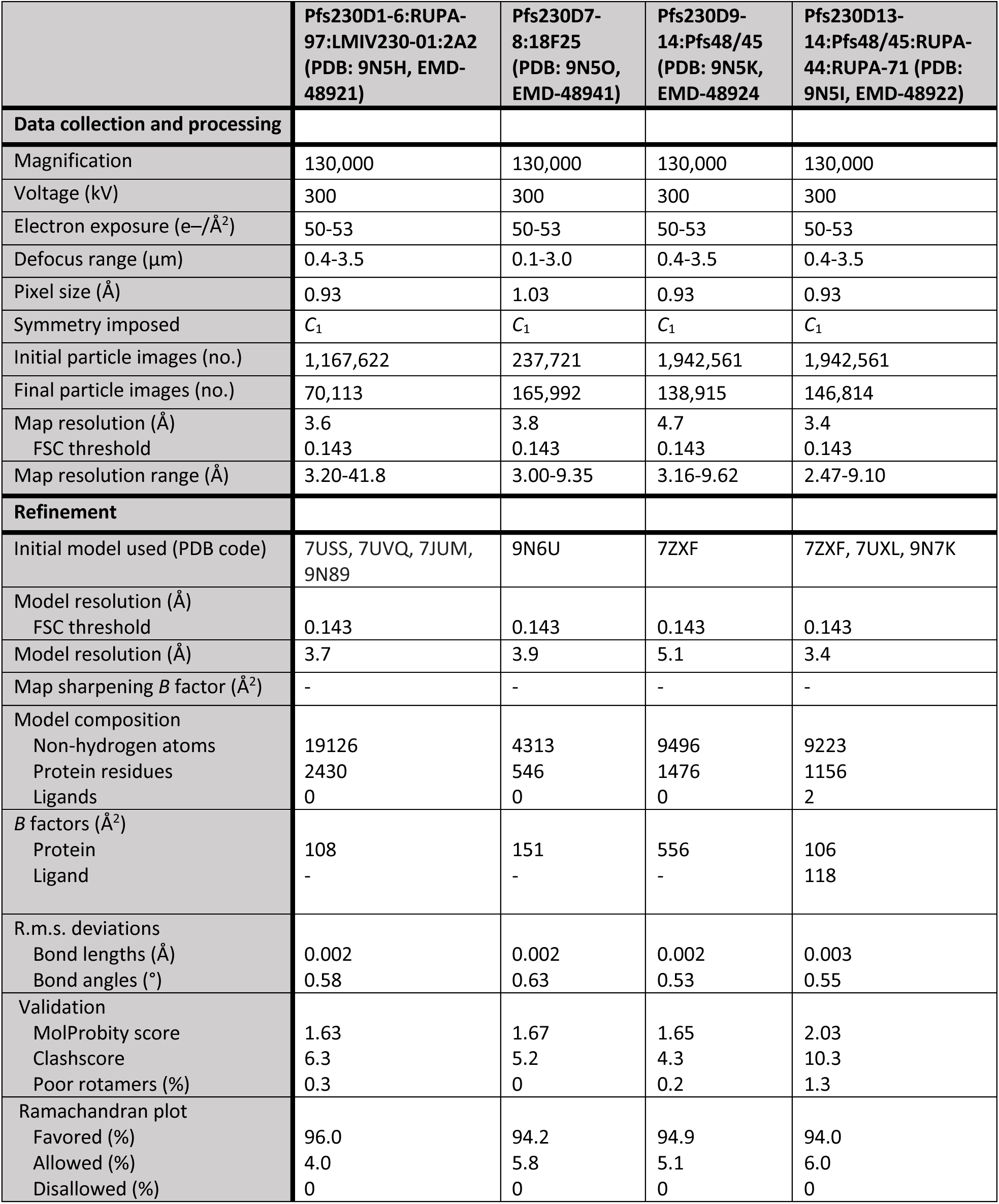
CryoEM data collection and refinement statistics.

|  | Pfs230D1-6:RUPA-97:LMIV230-01:2A2 (PDB: 9N5H, EMD-48921) | Pfs230D7-8:18F25 (PDB: 9N5O, EMD-48941) | Pfs230D9-14:Pfs48/45 (PDB: 9N5K, EMD-48924) | Pfs230D13-14:Pfs48/45:RUPA-44:RUPA-71 (PDB: 9N5I, EMD-48922) |
| --- | --- | --- | --- | --- |
| **Data collection and processing** |  |  |  |  |
| Magnification | 130,000 | 130,000 | 130,000 | 130,000 |
| Voltage (kV) | 300 | 300 | 300 | 300 |
| Electron exposure (e-/Å<sup>2</sup>) | 50-53 | 50-53 | 50-53 | 50-53 |
| Defocus range (μm) | 0.4-3.5 | 0.1-3.0 | 0.4-3.5 | 0.4-3.5 |
| Pixel size (Å) | 0.93 | 1.03 | 0.93 | 0.93 |
| Symmetry imposed | C<sub>1</sub> | C<sub>1</sub> | C<sub>1</sub> | C<sub>1</sub> |
| Initial particle images (no.) | 1,167,622 | 237,721 | 1,942,561 | 1,942,561 |
| Final particle images (no.) | 70,113 | 165,992 | 138,915 | 146,814 |
| Map resolution (Å) | 3.6 | 3.8 | 4.7 | 3.4 |
| FSC threshold | 0.143 | 0.143 | 0.143 | 0.143 |
| Map resolution range (Å) | 3.20-41.8 | 3.00-9.35 | 3.16-9.62 | 2.47-9.10 |
| **Refinement** |  |  |  |  |
| Initial model used (PDB code) | 7USS, 7UVQ, 7JUM, 9N89 | 9N6U | 7ZXF | 7ZXF, 7UXL, 9N7K |
| Model resolution (Å) |  |  |  |  |
| FSC threshold | 0.143 | 0.143 | 0.143 | 0.143 |
| Model resolution (Å) | 3.7 | 3.9 | 5.1 | 3.4 |
| Map sharpening B factor (Å<sup>2</sup>) | - | - | - | - |
| Model composition |  |  |  |  |
| Non-hydrogen atoms | 19126 | 4313 | 9496 | 9223 |
| Protein residues | 2430 | 546 | 1476 | 1156 |
| Ligands | 0 | 0 | 0 | 2 |
| B factors (Å<sup>2</sup>) |  |  |  |  |
| Protein | 108 | 151 | 556 | 106 |
| Ligand | - | - | - | 118 |
| R.m.s. deviations |  |  |  |  |
| Bond lengths (Å) | 0.002 | 0.002 | 0.002 | 0.003 |
| Bond angles (°) | 0.58 | 0.63 | 0.53 | 0.55 |
| Validation |  |  |  |  |
| MolProbity score | 1.63 | 1.67 | 1.65 | 2.03 |
| Clashscore | 6.3 | 5.2 | 4.3 | 10.3 |
| Poor rotamers (%) | 0.3 | 0 | 0.2 | 1.3 |
| Ramachandran plot |  |  |  |  |
| Favored (%) | 96.0 | 94.2 | 94.9 | 94.0 |
| Allowed (%) | 4.0 | 5.8 | 5.1 | 6.0 |
| Disallowed (%) | 0 | 0 | 0 | 0 |

### Insertion domains organize Pfs230 domains

Molecular models were built into each of the locally refined maps to elucidate structural features of the interdomain topologies and dispositions of potent mAbs. The models derived from the maps of Pfs230D1-6:RUPA-97:LMIV230-01:2A2 and Pfs230D7-8:18F25 (**Figure 1I-J**, **Table 1**) were initialized from experimentally determined crystal structures and predicted AlphaFold (AF) models, which were refined into maps with sufficiently high resolution to model sidechain positions (**Table 1, Table S3**). For the 4.7 Å resolution map of Pfs230D9-14:Pfs48/45, we report the backbone positions of a compositely refined Pfs230D9-12 structure in the presence of the Pfs230D13-14:Pfs48/45 structure discerned from the 3.4 Å resolution map (**Figure 1I-J**, **Table 1**). AF models of Pfs230D9-D12 were docked and manually rearranged into the Pfs230D9-14:Pfs48/45 map. A template-guided inference strategy in AF2 followed by refinement using map-constrained energy minimizations resulted in the final model. We attribute the lower local resolution of Pfs230D9-12 in our Pfs48/45:Pfs230D9-14 map to intrinsic flexibility between the D12 and D13 domains and to bound antibodies on Pfs48/45 that may create steric hindrance in this region (**Figure S3**).

Each tandem structural unit of Pfs230 consists of A-and B-type 6-Cys domains (**Figure S4**). Most A-type domains feature a 5-on-5 β-sandwich whereas B-type domains feature a 4-on-5 β-sandwich fold **(Figure S4A, inset**). A key differentiating feature of these folds is the way in which the first two β-strands are arranged (**Figure 2A, Figure S4A, inset**). In the A-type folds (i.e. Pfs230 D1, D3, D5, D7, D9, D11, and D13), the first canonical β1 strand is split between the two β-sheets (β1 and β1’) of the 6-Cys β-sandwich fold before transitioning into the β2 strand. In contrast, in the B-type folds (i.e. Pfs230 D2, D4, D6, D8, D10, D12 and D14), the first two β-strands (β1 and β2) are part of the same β-sheet.

**Figure 2.**
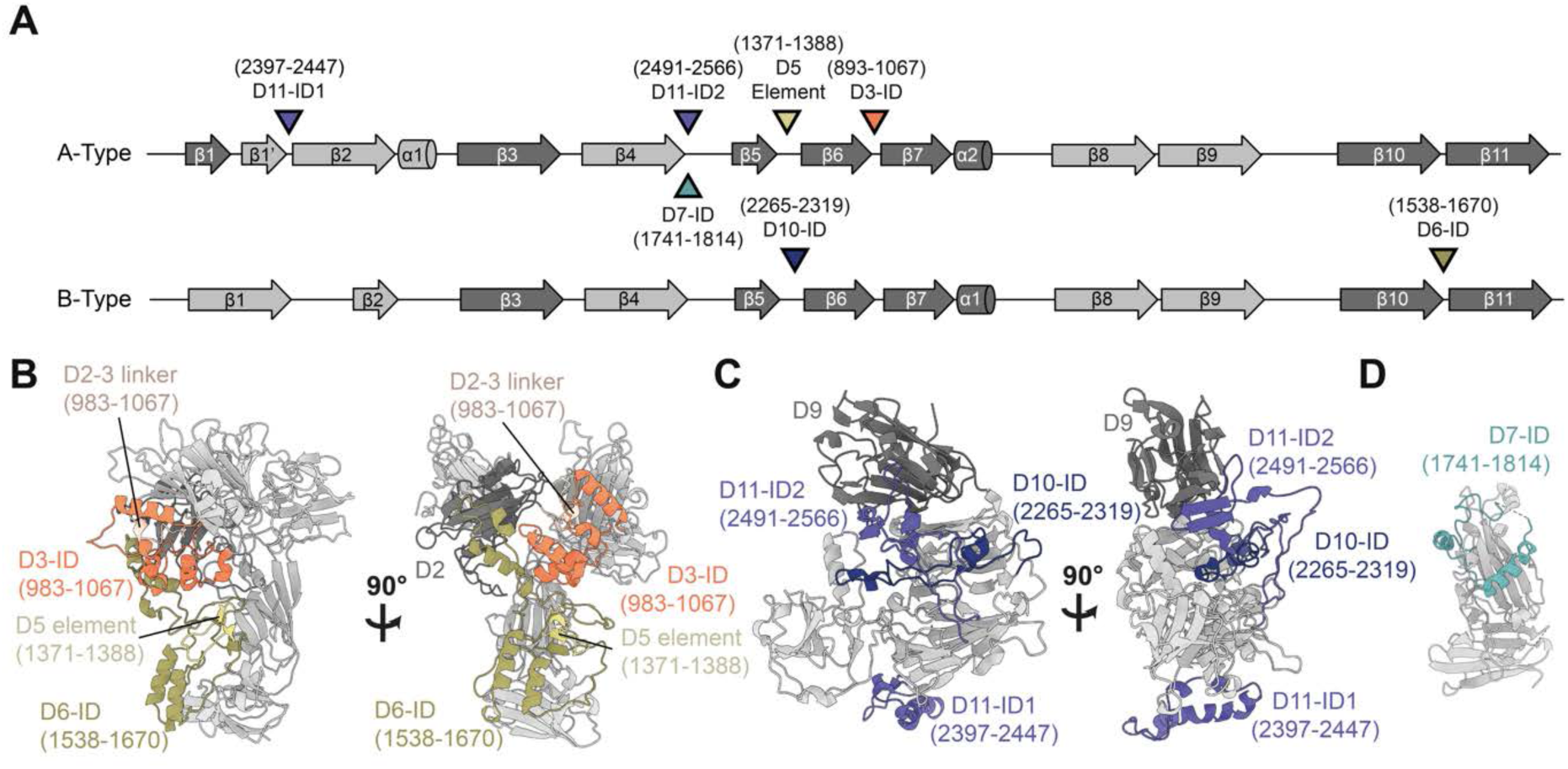
Insertion domains (IDs) facilitate Pfs230 domain clustering. **(A)** Secondary structural elements of A-and B-type domains, with insertion domains (IDs) or non-canonical structural element (D5 element) found in respective domains indicated by coloured arrows and with residue range in brackets. **(B)** The D2-D3 linker (light orange), D3-ID (orange), D5 structural element (yellow), and D6-ID (dark yellow) interact with one another, forming a structural fold that packs against the base of D2 (dark grey). **(C)** The D10-ID (blue) and Pfs230D11-ID2 (purple) interact with one another, positioning the D11-ID2 (purple) to form an extended β-sheet with D9 (dark grey). **(D)** D7-8 (silver) and D7-ID (cyan) lack interdomain contact. *See also Figure S4*.

While small deviations from the canonical 6-Cys domains exist, primarily near the perpendicular β-hairpin (**Figure S4B-C**), most of the structural diversity that distinguishes the different Pfs230 6-Cys domains lie in long sequence insertions in loop regions between the β-strands **(Figure 2A)**. These regions were previously coined as insertion domains (IDs) for another 6-Cys domain-containing protein (50, 51) and are present in Pfs230D3, D6, D7, D10 and D11 (**Figure S4D-I**). Many of these IDs (i.e. Pfs230D3, D6, D10, and D11-ID2) contribute to interdomain contacts (**Figure 2B-C**), while others (i.e. Pfs230D11-ID1 and D7) were not observed to interact with other domains (**Figure 2D**). The presence of these IDs mediates the formation of domain clusters in Pfs230.

### The Pfs230 C-terminus interacts with Pfs48/45

The interaction site between Pfs48/45 and Pfs230 was delineated at 3.4 Å resolution (**Figure 3A, Figure S3C, Table 1**). Our structure indicates that Pfs230 engages with all three domains of Pfs48/45 through its terminal 6-Cys domains, Pfs230D13 (296 Å^2^) and Pfs230D14 (1376 Å^2^) (**Figure 3B**). Pfs48/45 adopts a disc-like conformation in our structure with Pfs230, with Pfs48/45D1 (1057 Å^2^) and Pfs48/45D3 (478 Å^2^) accounting for the majority of contacts with Pfs230, while Pfs48/45D2 (112 Å^2^) contributes more modestly (**Figure 3B, Figure S5A, Table S2**). Pfs230 binds to Pfs48/45 on the opposite face of the disc conformation compared to the C-terminus of Pfs48/45, where the GPI linker is located for attachment to the PPM (**Figure 3B**). Previously, two different conformations of Pfs48/45 have been observed in the absence of Pfs230 binding, a compact, disc-like structure solved using X-ray crystallography (14) and an elongated Pfs48/45 structure determined by cryo-EM as bound to a different set of antibodies (15). A comparison of these models with our Pfs230-bound Pfs48/45 structure indicates very similar arrangements of Pfs48/45D1-2 (backbone RMSD = 1.4-1.7 Å for res. 45-288) (**Figure S5B**). However, the position of Pfs48/45D3 relative to Pfs48/45D1-2 varies considerably more. When aligned to Pfs48/45D1-2, there is almost no overlap between Pfs48/45D3 from Pfs230-bound Pfs48/45 and the elongated Pfs48/45 structure while modest positional differences are observed between Pfs48/45D3 of the two disc-like Pfs48/45 structures (backbone RMSD = 3.9 Å for res. 293-428; **Figure S5B**).

**Figure 3:**
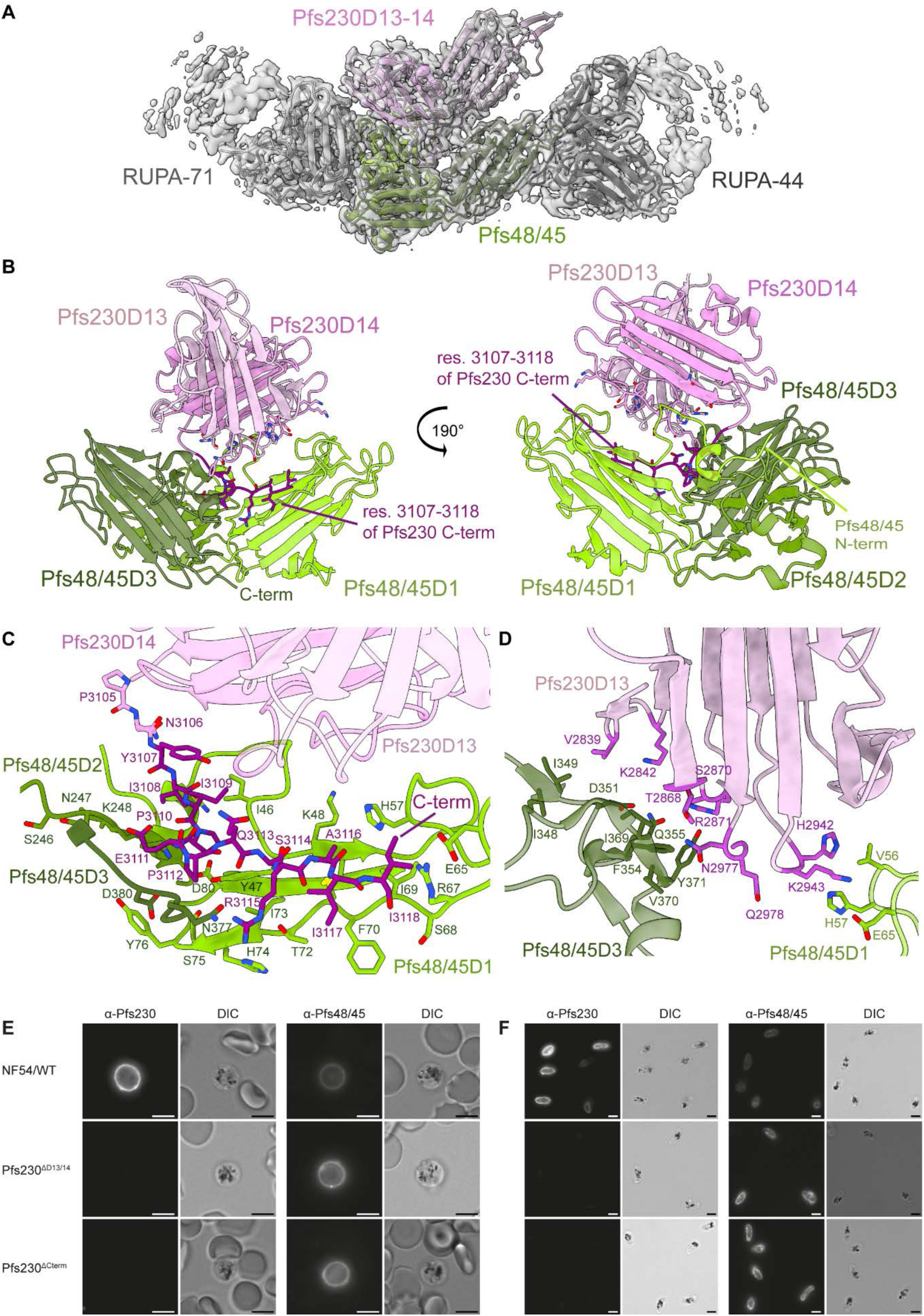
**Molecular description of the Pfs230 and Pfs48/45 interaction**. **(A)** Overview of the structure of Pfs48/45 (yellow green, olive drab, and dark olive drab for domains 1-3, respectively) bound to Pfs230D13-14 (plum), RUPA-71 Fab (dark grey), and RUPA-44 Fab (light grey) with the cryo-EM map shown in grey. **(B)** Pfs48/45 bound to Pfs230D13-14 with key Pfs230 contact residues shown as sticks and the C-terminal region of Pfs230 highlighted in dark magenta. **(C-D)** binding interactions between Pfs48/45 and Pfs230D14-Cterm (C) and Pfs230D13 (D) with key residue contacts shown as sticks. **(E-F)** Representative suspension immunofluorescence microscopy images of NF54/WT, 230^ΔD13/14^, and 230^ΔCterm^ female macrogametes (E) and saponin-treated stage V gametocytes (F). Parasites were incubated with 15 µg/ml 18F25-DyLight488 (α-Pfs230) or 45.1-DyLight488 (α-Pfs48/45). All images per experiment were taken and analyzed with the same settings. Scale bar represents 5 µm. *See also Figure S5-7 and Table S2*.

The C-terminus of Pfs230 (Pfs230Cterm; res. 3108-3118) – a sequence extending out of the canonical 6-Cys domain – is integral for mediating the Pfs230:Pfs48/45 interaction (**Figure 3C, Figure S6A**). These residues occupy a groove of the Pfs48/45 disc and form extensive interactions with loop 45-49 and strand 65-76 of Pfs48/45D1, loop 246-250 of Pfs48/45D2, and loop 377-380 of Pfs48/45D3 (**Figure 3B**). While accounting for fewer interactions than Pfs230D14, Pfs230D13 residues 2866-2871 and the intervening loop between Pfs230D13 and Pfs230D14 (res. 2977-2984) support complexation by engaging with a portion of a Pfs48/45D3 extended loop (res. 348-371) (**Figure 3D**). Additionally, the Pfs48/45 N-terminus wraps around Pfs48/45D1-2 and, through residues 41-50, forms interactions with loops 2993-2998 and 3108-3114 and strand 3007-3014 of Pfs230D14 (**Figure 3B**). While the N-terminal region of Pfs48/45 remained largely unresolved in previous structural studies of Pfs48/45 without Pfs230 implying flexibility (14, 15), this region is well-resolved in the Pfs230-bound structure and reveals that unlike other A-type domains, Pfs48/45D1 features a different β1 orientation and connectivity (**Figure S5C**), localizing to the interface. An analysis of single nucleotide polymorphisms (SNP) across *P. falciparum* revealed that the interface between Pfs48/45 and Pfs230 is highly conserved (**Figure S5A, S5D**).

Corroborating these structural findings, a recombinantly produced protein construct spanning Pfs230D13-14-Cterm bound to female gametes on which Pfs48/45 is anchored, while similarly produced constructs spanning either Pfs230D1 (previously established to be correctly folded and capable of inducing transmission-blocking immunity (36)) or Pfs230D10 could not (**Figure S7A**). To further confirm the interaction between Pfs230D13-14-Cterm and Pfs48/45, two transgenic parasite lines carrying a truncated *pfs230* gene were created (**Figure S7B-E**). Pfs230^ΔD13/14^ parasites express Pfs230 up until Pfs230D12, while Pfs230 expressed in Pfs230^ΔCterm^ gametocytes only lacks the last 29 C-terminal residues (res. 3107 – 3135), seen as central to the Pfs230:Pfs48/45 interface in our cryo-EM structure. Wildtype Pfs230 is expected to localize on the outside of the PPM, facing into the parasitophorous vacuole, and is expected to co-localize with PPM-bound Pfs48/45. Western blot analysis shows that late-stage Pfs230^ΔD13/14^ and Pfs230^ΔCterm^ gametocytes still contain (truncated) Pfs230. However, saponin-treated parasites with a disrupted parasitophorous vacuole membrane (PVM) lose all Pfs230, demonstrating that Pfs230 is no longer retained on the PPM (**Figure S7F-G**). In fixed immunofluorescence microscopy, full-length and truncated Pfs230 proteins localized in close proximity to Pfs48/45, suggesting that Pfs230 protein trafficking is unaffected in both mutant parasite lines (**Figure S7H**). When the PVM was naturally disrupted upon activation of macrogametes or chemically disrupted in saponin-treated gametocytes, Pfs230 remained bound on the PPM surface in wildtype parasites. In contrast, both Pfs230^ΔD13/14^ and Pfs230^ΔCterm^ gametocytes and gametes lacked surface-bound Pfs230 in these experiments (**Figure 3E-F**). Furthermore, we could not find evidence of a Pfs230-positive population in Pfs230^ΔCterm^ gametes in a flow cytometry-based binding assay using over 2,000 macrogametes (**Figure S7I-J**). Together, our structural and *in vivo* experiments demonstrate that the C-terminal region of Pfs230 is essential for the formation of the Pfs230:Pfs48/45 complex.

### Structural characterization of the Pfs48/45D1-ii epitope

The X-ray crystal structure of unliganded RUPA-71 was solved at 2.3 Å resolution (**Table S3**) to use as a high-resolution model to fit and refine into the 3.4 Å resolution Pfs230D13-14:Pfs48/45:RUPA-44:RUPA-71 cryo-EM map (**Figure 4A-B**, **Table 1**). RUPA-71 predominantly interacts with strand 126-135 of Pfs48/45D1, with additional contacts on Pfs48/45D1 loop 88-90 (**Figure 4A-B, Figure S6B, Table S4**). This is consistent with the previous epitope mapping derived from HDX-MS (15). Heavy chain Complementarity-Determining Region 1 (HCDR1; 108 Å^2^), HCDR2 (66 Å^2^), HCDR3 (472 Å^2^), and Kappa chain CDR1 (KCDR1; 52 Å^2^) of RUPA-71 all contact Pfs48/45D1, with the extended HCDR3 (20 aa) contributing most key contacts (**Figure 4A, Table S4)**. The epitope of RUPA-71 is highly charged with several negative and positive patches. RUPA-71 binds to these regions through salt bridges formed between RUPA-71 KCDR1 residue K^32^ and HCDR1-3 residues D^31^ and R^98^ and E_129_, K_89_, K_101_, E_130_, and D_132_ of Pfs48/45D1 (**Figure 4A, Table S4)**. Noticeably, the HCDR3 of RUPA-71 adopts a β-hairpin that forms a β-sheet-like structure with two β-strands of Pfs48/45D1 (res. 126-143) (**Figure 4B**). In addition to intra-chain hydrogen bonds, this β-sheet is held together by five backbone hydrogen bonds between RUPA-71 G^96^, R^98^, and Y^100^ and D_132_, E_130_, and I_128_ of Pfs48/45D1 (**Figure 4B, Table S4**). Sequence analysis across parasite field isolates revealed only five rare SNPs within the RUPA-71 epitope: K33T (0.015%), K33N (0.007%), S90R (0.007%), K101T (0.029%), and S119T (0.007%), making this epitope highly conserved (**Figure S8A**) (*38*). When compared to RUPA-58, which is the only other D1 antibody that has been structurally characterized to date, RUPA-71 binds to a largely separate epitope, although with some overlap in residue contacts (K_88_, K_89_, S_90_, K_101_, D_132,_ and R_140_) that would result in steric clashes (**Figure S8B**). RUPA-58 and RUPA-71, as well as other antibodies that compete with them, require low concentrations to inhibit in the SMFA (IC_80_ < 10 μg/ml), indicating that mAbs that bind to this region of Pfs48/45D1 (res. 86-101 and 126-139) are potently inhibitory (15, 32).

**Figure 4:**
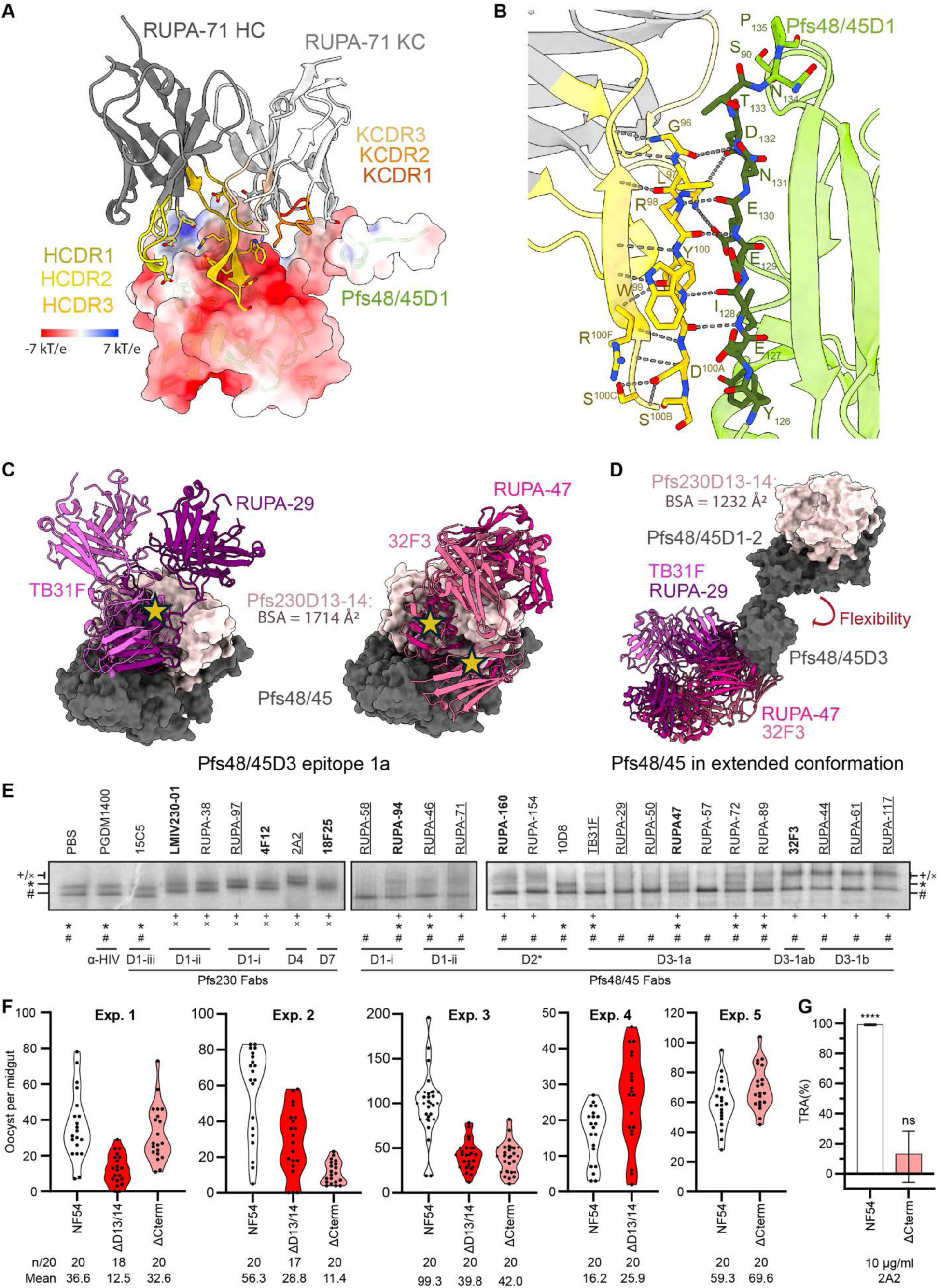
**Molecular characterization of anti-Pfs48/45 epitopes in the Pfs230-bound state of Pfs48/45**. **(A)** Structure of RUPA-71 bound to Pfs48/45D1 depicted as surface and coloured by electrostatic potential calculated by Adaptive Poisson-Boltzmann Solver (APBS) and visualized in ChimeraX. **(B)** RUPA-71 HCDR3 (gold) bound to Pfs48/45D1 (yellow green) with key residues shown as sticks and hydrogen bonds or electrostatic interactions indicated by dashed lines. **(C-D)** Model of Pfs48/45 (dark gray, depicted in surface) bound to Pfs230D13-14 (misty rose, depicted in surface) in the disc-like (C) or extended (D) conformation bound to Pfs48/45D3-1a binders (TB31F (orchid, PDB ID: 6E63), RUPA-29 (dark magenta, PDB ID: 7UXL), RUPA-47 (violet red, PDB ID: 7UNB), and 32F3 (pale violet red, PDB ID: 7ZWI)). Clashes are indicated with yellow stars. **(E)** Blue Native PAGE-based mobility shift assay of FLAG-tag purified Pfs230:Pfs48/45 complex, incubated with a tenfold molar excess of individual anti-Pfs230 or anti-Pfs48/45 Fab fragments. The first lane shows the Pfs230:Pfs48/45 complex with the addition of a mock liquid (PBS) as a reference, and PGDM1400, an anti-HIV envelope protein targeted Fab, was included as negative control. Fabs are sorted by antigen, epitope, and potency (underlined: IC_80_ < 10 µg/ml; bold: IC_80_ = 10-100 µg/ml, others: IC_80_ > 100 µg/ml; See also **Table S1**). Note that due to the nature of Blue Native PAGE, the hydrodynamic radius and therefore the angle of approach can heavily influence the extent of Fab-induced mobility retardation. The symbols #, *, ×, and + denote Pfs230, Pfs230:Pfs48/45, Pfs230:Fab, and Pfs230:Pfs48/45:Fab, respectively. **(F)** Infection of *An. stephensi* mosquitoes with wildtype NF54 (white), Pfs230^ΔD13/14^ (red), and Pfs230^ΔCterm^ (pink) as measured in five independent standard membrane feeding experiments with 20 mosquitoes per condition each. Dots represent the number of oocysts per midgut of an individual mosquito. Note that data from experiment 4 are the same as the FCS controls in Figure S8H experiment 2. N denotes the number of mosquitoes that carries at least 1 oocyst, mean is the mean number of oocyst per midgut. **(G)** Transmission reducing activity of 10 µg/ml Pfs230D4-targeting mAb 2A2 for mosquito infections with NF54 wildtype (white) and Pfs230ΔCterm (pink) gametocytes, in the presence of active human complement. Mean with 95% confidence intervals are shown. Statistical analysis was done using a one-tailed t-test to determine whether TRA was higher than 0%: ns = not significant, ****=p<0.0001. *See also Figure S8, Table S1, and S3-S4*.

### Pfs48/45 epitope accessibility in the Pfs230:Pfs48/45 complex

Next, we investigated how the Pfs230 interaction with Pfs48/45 might affect the accessibility of Pfs48/45-targeting antibodies. An overlay of structures of previously characterized antibodies with the Pfs230-bound structure revealed that Pfs48/45D1, Pfs48/45D2 and Pfs48/45D3-1b epitope antibodies approach from the side of the Pfs48/45 disc and are accessible when Pfs230 is bound **(Figure S8C-E)**. In contrast, many antibodies that target the Pfs48/45D3-1a epitope, including highly potent antibodies TB31F and RUPA-29, would have considerable clashes with both Pfs230D13 and Pfs230D14 in this conformation **(Figure 4C)**.

To determine whether these antibodies can bind to Pfs48/45 in the context of the Pfs230:Pfs48/45 complex, we used the BN-PAGE-based mobility shift assay to study the changes in migratory pattern of Fab-bound endogenous Pfs230:Pfs48/45 complex. To this end, the purified complex was incubated with a tenfold molar excess Fab derived from a range of both potent and non-potent TRA antibodies, spanning almost all currently known epitopes on the complex. As expected, α-Pfs230 Fabs shifted both bands upwards, with the exception of 15C5, which targets epitope III on Pfs230D1 that is shielded by Pfs230D2 and thus incompatible with binding full-length Pfs230 (**Figure 4E**) (17, 20). None of the other tested α-Pfs230 Fabs, targeting epitopes on Pfs230D1, Pfs230D4 and Pfs230D7, disrupted the Pfs230:Pfs48/45 complex, in agreement with the relative orientation of Pfs230 on Pfs48/45 in our structure. The α-Pfs48/45 Fabs did not shift the lower free Pfs230 band, but did affect migration of the upper band. While some Fabs i) bound to the intact complex (*e.g.* RUPA-44, RUPA-117), others ii) disrupted the Pfs230:Pfs48/45 complex (*e.g.* RUPA-58 and RUPA-50), or iii) showed a heterogeneous effect resulting in Fab-bound and unliganded Pfs230:Pfs48/45 (*e.g.* TB31F, RUPA-47) (**Figure 4E**). The non-potent 10D8, targeting Pfs48/45-D2, could not bind to the Pfs230:Pfs48/45 complex. To confirm the observation that some but not all Pfs48/45-targeted antibodies could disrupt the Pfs230:Pfs48/45 complex, we tested whether a subset of these Fabs could displace Pfs230 on the female gamete surface in a competition assay. In agreement with the BN-PAGE results, RUPA-50, RUPA-57, and RUPA-58 significantly decreased Pfs230 on the gamete surface in a dose-dependent manner, while *e.g.* TB31F, RUPA-46, and RUPA-89 Fabs showed no displacement of Pfs230 (**Figure S8F**).

Intriguingly, TB31F could still partially bind to the Pfs230:Pfs48/45 complex in BN-PAGE (**Figure 4E**) and did not displace Pfs230 on the gamete surface (**Figure S8F**), despite modeling based on our Pfs230-bound Pfs48/45 structure suggesting that significant steric clashes would occur. In this context, it is interesting to note again that it has previously been described how Pfs48/45 can adopt a disc-like conformation (similar to the structure reported here in complex with Pfs230), but also more extended conformations where Pfs48/45D3 can reorient from the rest of the protein. Given that Pfs48/45D1-2 contributes 1169 Å^2^ of a total 1647 Å^2^ buried surface area within the Pfs48/45-Pfs230 interface, it is conceivable that Pfs48/45D1-2 is sufficient for Pfs230 binding, allowing for Pfs48/45D3 to retain a range of motion in the complex. Modeling Pfs48/45D3 antibodies onto the previously determined extended conformation of Pfs48/45 with Pfs230D13-14 aligned to its Pfs48/45D1-2 binding site demonstrates that Pfs48/45D3-1a epitope would be accessible in this arrangement (**Figure 4D**). This elongated conformation can also accommodate binding for most anti-Pfs48/45 antibodies that target epitope 1b or Pfs48/45D1-2, with the exception of 10D8 (**Figure 4D, Figure S8I**) (14, 15). Together, this could indicate that Pfs230-bound Pfs48/45 retains flexibility, and that this mobility may allow for the binding of antibodies that would otherwise be blocked in the Pfs230-bound disc-like conformation of Pfs48/45.

### Pfs230 membrane retention is non-essential for transmission

Interestingly, the ability of Fab fragments to induce (partial) disruption of the Pfs230:Pfs48/45 complex does not seem to be associated with TRA of the corresponding antibody in SMFAs (**Figure 4E, Figure S8F, Table S1**). For example, when looking at Pfs48/45D3-1a targeting antibodies, the extremely potent antibody TB31F (IC_80_ < 2 µg/ml) shows only partial binding *in vitro* and no displacement of Pfs230 *in vivo*. More importantly, the non-potent RUPA-57 (IC_80_ > 100 µg/ml) is able to fully dissociate the complex *in vitro* and partially on the gamete surface, mimicking the phenotype of *e.g.* the potent RUPA-50 (IC_80_ < 2 µg/ml). The binding characteristics for RUPA-57 suggest that the capability for a mAb to induce Pfs230:Pfs48/45 complex dissociation does correlate with potent transmission-reducing activity.

To investigate whether Pfs230:Pfs48/45 complex disruption could still be a mode of action for some of the transmission-blocking antibodies, we further characterized Pfs230 truncation parasite lines. These parasite lines mimic the most severe outcome of antibody-induced complex dissociation, where all Pfs230 is dissociated from the parasite plasma membrane. Surprisingly, both Pfs230^ΔD13/14^ and Pfs230^ΔCterm^ gametocytes showed high oocyst intensities in multiple independent membrane feeding experiments using *Anopheles stephensi* mosquitoes (**Figure 4F**), in contrast to almost complete lack of oocysts reported for Pfs230^KO^ parasite lines (7). Although our truncation lines showed lower oocyst intensity than wildtype parasites in some experiments, other experiments showed no difference. We confirmed by midgut oocyst genomic DNA extraction and PCR that the oocysts formed by the truncation lines only contained mutant parasite DNA, ruling out accidental wildtype contaminations (**Figure S8G**). To further confirm that Pfs230 remains dissociated from the parasite surface within the mosquito midgut, we added the Pfs230D4-targeting antibody 2A2 into the bloodmeal with either wildtype NF54 and Pfs230^ΔCterm^ gametocytes. While 2A2 almost completely blocks transmission for wildtype parasites at 10 µg/ml (TR 99.1% (95% CI: 98.5 to 99.4)), we observed no significant TRA for 10 µg/ml 2A2 with the Pfs230^ΔCterm^ truncation gametocytes (TRA 13.0% (95% CI:-5.7 to 28.4)) (**Figure 4G, Figure S8H**). Together, these data indicate that Pfs230:Pfs48/45 complex disruption is not the mode of action for functional transmission-blocking α-Pfs48/45 (or α-Pfs230) antibodies.

### Delineation of Pfs230D4 and D7 epitopes

Our cryo-EM studies reveal potent epitopes outside of Pfs230D1 that had not been previously structurally delineated. Fab 2A2 could be resolved in the Pfs230D1-6 map, while Fab 18F25 could be resolved in the Pfs230D7-8 map (**Figure 1I**, **Table 1**). X-ray crystal structures of unliganded 2A2 and 18F25 Fabs were solved to 1.2 Å and 1.6 Å resolution (**Table S3**), respectively, and docked and refined into the maps. Our cryo-EM structure reveals that 2A2 targets Pfs230D4 (**Figure 5A**), confirming the epitope location previously proposed based on the differential binding and TRA of mAb 2A2 against *P. falciparum* strains that contain SNPs in Pfs230D4 (38). 2A2 buries 824 Å^2^ on Pfs230D4, primarily making contacts through the KCDR3 and all HCDR loops (**Figure 5A**, **Table S4**). Most residues contacted by the antibody lie in the canonical perpendicular β-hairpin of Pfs230D4 (β5-β6) and surrounding regions (β2-β4 regions) (**Table S4**). A moderate number of additional contacts are facilitated by the KCDR1/3 within the loop region between the β10 and β11 strands (**Table S4**). Analysis of the MalariaGEN Pf7 database reveals that many SNPs differing from the NF54 strain are found in the 2A2 epitope in various parasite strains (**Figure 5B**). Some of the SNPs are quite rare with frequencies less than 1% (Q1196E, E1198V, E1198D, L1208F, N1209K and N1209Y, Q1250P, Q1250L, Q1255E, K1259I, and K1261N), while others are seemingly more common (H1159D – 53.3%, Y1194S – 59.3%, N1209H – 19.2%, Q1250K – 86.3%, and Q1250E – 12.6%).

**Figure 5:**
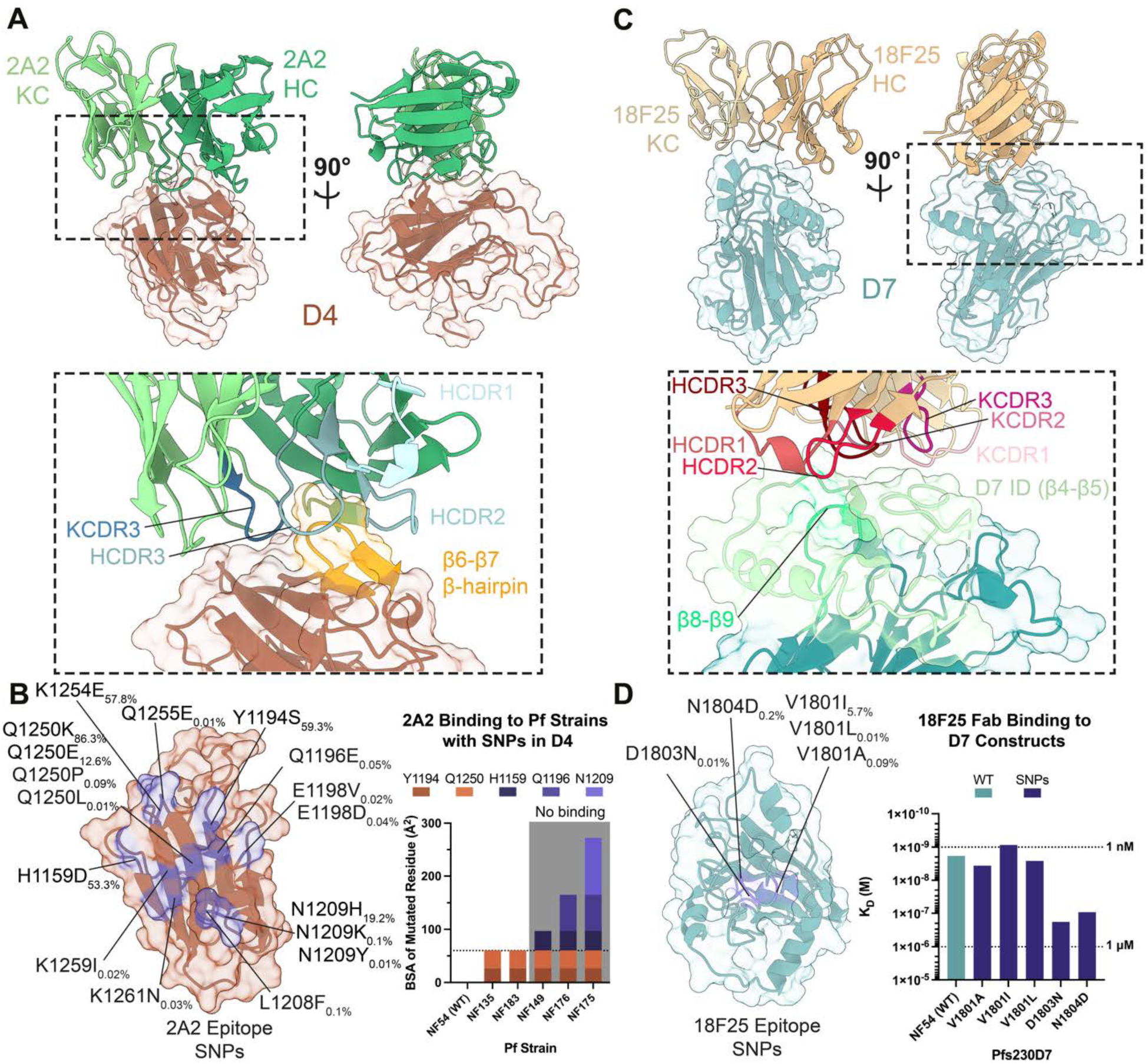
Structural delineation of potent Pfs230 epitopes outside of Pfs230D1. **(A)** The variable region of 2A2 (green) and Pfs230D4 (orange) are shown with key contacts (inset) made by the HCDR1, HCDR2, HCDR3, and KCDR3 (shades of blue) of mAb 2A2 with the β6-β7 perpendicular β-hairpin (yellow) of Pfs230D4 highlighted. **(B)** SNPs of Pfs230D4-2A2 epitope identified in the MalariaGEN Pf7 databse database (53) are indicated in purple and labeled. The frequency of the SNP is shown as a subscript. (right) Five strains of *P. falciparum* harbouring different combinations of five different SNPs (x-axis) with the BSA that that residue contributes to binding in the structural model (y-axis). Specific amino acid residue BSA contributions are indicated by shades of different colours. Residues found exclusively in *P. falciparum* strains that have perturbed ability to bind female gametes in surface immunofluorescence assays (asterisks) are coloured in shades of purple and appear above the dotted line. All other residues are coloured in shades of orange (**C)** The variable region of 18F25 (tan) and Pfs230D7 (cyan) are shown. 18F25 makes contacts with the Pfs230D7-ID (light green) and the β8-β9 loop (neon green) using all HCDRs (shades of red) and KCDRs (shades of pink) of 18F25 with Pfs230D7 (inset). **(D)** SNPs of Pfs230D7-18F25 epitope identified in the MalariaGEN Pf7 database (53) are indicated in purple and labeled. The frequency of the SNP is shown as a subscript. Summary of surface plasmon resonance experiments utilizing 18F25 Fab and Pfs230D7 variants (y-axis) (WT, teal; and SNP mutants, purple) and their determined affinity (y-axis). Dotted lines indicate K_D_ values of 1 nM and 1 μM. *See also Tables S3-4*.

A subset (H1159D, Y1194S, Q1196E, N1209Y, Q1250K) of these SNPs are found in strains of *P. falciparum* in which 2A2 had diminished TRA and perturbed binding to live female gametes in surface immunofluorescence assays (SIFA) (**Figure 5B**) (38). Two mutations (Y1194S and Q1250K) are both present in the NF135 and NF183 strains but are not sufficient to perturb binding of 2A2 **(Figure 5B)**. In contrast, the NF149, NF175, and NF176 strains also encode these mutations, and they also carry 1-3 additional SNPs (H1159D, Q1196E, and N1209Y) found in the 2A2 epitope **(Figure 5B)**. Live gametes obtained from these parasite lines are unable to bind mAb 2A2 in SIFAs (38).

The epitope of mAb 18F25 localizes to Pfs230D7 (**Figure 5C**) (37). Integrative modeling in the global 3.8 Å global resolution map suggests that 18F25 buries 515 Å^2^ on Pfs230D7 through all CDRs of the antibody (**Table S4**). 18F25 primarily engages with the Pfs230D7-ID found in between β4 and β5, with a small amount of interactions localized to the β8-β9 loop (**Figure 5C, Table S4**). Unlike the 2A2 epitope and Pfs230D4 overall, both the 18F25 epitope and Pfs230D7 are more conserved (**Figure 5D**). Only two SNPs in Pfs230D7 identified from the MalariaGEN Pf7 database have frequencies higher than 10% (Y1829N – 69.4% and I1870V – 13.5%) and these are not contained within the 18F25 epitope (53). Only three residues within the epitope are reported with mutations in the database. V1801I, V1801L, and V1801A are conservative mutations that have a combined frequency of 5.8% (5.7%, 0.01%, and 0.09%, respectively). Two other SNPs of D1803N and N1804D are also conservative mutations that occur relatively infrequently (D1803N – 0.01% and N1804D – 0.2%).

To better understand how the SNPs found in the 18F25 epitope affect the affinity of the mAb to Pfs230D7, we performed surface plasmon resonance (SPR) experiments (**Figure 5D**) using Pfs230D7 constructs expressed using a wheat germ cell-free protein expression system (54). SPR experiments utilizing the 18F25 Fab and WT Pfs230D7 revealed that the antibody binds to its native epitope with single digit nanomolar affinity (K_D_ = 1.8 nM). Mutation of the most polymorphic residue, V1801, to the three different SNPs identified in the MalariaGEN Pf7 database revealed a negligible effect on binding affinity, with Fab 18F25 maintaining wild-type level nanomolar binding affinity (V1801A: K_D_ = 3.6 nM, V1801L: K_D_ = 2.6 nM and V1801I: 0.9 nM, respectively). While the rarer SNPs D1803N and N1804D have a more moderate effect (K_D_ = 181 nM and K_D_ = 92 nM, respectively), nanomolar-level binding affinity is still maintained in these cases. Together, our data elucidate the 2A2 and 18F25 epitopes on Pfs230D4 and D7, respectively, providing structural insights into Pfs230-directed antibody responses outside of Pfs230D1. In contrast to 2A2, mAb 18F25 targets a conserved epitope with rare naturally occurring non-synonymous SNPs that have small impact on 18F25 binding affinity.

### Potent Pfs230 antibodies target membrane distal epitopes

mAb 2A2, while showing strain-dependent TRA, was highly potent in SMFAs utilizing the NF54 strain (IC_80_ = 1.9 μg/ml (38)). The potency was on par with the highly potent RUPA-97 that targets Pfs230D1 (IC_80_ = 0.7 μg/mL (20)), suggesting that a common feature of these mAbs might be responsible for the high potency. These two mAbs, despite binding different domains, bind to the same side of Pfs230 with a similar angle of approach (**Figure 6A**). mAb 18F25 (IC_80_ < 30 μg/ml (37)) also binds to the same face of Pfs230, while differing slightly in its angle of approach (**Figure 6A**). Interestingly, Pfs230D1-directed mAbs of lower potency, such as RUPA-38 (IC_80_ > 100 μg/mL) (20), bind to the opposite face of where RUPA-97, 2A2, and 18F25 bind to Pfs230 (**Figure 6A**). This trend is generally applicable to α-Pfs230D1 mAbs published to date (**Figure S9**) (17).

**Figure 6:**
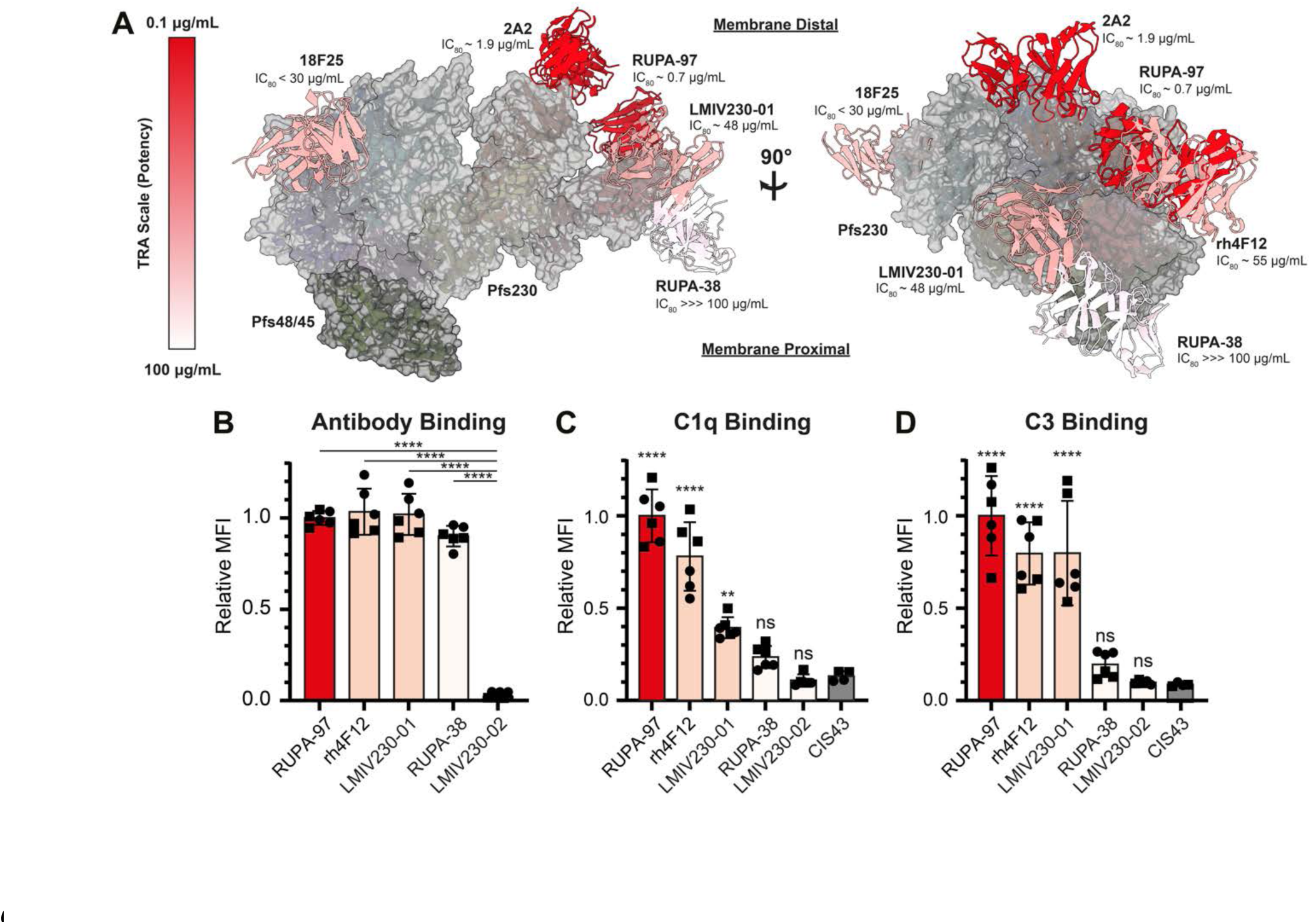
Molecular basis of Pfs230-directed mAb potency. **(A)** The binding sites of 18F25, LMIV230-01, 2A2, RUPA-97, RUPA-38, and rh4F12 shown in the context of Pfs230 (surface: gray, ribbons: domains coloured as done in Figure 1) and Pfs48/45 (surface: black, ribbons: domains coloured as done in Figure 1). The antibodies are coloured in accordance with their IC_80_ values or TRA via heat map with darker colours indicating higher TRA or potency (left scale). The membrane distal side corresponds to the top of the panel and the membrane proximal side at the bottom of the panel. **(B-D)** Anti-Pfs230D1 mAb binding (B), or antibody-mediated human C1q recruitment (C) and human C3 deposition (D) at 1 µg/ml on the surface of wildtype NF54 macrogametes as determined by flow cytometry. CIS43, an anti-PfCSP antibody, was included as a negative control for the C1q and C3 deposition assays. Mean fluorescence intensity (MFI) values were normalized against the mean of RUPA-97 replicates, to allow averaging across two independent experiments (depicted as different symbols) with two to three technical replicates each. Mean with standard deviations are shown. Statistical analysis was done using an ordinary one-way ANOVA with either a Tukey’s multiple comparisons test with a single pooled variance comparing all groups to each other (panel B, only showing significant differences) or Dunnett’s multiple comparisons test with a single pooled variance comparing all groups to CIS43 (C-D). ns = not significant; **=p<0.01, ****=p<0.0001. *See also Figure S9 and Table S1*.

These Pfs230-targeting mAbs have been demonstrated to be complement-dependent for their TRA (20, 36–38, 40). Previously, it has been shown that for efficient C1q deposition on an antigenic surface, the first step in the classical complement activation pathway, IgGs need to form into ordered hexameric antibody structures (55, 56). Intriguingly, the epitopes of potent mAbs RUPA-97, 2A2, and 18F25 are localized to the opposite face of where Pfs48/45 and presumably the PPM would be (**Figure 6A**). In contrast, the epitope of non-potent RUPA-38 is localized near the presumed location of the PPM (**Figure 3A**). These observations led us to hypothesize that the angle of approach of the antibody is a determinant of Pfs230-directed antibody potency, as antibodies that are near the PPM might be prevented from efficient recruitment of complement components and thus complement activation. To test this hypothesis, we selected five different mAbs targeting Pfs230D1 with different potencies (RUPA-97 (IC_80_ = 0.7 μg/mL), rh4F12 (IC_80_ = 55 μg/mL), LMIV230-01 (IC_80_ = 48 μg/mL), RUPA-38 (IC_80_ > 100 μg/mL)), and LMIV230-02 (IC_80_ > 1000 μg/mL) (19, 20), expressed these as IgGs of the same complement-fixing subclass, and tested antibody binding, C1q recruitment and C3 deposition on live female NF54 gametes. All antibodies except LMIV230-02 bound gametes with similar intensity (**Figure 6B**), which is in agreement with previously determined *in vitro* affinities (20). In contrast, the highly potent antibody RUPA-97 and moderately potent antibodies rh4F12 and LMIV230-01 were able to recruit C1q and deposit C3 on the surface, while the non-potent RUPA-38 was not able to do so efficiently (**Figure 6C-D**). These findings, as implied by the Pfs230:Pfs48/45 complex structure, suggest that the angle of approach relative to the PPM might be a critical determinant for potency in complement-dependent Pfs230-directed antibodies.

## DISCUSSION

Targeting human-to-mosquito transmission, a developmental bottleneck in the *Plasmodium* life cycle, is a promising intervention strategy for malaria control and elimination. Here, we reveal the structural disposition of the heterodimer complex Pfs230:Pfs48/45, which is essential for *P. falciparum* transmission and a major target for malaria transmission-blocking vaccine development. Using advances in molecular parasitology and cryo-EM, we were able to circumvent recombinant expression by purifying the approximately 410 kDa large, membrane-bound Pfs230:Pfs48/45 complex from its native source, late-stage *P. falciparum* gametocytes. Previously, similar methods have been used to elucidate the structure of the *Plasmodium* Translocon of Exported Proteins (PTEX) complex (57) and the soluble RhopH complex (58, 59), both crucial protein complexes within the *Plasmodium* life cycle non-amenable to standard recombinant protein production. Using genome engineering to enable the purification of endogenous proteins for cryo-EM studies has also found traction in research on other organisms, including human cells (60), and bacterial pathogens like *Pseudomonas spp.* (61) and *Mycobacteria spp.* (62). This highlights the opportunities that this approach, combined with recent advances in the field of structural biology including bottom-up structural proteomic platforms such as cryoID (63) and protein folding prediction software such as AlphaFold 3 (64), may offer to the many other *Plasmodium* proteins refractory to recombinant expression.

Our structure reveals how insertion domains present within Pfs230 6-Cys domains mediate inter-domain interactions **(Figure 2)** as seen before with the Pf12:Pf41 heterodimer complex, which are two other malarial proteins that are composed of 6-Cys domains (50, 51). Additionally, inter-protein contacts between Pfs230 and Pfs48/45 appear to be mediated by regions outside of the canonical 6-Cys fold; we have shown how the C-terminal extension of Pfs230 is critical for binding, and observed that the N-terminal extension of Pfs48/45 also localized to this interface **(Figure 4B-C, Figure S6A)**. This suggests that regions outside of the canonical 6-Cys fold may have been introduced to provide functional roles by mediating interactions, which could be a generalizable property of 6-Cys domain family of proteins.

Although it is currently unknown what exact cellular process the Pfs230:Pfs48/45 complex is involved in, knockout studies in *P. falciparum* and the rodent malaria parasite *Plasmodium berghei* suggest the Pfs230:Pfs48/45 complex is involved in microgamete-to-macrogamete adhesion and microgamete-to-RBC adhesion resulting in exflagellation centers (7, 8, 65). The presented structure will enable the design of domain clusters with more precision than previously possible for the generation of genetically modified parasite lines that can interrogate Pfs230 function. Our structure also shows that the N-terminal domains are membrane-distal, potentially allowing interactions with other elements. This is in line with previous descriptions of some functional complement-independent α-Pfs230D1 antibodies, suggesting that Pfs230D1 might be involved in some protein interactions that are disrupted upon binding of these antibodies (16, 34). Furthermore, competition assays with recombinant Pfs230Pro-D1-2 was previously shown to inhibit the formation of exflagellation centers (16), further suggesting that the N-terminus of Pfs230 might be involved in cell-to-cell adhesion. The C-terminal Pfs230 domains are involved in the Pfs48/45-based membrane anchoring onto the PPM. We showed that functional activity of complement-independent Pfs48/45 antibodies does not correlate with their ability to disrupt the Pfs230:Pfs48/45 complex. The transgenic parasite lines expressing truncated Pfs230, which lose Pfs230 membrane-retention but are able to efficiently infect *Anopheles* mosquitoes, fully support the notion that complex-dissociation is not a mode of action for transmission-blocking antibodies. These observations imply that Pfs48/45 does not merely serve as an anchor for Pfs230 membrane retention, but carries out an additional critical function during parasite transmission that anti-Pfs48/45 antibodies can potently inhibit.

Surprisingly, there is a stark difference between the truncation parasite lines Pfs230^ΔD13/14^ and Pfs230^ΔCterm^ on one hand that do infect mosquitoes, and the previously reported Pfs230^KO^ parasite lines on the other hand that almost completely lost capability for parasite transmission (7). Future studies should focus on uncovering the roles of individual domains of Pfs230 and Pfs48/45 during parasite transmission.

Next to characterizing the full-length structure of the Pfs230:Pfs48/45 heterodimer, we also molecularly characterized the epitope of multiple transmission-blocking antibodies. The potent Pfs48/45D1 antibody, RUPA-71, binds an adjacent epitope to the previously characterized mAb, RUPA-58 (*15*). All highly potent Pfs48/45D1 antibodies compete with either RUPA-58 or RUPA-71, indicating that binding to this conserved portion of Pfs48/45D1 is associated with transmission-reducing activity (*15*). Additionally, we structurally characterized potent epitopes outside of Pfs230D1 found on Pfs230D4 and Pfs230D7 targeted by murine mAbs, 2A2 and 18F25, respectively. While mAb 2A2 is notably potent when used against the NF54 strain, the epitope is highly polymorphic with many Pfs230D4 SNPs observed in field isolates (38) and described in the MalariaGEN Pf7 database (53). In contrast, both Pfs230D7 and the 18F25 epitope are highly conserved and may represent a yet unexplored opportunity for subunit vaccine development. Excitingly, the few naturally occurring SNPs found in the 18F25 epitope were non-detrimental for antibody binding *in vitro* **(Figure 5D)**. However, in comparison to 2A2 and highly potent Pfs230D1-bin I binders, 18F25 exhibits approximately 10-fold lower potency. The ability to design Pfs230 probes based on the full-length structure will now enable discovery efforts seeking more potent antibodies. In addition, the structure of the full Pfs230:Pfs48/45 heterodimer complex reported here with structure-function relationships of antibody inhibition, now enables a more fulsome opportunity for structure-guided immunogen design, which has been transformative in vaccinology during the development of *e.g.* SARS-CoV-2, and respiratory syncytial virus vaccines (66, 67).

Immunogens based on Pfs230 and Pfs48/45 have already shown promising results in clinical trials (44–47), though the induced inhibitory activity was incomplete and the durability of the response remains uncertain. Our findings suggest that current TBV candidates can be further optimized, for example by including Pfs230 domains that were previously thought to be unable to induce functional transmission-blocking antibodies (36–38), or by rational structure-based immunogen design (68, 69). To date, most Pfs48/45-based interventions focus on Pfs48/45D3, but Pfs48/45D1 epitopes may be beneficial inclusions into future Pfs48/45-based immunogens. Currently, in the case of Pfs230 domains, Pfs230D1, D4, D7, and D12 have been demonstrated to elicit potent antibodies (20, 36–38). Interestingly, these Pfs230 domains share a common feature in that they are membrane distal, suggesting that domains present on this side should be prioritized for subunit vaccine development. By this logic, Pfs230D8 and D9 may also be relevant targets (**Figure 6**). Future immunogen design efforts should integrate inter-domain structural elements to present well-folded epitopes. Approaching immunogen design through stabilizing structural clusters, guided by the structure reported here, instead of individual domains will help eliminate the presentation of non-neutralizing epitopes that are absent endogenously, as previously evidenced by the elicitation of Pfs230D1-binIII mAbs (15C5, LMIV230-02, and 230AS-26) and 230AL-20 by a Pfs230D1-only antigen (**Figure S9**). The presented Pfs230:Pfs48/45 structure might also allow for future protein engineering campaigns that graft multiple potent epitopes onto a single chimeric immunogen (70), or use stabilizing mutations in conjunction with display on nanocages (71). Additionally, advances in mRNA vaccination approaches combined with precise immunogen design might allow for the *in situ* expression of stable, native Pfs230:Pfs48/45 co-assemblies. Ultimately, the aim will be to create the most potent next-generation TBVs possible to aid in meeting the ambitious goals of malaria elimination and eradication.

### Limitations of the study

A limitation of our structure is the low resolution obtained for the Pfs230D9-12 domains (global resolution of 4.7 Å). This prevented building an atomic model for these domains, limiting the information that could be confidently extracted from this part of the map. Instead, we combined this low-resolution map with an AF-based modeling approach to obtain a backbone model with reasonable confidence regarding the domain orientations in this region of Pfs230. Additionally, while the presence of 2A2 does not seem to affect the overall structure of Pfs230:Pfs48/45 significantly **(Figure S3A-B)**, other antibodies in our structure could have potentially altered the structure of Pfs230:Pfs48/45 compared to its unliganded state. An unliganded Pfs230:Pfs48/45 heterodimer structure reported by others in parallel to our study described a similar overall structure for the heterodimer complex, indicating that antibody binding in our structure did not have a dramatic effect (72). One notable difference in the 6Fab-bound Pfs230:Pfs48/45 structure compared to the unliganded structure is the increased flexibility within the Pfs230D9-14 region, specifically between Pfs230D12 and D13. In 2D classes of the full complex, the constant region of anti-Pfs48/45 Fab, RUPA-71, appears to be in proximity to the Pfs230D1-6 and Pfs230D7-8 clusters of the molecule, potentially causing steric hinderance. Additionally, loops at the periphery of Pfs48/45D1 which come into close proximity of Pfs230D13-14 is not localized to the same region in the antibody-bound structure, pushing Pfs48/45 away from Pfs230. We propose that this steric hindrance induced by RUPA-71 binding decompresses the Pfs230D9-12 cluster, resulting in a more dynamic disposition, and consequently, poorer resolution of this portion of the map. Finally, we purified the Pfs230:Pfs48/45 from late-stage gametocytes instead of gametes, for practical reasons. However, during gamete formation the Pro-domain of Pfs230 is proteolytically cleaved (73), and therefore our structure does not fully recapitulate the mature protein species on gamete surfaces. How these additional Pro-domain residues add onto the structure and if these influence the overall Pfs230:Pfs48/45 conformation, remains currently unknown.

## RESOURCE AVAILABILITY

### Lead contact

Further information and requests for resources and reagents should be directed to and will be fulfilled by the lead contact, Jean-Philippe Julien upon reasonable request.

## Materials availability

All unique reagents generated in this study are available from the lead contact with a completed Materials Transfer Agreement.

## Data and code availability

- Previously unpublished antibody sequences are available in Table S6. Crystal and cryo-EM structures have been deposited in the Protein Data Bank and are publicly available as the date of publication. EMDB and PDB IDs are listed in Table 1 and Table S3. The proteomics dataset has been deposited in the ProteomeXchange Consortium via the PRIDE partner repository (74) with the dataset identifier PXD060716.
- This paper does not report original code.
- Any additional information required to reanalyze the data reported in this paper is available from the lead contact upon reasonable request.

## ACKNOWLEDGEMENTS

We thank Roos de Jong and Sanne Grievink for assistance with protein purification; Laura Pelser, Astrid Pouwelsen, Jacqueline Kuhnen, Jolanda Klaassen, and Wouter Graumans for mosquito rearing and dissections; April Yu for assisting in 18F25 Fab structural determination; Hikaru Nagaoka and Takafumi Tsuboi for their assistance and guidance in Pfs230D7-18F25 binding characterization; Dick Zijlmans for assistance with Mass spectrometry data analysis; Aran Labrijn (Genmab) for antibody sequencing; the Raboudumc Technology Centers for Microscopy and Flow Cytometry for access to their instruments; Samir Benlekbir and Zhijie Li from the SickKids Nanoscale Biomedical Imaging Facility for assistance and insights during cryo-EM data collection; Greg Wasney and James Magnus Jorgensen at the Structural & Biophysical Core (SBC) Facility; past and present members of the Molecular and Cellular Parasitology (Radboudumc) and the Malaria Transmission (Radboudumc) teams for insightful discussions. This work was supported by a Radboudumc Master-PhD grant awarded to E.T.B. and a VIDI grant from the Netherlands Organisation for Scientific Research to M.M.J. (fellowship number 192.061). This work was also undertaken, in part, thanks to funding from a Canadian Institutes for Health Research grant (428410) and was supported by the Canada Research Chair program awarded to J.L.R. and J-P.J. E.T.B. was further supported by a FEMS Research and Training Grant. The Pfs230D7 production and SPR analysis were performed with support from AMED (JP24wm0225045).R.Y. was supported by a Canada Graduate Scholarship - Master’s (CGS-M). S.H. was supported by a Canada Graduate Scholarship - Doctoral. C.M.A. is supported by a VENI grant from the Netherlands Organisation for Scientific Research (VI.Veni.222.381). The Vermeulen lab is part of the Oncode Institute, which is partly funded by the Dutch Cancer Society (KWF). X-ray diffraction experiments for the 2A2 and 18F25 Fab datasets were performed at the AMX-17-ID-1 at the National Synchrotron Light Source II, a U.S. Department of Energy (DOE) Office of Science User Facility operated for the DOE Office of Science by Brookhaven National Laboratory under Contract No. DE-SC0012704. X-ray diffraction experiments for RUPA-71 Fab was performed using beamline CMCF-ID at the Canadian Light Source, a national research facility of the University of Saskatchewan, which is supported by the Canada Foundation for Innovation (CFI), the Natural Sciences and Engineering Research Council (NSERC), the National Research Council (NRC), the Canadian Institutes of Health Research (CIHR), the Government of Saskatchewan, and the University of Saskatchewan. Molecular graphics were generated using UCSF ChimeraX, developed by the Resource for Biocomputing, Visualization, and Informatics (University of California, San Francisco) with support from the National Institutes of Health (R01-GM129325) and the Office of Cyber Infrastructure and Computational Biology, National Institute of Allergy and Infectious Diseases. The size exclusion chromatography instrument was accessed at the Structural and Biophysical Core Facility, The Hospital for Sick Children, and EM data was collected at the Nanoscale Biomedical Imaging Facility, The Hospital for Sick Children, supported by the Canada Foundation for Innovation and Ontario Research Fund.

## AUTHORS CONTRIBUTION

**Conceptualization**: E.T.B., R.Y., S.H., M.M.J., J-P.J.; **Investigation**: E.T.B., R.Y., S.H., N.I.P., R.S., G-J.v.G., A.M., T.Y., O.T.W., R.C.v.D., M.R.I., C.M.A., P.W.T.C.; **Formal Analysis**: E.T.B., R.Y., S.H., F.H, D.I., M.J.; **Resources**: M.V., T.B., J.L.R.; **Writing – Original Draft**: E.T.B., R.Y., S.H., M.M.J., J-P.J.; **Writing – Review & Editing**: E.T.B., R.Y., S.H., F.H., D.I., M.J., N.I.P., C.M.A., T.B., J.L.R., T.W.A.K., M.M.J., J-P.J.; **Supervision**: M.V., T.B., E.T., J.L.R., T.W.A.K., M.M.J., J-P.J.; **Funding acquisition:** E.T.B., M.M.J., J-P.J.

## METHODS

### *Plasmodium falciparum* genome modification plasmids

To create the Cas9 guide plasmids, a pair of complementary guide-encoding oligonucleotides 5’-TTAATGATGGCTCTTGATTG-3’ (targeting Pfs230 C-terminus to create iGP2^230-tag^ / 230^ΔD13/D14^ / 230^ΔCterm^) and 5’-ATTCTATTACATTATCAAGA-3’ (targeting Pfs230D13 to create 230^ΔD13/14^) were treated with T4 polynucleotide kinase, annealed, and ligated into BbsI-digested pMLB626 (a kind gift from Marcus Lee) (76).

To generate the iGP2^230-tag^ homology-directed repair (HDR) template, partial 5’ and full 3’ homology regions (HR) were PCR amplified from NF54 genomic DNA (**Table S5**). Synonymous mutations to shield the Cas9 guide region in the full 5’HR were inserted by overlap PCR using the partial 5’HR and two annealed custom-made oligonucleotides (Sigma-Aldrich, **Table S5**). A selection cassette containing the i) PfH2B promoter, ii) the *mScarlet* gene, and iii) a bidirectional 3’UTR (PBANKA_142660) was PCR amplified from an in-house plasmid (77) as two separate amplicons to remove internal restriction sites. These two amplicons were cloned into pGGASelect using a BsaI-HFv2 Golden Gate reaction, and further sub-cloned using AatII and BamHI-HF into a pUC19-based plasmid that had all internal type IIS restriction enzymes removed via site-directed mutagenesis, yielding pRF0508. The final HDR template was created using the NEBridge Golden Gate Assembly Kit (BsmBI-v2), combining i) pGGAselect, ii) the 5’ HR amplicon, iii) two annealed oligonucleotides coding for a tandem affinity purification tag (GTSG-(3xFLAG)-GSG-EPEA-stop) (**Table S5**), iv) pRF0508, and v) the 3’ HR amplicon.

To generate HDR templates to make Pfs230^ΔD13/14^ and Pfs230^ΔCterm^, the respective 5’HRs (Pfs230^ΔD13/D14^ spanning Pfs230D11-12, Pfs230^ΔCterm^ spanning Pfs230D13-14) and 3’HR were PCR amplified from NF54 genomic DNA (**Table S5**). The selection cassette was modified by swapping the PfH2B promoter in pRF0508 for the PfGAPDH promoter (Pf3D7_1462800). The HDR templates were created in a golden gate reaction combining i) pGGAselect, ii) respective 5’HR amplicon, iii) PCR amplicon coding for (GGSG-3xFLAG-stop), iv) GAPDH selection marker, and v) the 3’ HR amplicon.

All PCR reactions were performed using Primestar® GXL DNA polymerase (Takara Bio), all other cloning enzymes were obtained via New England Biolabs. Sequences were confirmed by Sanger Sequencing (Baseclear).

### *Plasmodium falciparum* culturing, transfections, and gametocyte production

*P. falciparum* cultures were maintained at 37 °C with 3% O_2_ and 4% CO_2_, in complete medium (RPMI1640 medium with 25 mM HEPES, 25 mM NaHCO_3_, 10% human type A serum) with 5% O^+^ human red blood cells (Sanquin, The Netherlands) (78). Asexual NF54/iGP2 and NF54/iGP2^230-tag^ parasites were cultured in the presence of 2.5 mM D-(+)-glucosamine hydrochloride (Sigma #1514). Parasites were synchronized using sorbitol treatment before transfection or gametocyte induction (79).

For transfection, 80 µg of HDR plasmid was linearized overnight, ethanol precipitated and co-transfected with 80 µg of Cas9 guide plasmid by ring-stage transfection as described previously (80, 81). Briefly, plasmids were resuspended in cytomix (10 mM K_2_HPO_4_/KH_2_PO_4_ pH 7.6, 120 mM KCl, 0.15 mM CaCl_2_, 5 mM MgCl_2_, 25 mM HEPES, 2 mM EDTA) and added to a 3% ring-stage NF54/iGP2 (to generate iGP2^230-tag^) or NF54 (to generate 230^ΔD13/D14^ / 230^ΔCterm^) culture. Parasites were electroporated (310 V, 950 µF), allowed to recover for 4 h, after which transfected parasites were selected using 2.5 nM WR99210 (Jacobus Pharmaceuticals) for five days. To obtain single-cell clones for iGP2^230-tag^, late-stage parasites were stained using 1 µg/ml Rhodamine123 solution, and single-cell sorted using a FACSAria III Cell Sorter (BD Biosciences) in a round-bottom 96 well plate containing 100 µl complete medium with 3% haematocrit. To obtain an isogenic population for 230^ΔD13/D14^ and 230^ΔCterm^, parasites were sorted based on mScarlet expression using a Cytoflex SRT Benctop Cell Sorter (Beckman Coulter). Successful integration was confirmed by diagnostic PCR (**Table S5**).

To induce gametocytogenesis, a 1% early trophozoite culture (±24 hours post invasion) was induced with AlbuMAX medium (RPMI1640 supplemented with 25 mM HEPES, 25 mM NaHCO_3_, and 0.5% AlbuMAX-II (Gibco, #11021-045)), in a previously described automatic shaker setup with media changes every 12 h (82, 83). Additionally, glucosamine was removed from iGP2-based cultures to further increase gametocyte induction. 36 h post induction until completion, cultures were maintained in complete medium. From day 4 – 8 post induction, gametocytes were cultured in the presence of 20 units/ml heparin (Sigma, #H3393) to eliminate asexual parasite growth. To obtain late-stage gametocyte saponin pellets, gametocytes were harvested on day 13 post induction by incubating the cells for 10 min in a 10x pellet volume ice-cold 0.06% (w/v) saponin in PBS supplemented with 1x Complete™ Protease Inhibitor (PI) Cocktail (Roche), and subsequent centrifugation at 3,000xg for 5 min at 4 °C. Pellets were washed three times in ice-cold PBS with 1x PI, flash frozen and stored at-70 °C.

### Western blot

Saponin pellets from late-stage gametocyte cultures were resuspended in lysis buffer (10 mM Tris-HCl (pH 8.0), 150 mM NaCl, 1% (w/v) sodium deoxycholate, and 1x PI) for 15 min at RT. Lysate was cleared by centrifugation, and supernatant was supplemented with 1x NuPAGE LDS buffer, and heated to 56 °C for 15 min. Samples were loaded on 4-12% Bis-Tris SDS-PAGE gels (SurePAGE), using Precision Plus Dual Color (BioRad) as a reference. After electrophoresis, proteins were transferred to 0.45 µm Immun-Blot PVDF membrane (Bio-Rad) using the Trans-Blot Turbo transfer system (Bio-Rad). Membranes were blocked in PBS with 3% BSA, before incubation with primary antibody diluted in PBS-T with 1% BSA. The following primary antibodies were used: 18F25 (mouse anti-Pfs230 (*36*), 5 µg/ml), 32F3 (mouse anti-Pfs48/45 (*25*), 5 µg/ml), M2 (mouse anti-FLAG, F1804 Sigma, 1:2000), CaptureSelect^TM^ (biotinylated single domain antibody fragment anti-C-tag, Thermofisher, 1:1000). After washing, blots were incubated with one of the following secondary antibodies: HRP-conjugated rabbit anti-mouse (P0260, Dako, 1:2000), HRP-conjugated streptavidin (890803, R&D Systems, 1:500). Blots were developed with Clarity Western ECL substrate (BioRad) and imaged on an ImageQuant^TM^ LAS 4000 (GE Healthcare).

### Immuno-fluorescence microscopy

Heparin-treated stage V gametocytes (day 13) were fixed (4% EM-grade paraformaldehyde, 0.0075% glutaraldehyde in PBS) for 20 min on pre-warmed poly-L-lysine coated coverslips at 37 °C. Cells were permeabilized with 0.1% Triton X-100, washed in PBS and blocked using 3% BSA in PBS for 1 h. Primary and secondary antibodies were incubated for 1 h at RT, with PBS washes in between. Primary antibodies were diluted in 1% BSA in PBS, and included M2 (mouse anti-FLAG, F1804 (Sigma), 1:500), RUPA-55 mAb (human anti-Pfs230 (*20*), 5 µg/ml), and 45.1 mAb (rat anti-Pfs48/45 (*25*), 5 µg/ml). Secondary antibodies (goat anti-mouse 594 (A11031), goat anti-human 488 (A11013), and chicken anti-rat 674 (A21472) (ThermoFisher)) were all diluted 1:500 in PBS with 1 µM DAPI. Coverslips were mounted using Vectashield® Antifade Mounting Medium (Vector Laboratories). Images were taken on a Zeiss LSM900 Airyscan confocal microscope using a 63x oil objective with 405, 488, 561, and 633 nm laser excitation and an Electronically Switchable Illumination and Detection Model (ESID) for transmitted light. Images were processed for airyscan, using the Zeiss Zen Blue software and analysed using FIJI software (84).

### Suspension Immunofluorescence Microscopy (SIFA)

For gamete SIFAs, heparin-treated stage V gametocytes (day 13 post-induction) were collected, spun down to remove the culture medium, and resuspended in half the culture volume of FCS. After incubation with gentle agitation at RT for 1 h, parasites were spun down and washed in ice-cold PBS. For saponin-treated gametocytes, heparin-treated stage V gametocytes (day 13 post-induction) were collected, spun down, and resuspended in half the volume of ice-cold PBS supplemented with 0.06% Saponin and 1x protease inhibitor. After 10 min incubation on ice, saponin-treated gametocytes were harvested by centrifugation and washed three times in ice-cold SIFA buffer (PBS supplemented with 0.5% FCS). All subsequent steps were performed at 4 °C.

Both gametes and gametocytes were incubated for 1 h in SIFA buffer with 15 µg/ml mAb 18F25 or 45.1, labeled with LYNX Rapid Plus DyLight488 Antibody Conjugation Kit (Bio-Rad). After washing in ice-cold SIFA buffer, parasites were resuspended in ice-cold PBS and deposited in pre-cooled poly-L-lysin-coated µ-slide 8 well Ibidi chambers. Parasites were allowed to settle at 4 °C, after which the Ibidi chamber was transferred to a room-temperature Axio Observer 7 Inverted LED microscope equipped with a Colibri 7 LED source and Axiocam 705 mono (Zeiss). Images were taken with a 63x oil objective using the 475 nm LED module and a Differential Interference Contrast enhanced brightfield channel. Images for different parasite lines were taken on the same day with the same microscope settings, and were analyzed with the same settings in FIJI software (84).

### Exflagellation assay and standard membrane feeding assay

Day 15 cultures of non-heparin treated gametocytes were used to test exflagellation. Cultures were mixed 1:1 with 50 µM xanthurenic acid in complete media, and incubated at room temperature in a dark, humid chamber for 15 min. The number of exflagellation centres per ml was determined by bright-field microscopy using a Neubauer chamber. To assess mosquito infectivity, day 15 non-heparin treated gametocytes were added to the bloodmeal of *Anopheles stephensi* mosquitoes (colony maintained at Radboudumc (Nijmegen, The Netherlands)), as described previously (85). In the SMFA experiments with 2A2, purified 2A2.2a IgG was diluted in FBS to a final concentration within the total bloodmeal of 10 µg/ml, and mixed with mature *P. falciparum* gametocytes and human serum that contains active complement. Mosquito midguts from 20 mosquitoes were dissected 6-8 days after the blood meal, stained with mercurochrome, and oocysts were counted by microscopy. Transmission reducing activity was calculated as described previously (86).

### Affinity purification of Pfs230:Pfs48/45 complex from stage V gametocytes

Frozen stage V gametocyte pellets were resuspended in solubilization buffer (25 mM HEPES pH 7.4, 150 mM KCl, 10% glycerol, 1x PI, and 0.25% (v/v) DDM), and rotated at 4 °C for 1 h. The cell suspension was centrifuged at 18,000xg at 4 °C for 30 min, after which the pellet was solubilized a second time following the previous steps. The combined supernatants were applied to anti-FLAG® M2 affinity resin (Merck), and incubated overnight at 4 °C. The resin was extensively washed in Wash buffer (25 mM HEPES pH 7.4, 150 mM KCl, 10% glycerol, 1x PI, and 0.02% DDM), after which proteins were eluted using 5x resin volume of elution buffer (25 mM HEPES pH 7.4, 150 mM KCl) supplemented with 0.02% DDM, and 150 µg/ml 3xFLAG peptide (Merck)). For cryo-EM studies, the eluted proteins were mixed with a fivefold molar excess of RUPA-97, LMIV230-01, 2A2, 18F25, RUPA-71, and RUPA-44 Fabs and incubated on ice for 30 min. After concentrating the protein fraction, the Fab-bound complex was separated from unbound Fabs by high-pressure size exclusion chromatography using a Bio SEC-3 300 Å column (Agilent, 5190-5213), pre-equilibrated with elution buffer.

### Expression and purification of Fabs, mAbs, and recombinant proteins

Expression and purification of individual Fabs was performed as described previously (20, 32). Variable light and heavy chains were gene synthesized (GeneArt), and cloned into a custom pcDNA3.4 expression vector directly upstream of the constant chains. The variable chain sequences for 32F3, TB31F, LMIV-230s, and all RUPA mAbs were described previously (20, 32, 41). Variable sequences for 18F25 (**Table S6**) were determined by a previously published workflow for hybridoma sequencing (87). In short, RNA was obtained from 18F25.1 hybridoma cells by Trizol purification, which was used as template for RT-PCR with heavy/kappa constant chain specific reverse primer and a universal template-switching oligonucleotide (**Table S5**). The obtained cDNA was used for a PCR, and obtained PCR products were inserted into a TOPO vector, and sequenced by Sanger Sequencing (Baseclear, The Netherlands). Variable sequences for 2A2 were sequenced by Genmab (**Table S6**). For the IgGs produced for complement-fixation assays, sequences of the variable chain of α-Pfs230D1 IgGs were described previously (20), and ordered as synthetic products subcloned into custom-made pcDNA3.4 vectors directly upstream of Igγ1-CH1-CH3 or Igκ-CL1.

Fab and IgG purification were done in a similar manner. Plasmids coding for Fab/IgG heavy chain and light chain were co-transfected at a 1:1 molar ratio using PEI MAX® transfection reagent (PolySciences) into Freestyle 293-F or 293-S cells (ThermoFisher). Cells were cultured in FreeStyle 293 Expression medium (Gibco) at 37 °C and 8% CO_2_ while shaking for seven days, after which the supernatant containing secreted recombinant protein was collected. After filtering, the supernatant was used for affinity purification using a HiTrap Kappaselect column (Cytiva) for recombinant Fab, or the HiTrap Protein G HP column (Cytiva) for recombinant IgG, pre-equilibrated in 1x PBS. In both cases, recombinant protein was eluted in 100 mM glycine (pH 2.2). Fabs were further purified by cation-exchange chromatography using a MonoS column (Cytiva) with a buffer of 20 mM sodium acetate (pH 5.6) across a 0-1.0 M potassium chloride gradient.

Single Pfs230 domains Pfs230D1 and Pfs230D10 were expressed and purified as described previously (36). Coding sequences for the double Pfs230 domain D13-14-Cterm (residue 2831-3105) were ordered and synthesized as *Drosophila melanogaster* codon-optimized sequences (Baseclear). The synthetic gene was subcloned into pExpreS2.2 plasmid (ExpreS2ion Biotechnologies), in frame with an N-terminal BiP signal peptide – His_6_ tag. The expression plasmid was used to transfect *D. melanogaster* S2 cells (ExpreS2ion Biotechnologies) to generate stable cell lines as described previously (36). Pfs230D13-14-Cterm-containing supernatant was loaded directly on a 1 ml cOmplete™ His-Tag Purification Column (Roche). The column was washed with PBS, and Pfs230D13-14-Cterm was eluted using PBS with 250 mM imidazole. After overnight dialysis against PBS, the sample was concentrated and further purified by size-exclusion chromatography using a Superdex200 Increase column (Cytiva), using PBS as running buffer.

### Blue Native Polyacrylamide Gel Electrophoresis

For BN-PAGE, the elution fraction from FLAG-based affinity purification, containing purified Pfs230:Pfs48/45 protein complex, was used. To test Fab binding to the complex, a tenfold molar excess of Fab fragment was added to the purified Pfs230:Pfs48/45 complex, and incubated on ice for 30 min. Protein samples were supplemented with a home-made 1x loading solution (final concentration 10% glycerol, 0.025% Coomassie Blue G-250), and loaded on a NativePAGE^TM^ 3-12% Bis-Tris protein gels (ThermoFisher) following manufacturer’s protocols. NativeMark^TM^ Unstained Protein Standard was used as marker. After electrophoresis, gels were either stained with Coomassie InstantBlue® protein stain (Abcam) or incubated in 0.1% SDS for western blotting. For western blotting, proteins were transferred to 0.45 µm Immun-Blot PVDF membrane (BioRad) using the Trans-Blot Turbo Transfer system (BioRad). After transfer, proteins were fixed to the membrane using 8% acetic acid. Membranes were air-dried, washed in pure methanol and re-hydrated using MQ. Blocking, antibody incubation and detection were performed as described above.

### Mass spectrometry

Three batches of 40 ml heparin-treated, late-stage gametocyte cultures (iGP2^230-tag^ and iGP2 WT) of comparable gametocytaemia were harvested and lysed in parallel, as described as above for purification of the endogenous Pfs230:Pfs48/45 complex. Protein concentration in parasite lysate was determined using the PierceTM BCA Protein Assay kit. 120 µg of protein extract in 300 µl solubilization buffer (25 mM HEPES pH 7.4, 150 mM KCl, 10% glycerol, 1x PI, and 0.25% (v/v) DDM) was applied on 10 µl anti-FLAG® M2 affinity resin, and incubated overnight. The resin was washed in ice-cold wash buffer (25 mM HEPES pH 7.4, 150 mM KCl, 10% glycerol, 1x PI, and 0.02% DDM) three times, and consequently washed three times in ice-cold PBS (Gibco). After harvesting, beads were resuspended in 50 μl elution buffer (100 mM Tris-HCl pH 8.0, 2 M Urea, 10 mM DTT), and incubated for 20 min at 25 °C with shaking. 50 mM iodoacetamide was added to alkylate cysteines, after which samples were kept in the dark for 10 min at 25 °C. After adding 0.25 µg sequencing grade trypsin (Promega), samples were incubated at 25 °C for 2 h with gentle shaking. Samples were spun down, supernatants were collected, and the beads were resuspended in 50 µl fresh elution buffer. After 5 min incubation on a shaker, the samples were spun down again and the supernatant combined with the previous elution fraction. Another 0.1 µg of fresh trypsin was added to the combined supernatants, which was left to digest overnight at 25 °C. The following day, samples were concentrated and purified on C18 StageTips (88). Samples were analyzed on a Obitrap Exploris 480 mass spectrometer (ThermoScientific) ran in Top20 mode (with dynamic exclusion enabled for 45 s), operated with an online Easy-nLC 1000. A gradient of buffer B (80% acetonitrile, 0.1% formic acid) was applied for 60 min. Raw data was analyzed using Maxquant software (version 2.1.4.0), and analyzed against a *Plasmodium* database (PlasmoDB, downloaded 21-10-2022). LFQ, iBAQ and match between runs were enabled, and deamidation (NQ) was added as additional variable modification. The resulting output was filtered using Perseus (version 1.0.15), removing potential contaminants, reverse hits, and proteins with less than two unique peptides or with less than three valid values in at least one group. Missing values were imputed using default settings, and a t-test was performed to identify outliers. Data was visualized using GraphPad Prism.

### Cryo-EM data collection and image processing

The Pfs230:Pfs48/45:RUPA-97:LMIV230-01:2A2:18F25:RUPA-71:RUPA-44 Fab complex was concentrated to 0.8 mg/ml, and 1.8 µl sample was deposited on homemade holey gold grids (89) that were glow-discharged in air for 15 s. Excess sample was blotted away for 1.7 s using a Leica EM GP2 Automatic Plunge freezer at 4 °C and 90% humidity, and plunge-frozen in liquid ethane. Data was collected on a Thermo Fisher Scientific Titan Krios G3 equipped with a Selectris X energy filter (slit width: 10 eV) and Falcon 4i camera, operated at 300 kV and automated with the EPU software. Data was collected at a magnification of 130,000× with a calibrated pixel size of 0.93 Å. Exposures were collected for 6.5 s as movies with a camera exposure rate of ∼7.11 or ∼7.15 e^−^ per pixel per s, and total exposure of ∼50 or ∼53.7 electrons/Å^2^ and recorded in Electron Event Representation mode (90). Before processing, movies were fractionated into 30 frames. A total of 8,545, 1,572, and 6,533 raw movies were obtained at 0°, 35°, and 40° tilt, respectively.

The collected cryo-EM data was processed using CryoSPARC v4.6.0 (91). The movies from the 35° tilted, 40° tilted, and untilted data collections were corrected for motion in patches, and contrast transfer function (CTF) parameters were estimated in patches. Resulting micrographs were curated using thresholds for average defocus values (4,000-35,000 Å), CTF fit resolution (2.0-5.0 Å), and relative ice thickness (0.95-1.10). An *in silico* model of the Pfs230:Pfs48/45:RUPA-97:LMIV230-01:2A2:18F25:RUPA-71:RUPA-44 complex was built from AF models and arranged based on views seen from 2D classes. The resulting model was used to generate particle templates corresponding to 40 evenly distributed projections which were low-pass filtered to 20 Å to avoid model bias. Initially, particle images were selected from a subset of 2,000 curated micrographs, which were then extracted in 450×450 pixel boxes and subjected to 2D classification. Classes where the average showed clear views of the complex were used to train a Topaz particle selection model (92), with subsequent 2D classification steps used to clean the dataset of selected particle images. To focus on the various subsections of the full complex, 2D class average images showing the appropriate domains were used as templates to select more particles from the full dataset of 12,192 curated micrographs. Further Topaz models were trained on specific regions (Pfs230D1-6:RUPA-97:LMIV230-01:2A2, Pfs230D9-14:Pfs48/45:RUPA-71:RUPA-44) of the full complex, after which particles images were selected and extracted with the trained models. Initial junk particle image removal was performed with multiple rounds of 2D classification, followed by *ab initio* 3D reconstruction. Non-particle images and poorly behaved particle images were removed with a combination of heterogeneous refinement, 3D classification, and 2D classification. Selected particle images were used to generate 3D maps with reconstructions in non-uniform refinement (93) jobs without enforced symmetry (C1). The resulting maps were further improved by local refinement and post-processing with DeepEMhancer (94). Heterogeneity for 2A2 Fab binding could be distinctly observed, such that two maps of Pfs230D1-6:LMIV230-01:RUPA-97 were obtained: with and without 2A2 Fab. A detailed processing flow path and experimental map details are available in the supplementary document (**Figure S2, 3A-3D**).

For the Pfs230D7-8:18F25 map, particle images from a subset of 500 micrographs were used from one set of collections of the untilted (2,003), 35° tilted (337), and 40° tilted (2,866) micrographs for blob picking after motion correction, CTF estimation, and curation of the micrographs (average defocus = 1000-30,000 Å; CTF fit resolution < 5 Å; relative ice thickness = 0.954-1.1; astigmatism < 4000 Å). The 101,225 particle stack was cleaned using multiple rounds of 2D classification, ab initio reconstruction, and homogeneous refinement (4.89 Å). The rebalance orientations job was used, followed by ab initio reconstruction and non-uniform refinement (4.19 Å). A map containing the interface between Pfs230 and 18F25 Fab was determined by cleaning the particle stack with 2D classification, followed by ab initio reconstruction and homogeneous refinement (10.03 Å). This map, along with the 4.19 Å map and a junk class generated with ab initio reconstruction were used as input for heterogeneous refinement, resulting in a class containing 18F25 with 35,162 particles. The resulting particle stack was cleaned with multiple rounds of 2D classification, with the final particle stack used for Topaz training on denoised micrographs. Ab initio reconstruction with five classes was run using the resulting 100,481 particle stack, followed by heterogeneous refinement. The particle stack from the class containing 18F25 was cleaned using 2D classification, then used again for Topaz training on all micrographs (6,372 untilted, 1,125 35° tilted, 2,866 40° tilted). To ensure the particles were centered on 18F25, ab initio reconstruction, followed by non-uniform refinement was run with the resulting 366,876 particle stack at a box size of 600 pix and Fourier crop of 300 pix (5.79 Å), followed by centering of the particles using volume alignment tools and re-extraction of the particles to 300 pix boxes. 2D classification without re-centering, 2 class ab initio reconstruction, and heterogeneous refinement were used to obtain a clean 216,684 particle stack. A further three rounds of 2D classification and two rounds of non-uniform refinement and local refinement resulted in a 3.88 Å map using 106,349 particles. Post-processing was performed in EMReady to prepare the map for model building. A detailed processing workflow and experimental map details are available in the supplementary document (**Figure S2, 3E**).

### X-Ray Crystallography

Fractions containing purified 2A2, 18F25, and RUPA-71 Fab following cationic exchange chromatography were pooled and concentrated at 16.8 mg/mL, 27.3 mg/mL, and 10 mg/ml, respectively in preparation for crystallization trials. RUPA-71 Fab was mixed in a 1:1 ratio with reservoir conditions from a JCSG Top96 screen (0.2 μL + 0.2 μL) using the sitting drop vapor diffusion method. RUPA-71 crystals then formed in a drop containing 0.2 M (NH4)_2_SO_4,_ 0.1 M MES 6.5 pH, and 30 %w/v PEG MME 5K. For the 2A2 and 18F25 Fab crystallization trials, reservoir conditions from a JCSG Top96 screen and Fab were mixed at a 1:1 volumetric ratio (0.3 μL + 0.3 μL) resulting in 0.6 μL drops. Approximately 48 h after setting drops at room temperature, 2A2 crystals were obtained from a drop containing reservoir solution: 30% (w/v) PEG 4K. 18F25 crystals were obtained from a drop containing reservoir solution: 0.2 M Na_2_SO_4_ and 20% (w/v) PEG 3350. The 2A2 and 18F25 crystals were cryo-protected with 25% glycerol (v/v) and 15% polyethylene glycol 200 (PEG200), respectively. Crystals of all three samples were flash-frozen in liquid nitrogen. Data collection on 2A2 and 18F25 was performed at the 17-ID-1 beamline at the Brookhaven National Laboratory Synchrotron Light Source (Beam wavelength = 0.920194 Å). RUPA-71 data collection occurred at the CMCF-ID beamline at the Canadian Light Source (Beam wavelength = 0.95357 Å).

### X-Ray Crystallography model building and refinement

Datasets were processed using autoproc (95) and xds (96). A molecular replacement solution was obtained using PhaserMR (97) using an *in silico-*derived model of the Fab domains predicted via ABodyBuilder2. Model building and refinement was performed using Coot (98) and phenix.refine (99). For the RUPA-71 Fab crystal structure, 98.0% of the dihedral angles are in “favoured regions” of the Ramachandran plot; 2.0% are in the “allowed regions;” and 0% are “outliers.” For the 2A2 Fab crystal structure, 98.3% of the dihedral angles are in “favoured regions” of the Ramachandran plot; 1.7% are in the “allowed regions;” and 0% are “outliers.” For the 18F25 Fab crystal structure, 97.4% of the dihedral angles are in “favoured regions” of the Ramachandran plot; 2.6% are in the “allowed regions;” and 0% are “outliers.” **(Table S3).** Inter-and intra-molecular contacts were determined using PISA (100) and manual inspection. Structural figures were generated using UCSF ChimeraX (101).

### Cryo-EM model building

Starting structural models were obtained by manually fitting previously determined structural models (PDB: 8U1P, Pfs48/45-D1D2; 7UXL, Pfs48/45-D3:RUPA-44; 7UVQ, Pfs230-D1:RUPA-97; and 7UFW, Pfs230-D1D2:LMIV230-01), novel crystal structures (Fabs of 18F25, RUPA-71, and 2A2), and models generated by AlphaFold2 (102)and AlphaFold3 (64) into the experimentally determined maps. Structural refinements were performed using Isolde (103), PyRosetta (104), and PHENIX (105); models were manually checked and improved with Coot (98). Images were generated using ChimeraX (101). Access to all software was supported through SBGrid (106).

### Single nucleotide polymorphism detection

Single nucleotide polymorphisms were obtained from the MalariaGEN Catalogue of Genetic Variation in Pf version 7 (53). Genotype calls for chromosomes 2 and 13 (Pf3D7_02_v3 and Pf3D7_13_v3, respectively) were downloaded (https://ngs.sanger.ac.uk/production/malaria/Resource/34/Pf7_vcf/). Bcftools (107)was used to subset calls and calculate the allele frequencies between nucleotide positions 369,351 and 380,156(+) on chromosome 2 that coincide with Pfs230 and between nucleotide positions 1,875,452 and 1,878,087(-) on chromosome 13 that coincide with Pfs48/45. SNPs calls below MalariaGEN’s quality filter (Low_VQSLOD) were removed before identification of missense variants.

### Gamete binding assay with flow cytometry

Heparin treated stage V gametocytes (day 13 – 16 post induction) were collected and resuspended in FCS with half the volume of the original culture volume. Gamete maturation was induced for 30 min at room temperature while shaking, after which samples were centrifuged at 2,000xg at 4 °C. The pellet was resuspended in PBS and placed on a layer of 12.4% Nycodenz (Serumwerk Bernburg) and centrifuged for 30 min at 3,500xg without break at 4°C. The top layer containing female gametes was harvested and washed in PBS before continuing.

50,000 female gametes per well were deposited in a V-bottom non-treated 96-well plate (Costar). All gamete incubations were carried out at RT in PBS supplemented with 2% Fetal Calf Serum (FCS, Gibco) and 0.02% sodium azide, and washed in ice-cold PBS. All secondary antibody incubations included eBioscience^TM^ Fixable Viability Dye eFluor^TM^ 780 (Invitrogen, 1:1000). For the Pfs230 domain binding experiment (**Figure S6A**), gametes were incubated for 1 h with recombinant proteins and washed in PBS. Afterwards, gametes were incubated with mouse anti-his tag antibody (Sigma-Aldrich, A7058, 1:200) for 1 h, washed, and stained with chicken-anti-mouse 488 secondary antibodies (Invitrogen, A21200, 1:200) for 30 min. For the antibody gamete binding experiment (**Figure S6H, Figure 5G**), gametes were incubated with 1 µg/ml antibody for 1 h, washed, and stained with goat-anti-human 488 secondary antibody (Invitrogen, A11013, 1:200). For the Fab competition experiment (**Figure S7C**), gametes were incubated for 1 h with Fabs. After washing in PBS, gametes were stained with 18F25 antibody labeled with LYNX Rapid Plus DyLight488 Antibody Conjugation Kit (Bio-Rad, 5 µg/ml). Finally, for complement deposition assay, gametes were incubated with 1 µg/ml antibody for 1 h and washed. Gametes were then incubated with 10% Normal Human Serum (Sanquin, The Netherlands) and incubated for 45 min at RT. After washing, parasites were first stained with polyclonal goat anti-human C1q (CompTech, 1:5,000) or anti-human C3 (CompTech, 1:10,000), washed, and finally stained with donkey-anti-goat 488 secondary antibody (Invitrogen, A11055, 1:200). After staining, gametes were resuspended in PBS and fluorescence was measured for approximately 2,000 gametes with the Gallios^TM^ 10-color system (Beckman Coulter), and analyzed with FlowJo (BD, version v10.10.0). For all gamete binding experiments only live gametes were analyzed, except for the complement deposition assays for which both live and dead gametes were included in the analysis (gating strategy in **Figure S7K**). Statistical analyses were performed using GraphPad Prism (version 10.4.1), and are described in the figure legends.

### DNA extraction and PCR analysis of midgut oocysts

Mosquito midguts were dissected into individual tubes seven days after the infectious blood meal, and DNA was extracted using a protocol modified from Ranford-Cartwrigth *et al.* (108). In short, the midgut was incubated at 56 °C overnight in oocyst lysis buffer (10 mM Tris-HCl pH 8.8, 100 mM NaCl, 25 mM EDTA, 0.5% N-lauroyl sarcosine, 1 mg/ml Proteinase K). DNA was extracted from the lysate using phenol/chloroform/isoamyl alcohol (25:24:1), followed by isopropanol precipitation. The DNA pellet was washed with 70% EtOH, dried and redissolved in 20 µl ultra-pure water. The presence of wildtype or truncated *pfs230* gene was confirmed with a semi-nested PCR protocol (**Table S5**) as described previously (38).

### Pfs230D7-18F25 SPR binding analysis

The gene sequences coding for Pfs230D7, including the SNP mutant sequences, were optimized for wheat codon usage, purchased from GenScript, and cloned into a pEU-E01 expression vector (CellFree Science, Matsuyama, Japan) (37). The recombinant Pfs230D7 proteins were expressed as C-terminally His-tagged proteins in wheat germ cell-free extract WEPRO7240H (CellFree Science), supplemented with Disulfide Bond Enhancer Enzyme Set (CellFree Science). Proteins were purified by single-step His-tag purification using a Ni Sepharose 6 Fast Flow column (Cytiva), following the manufacturer’s instructions.

The binding kinetics between recombinant Pfs230D7 proteins and 18F25 were measured using a Biacore X100 instrument (Cytiva). The antibody 18F25 was immobilized on the sensor chip using the Mouse Antibody Capture Kit (Cytiva) according to the manufacturer’s instructions. Recombinant Pfs230D7 proteins were injected in HBS-EP+ running buffer (Cytiva) using a single-cycle kinetic format. Data were analyzed with Biacore X100 Evaluation Software, and kinetic parameters were calculated using a 1:1 binding model.

## SUPPLEMENTARY FIGURES

**Figure S1:**
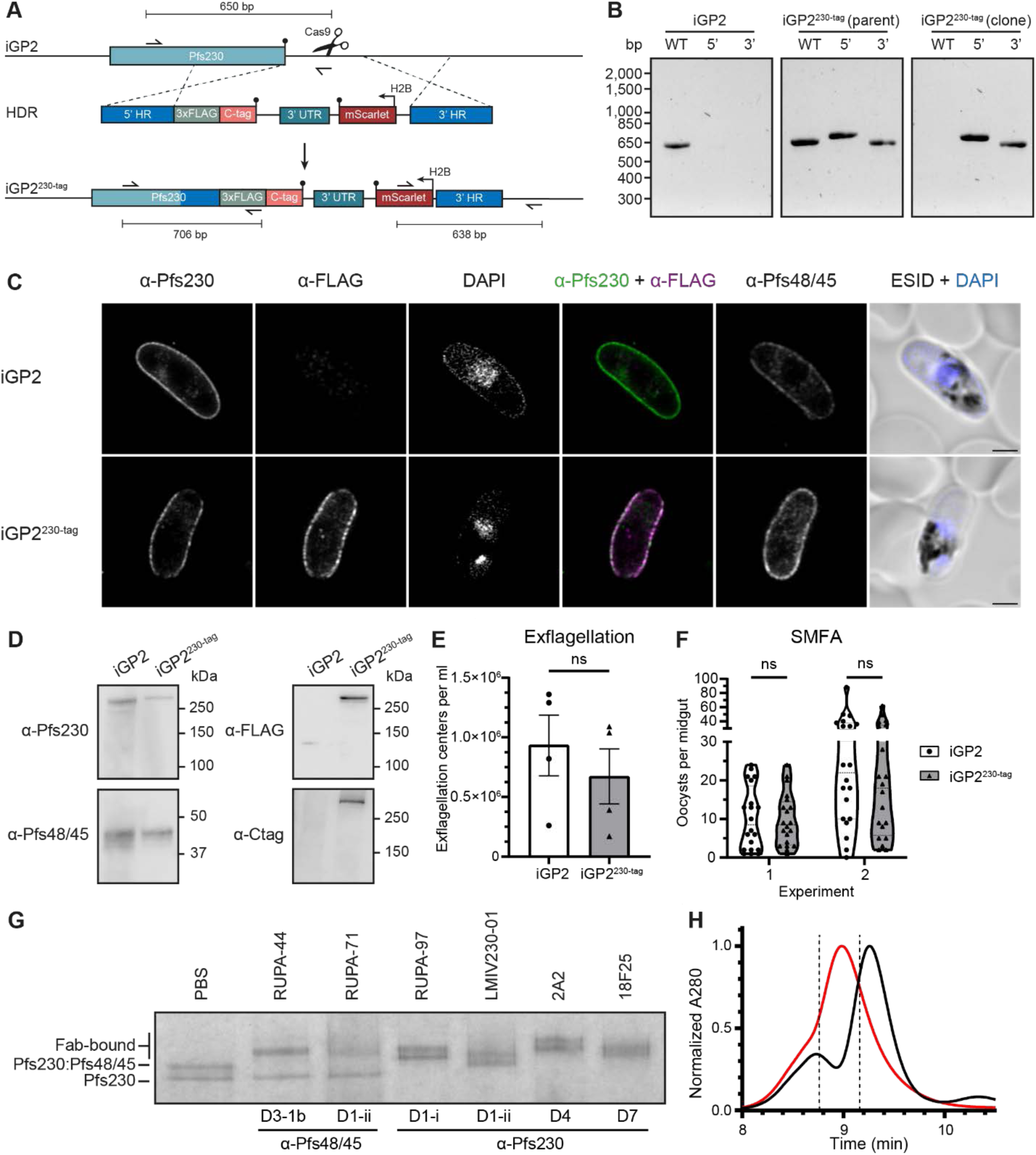
The iGP2^230-tag^ parasite line allows for purification of endogenous Pfs230:Pfs48/45 complex recognized by transmission-blocking antibodies. **(A)** Schematic overview of the genomic modification to obtain the iGP2^230-tag^ parasite line. HDR = Homology Directed Repair, HR = homology region, 3xFLAG = triple FLAG tag, C-tag = EPEA* tag, 3’UTR = bidirectional 3’untranslated region of PBANKA_142660, H2B = promoter of Pf3D7_1105100. **(B)** Genomic integration PCR to confirm 5’ and 3’ integration and the absence of wildtype parasites. Primers and expected PCR product are indicated in (A). **(C)** Immunofluorescence microscopy images of paraformaldehyde/glutaraldehyde-fixed and permeabilized iGP2 wildtype and iGP2^230-tag^ stage V gametocytes. Parasites were stained for Pfs230 (green, RUPA-55), FLAG-tag (magenta, M2), and Pfs48/45 (45.1), and DAPI (blue). Scalebar represents 5 µm. ESID = Electronically Switchable Illumination and Detection brightfield image. **(D)** Western blot analysis of iGP2 and iGP2^230-tag^ to confirm FLAG-tag and C-tag integration. **(E)** Exflagellation of stage V iGP2 (white) and iGP2^230-tag^ (gray) gametocytes. Bars represent mean ± standard deviation of number of exflagellation centers per ml from four independent cultures (dots). No statistical significance was found using an unpaired t-test. **(F)** Transmission of iGP2 (white) and iGP2^230-tag^ (gray) to mosquitoes in two independent standard membrane feeding experiments. Dots represent the number of oocysts per midgut *Anopheles stephensi* mosquitoes (n=20 per experiment) in two independent standard membrane feeding assays using iGP2 (white) and iGP2^230-tag^ (gray) parasites. Statistical analysis was done using a Mann-Whitney test with Holm-Šídák’s multiple comparisons test (α = 0.05). ns = not significant. **(G)** Coomassie blue stained Blue Native PAGE gel of purified Pfs230:Pfs48/45 (and free Pfs230) incubated with molar excess of the following individual Fab fragments: RUPA-44, RUPA-71, RUP-97, LMIV230-01, 2A2, or 18F25. Epitopes are defined in Figure 1A. **(H)** Normalized absorption at 280 nm during high-pressure liquid size-exclusion chromatography of Pfs230:Pfs48/45 (black) or Pfs230-Pfs48/45:6Fab (red) complexes. Dotted lines indicate fractions of eluted Pfs230:Pfs48/45:6Fab that were used for cryo-electron microscopy. *Related to* Figure 1.

**Figure S2:**
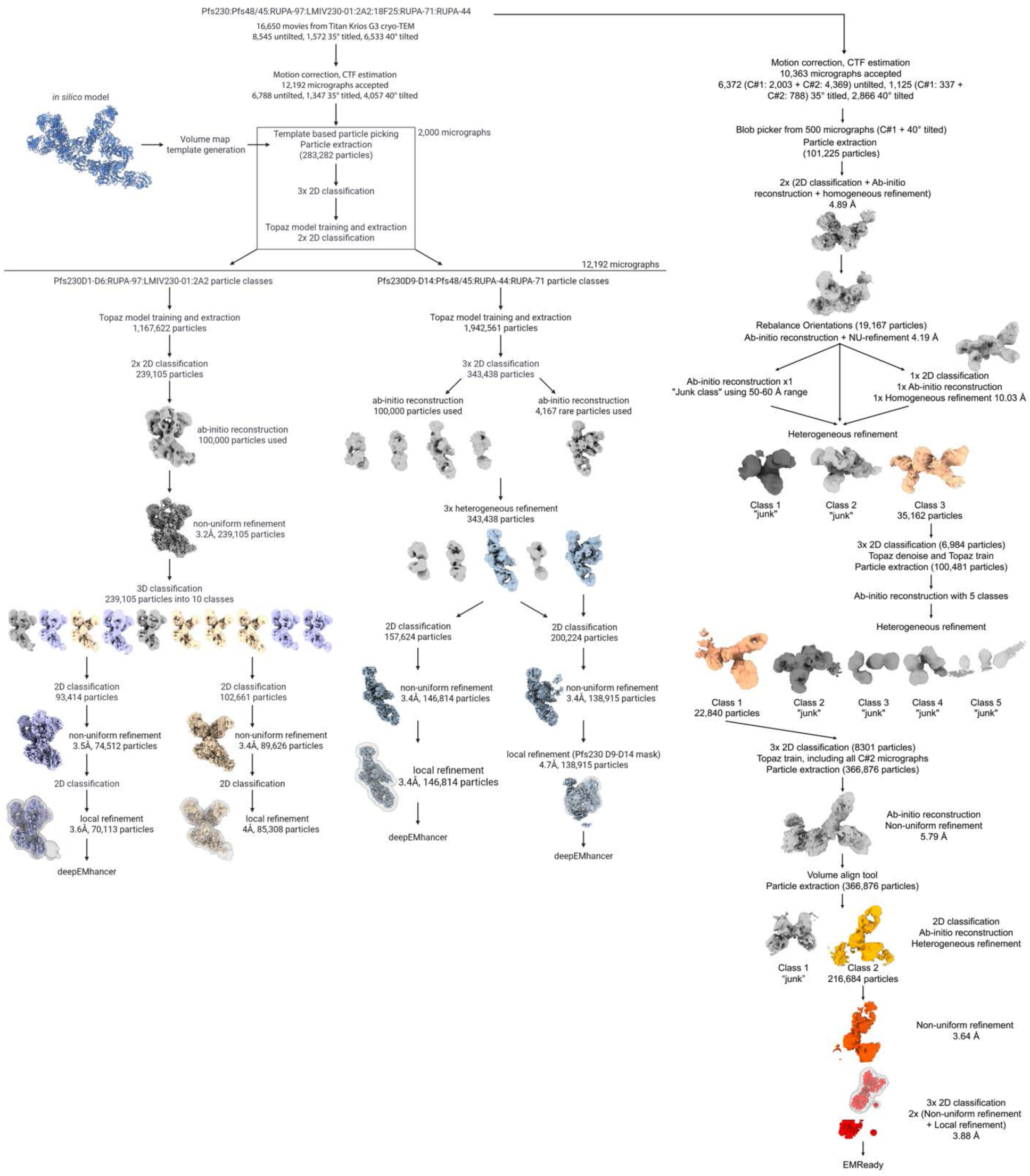
**Cryo-EM processing workflow**. Shown from left to right are the maps of Pfs230D1-6:LMIV230-01:RUPA-97:2A2, Pfs230D1-6:LMIV230-01:RUPA-97, Pfs230D13-14:Pfs48/45:RUPA-44:RUPA-71, the focused refinement for Pfs230D9-14:Pfs48/45, and the Pfs230D7-8:18F25 map. Abbreviations for the two data collection sessions are abbreviated in the Pfs230D7-8:18F25 workflow (C#1: collection C#2: collection 2). *Related to* Figure 1.

**Figure S3:**
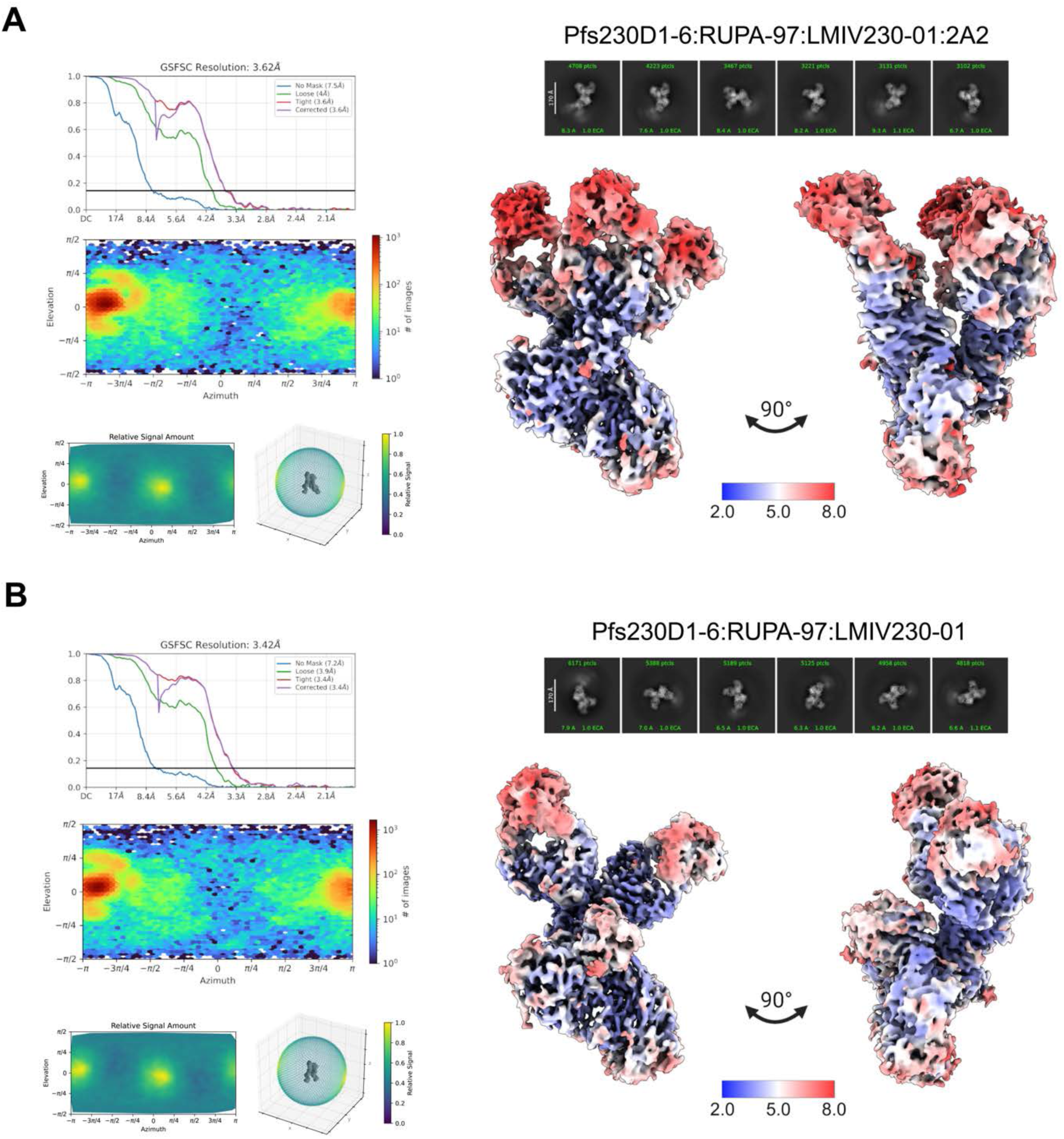
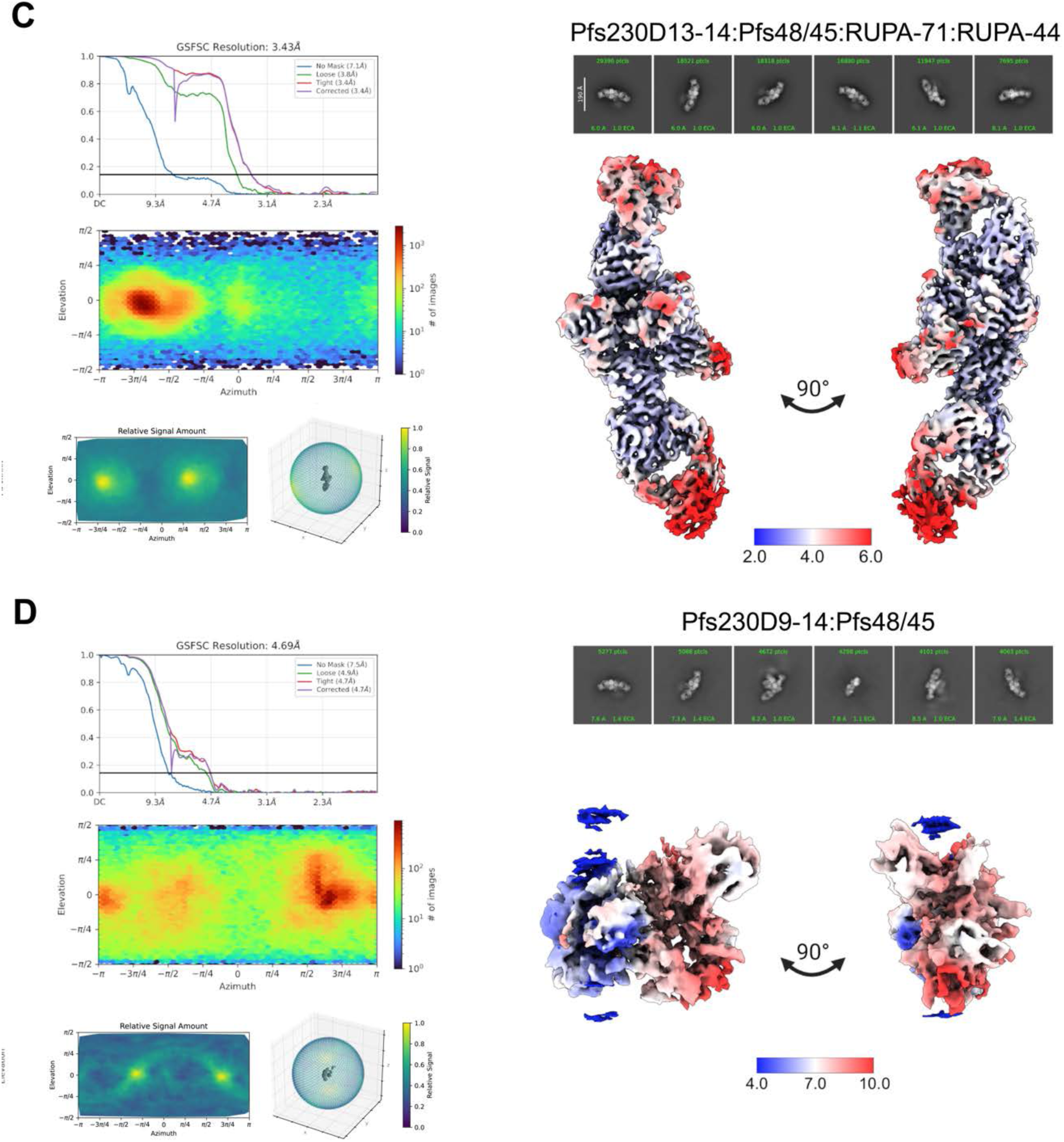
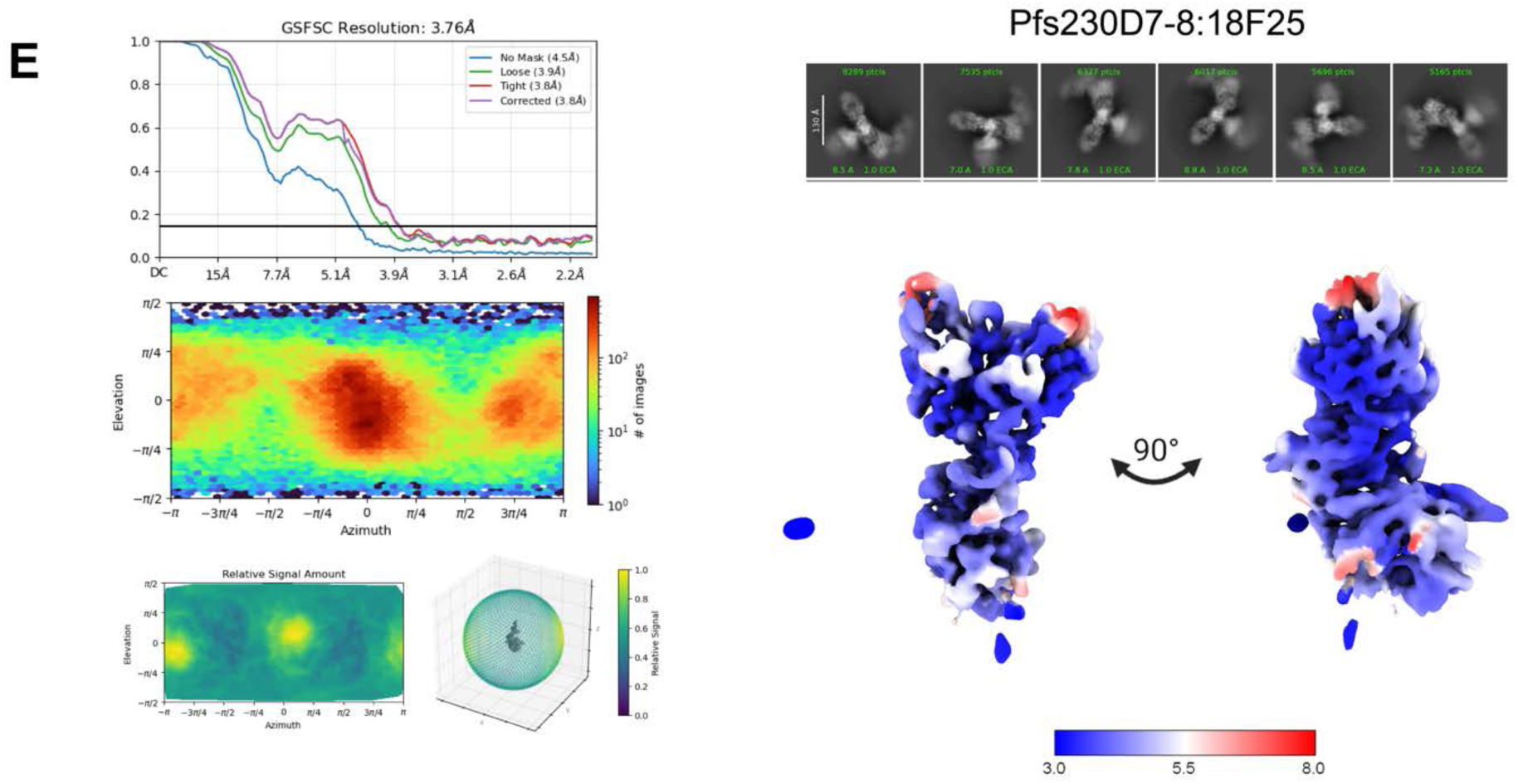
**Cryo-EM map validation data**. **(A-E)** The structures of (A) Pfs230D1-6:RUPA-97:LMIV230-01:2A2; (B) Pfs230D1-6:RUPA-97:LMIV230-01; (C) Pfs230D13-14:Pfs48/45:RUPA-44:RUPA-71; (D) focused refinement for Pfs230D9-14:Pfs48/45; and (E) Pfs230D7-8:18F25. The Fourier shell correlation curves following a gold-standard refinement with correction for the effects of masking for varying masks, the viewing direction particle distribution, and orientation diagnostic plots are shown (left panel). Additionally, representative 2D classes and cryo-EM maps coloured by local resolution are shown for each processed map (right panel). *Related to* Figure 1.

**Figure S4:**
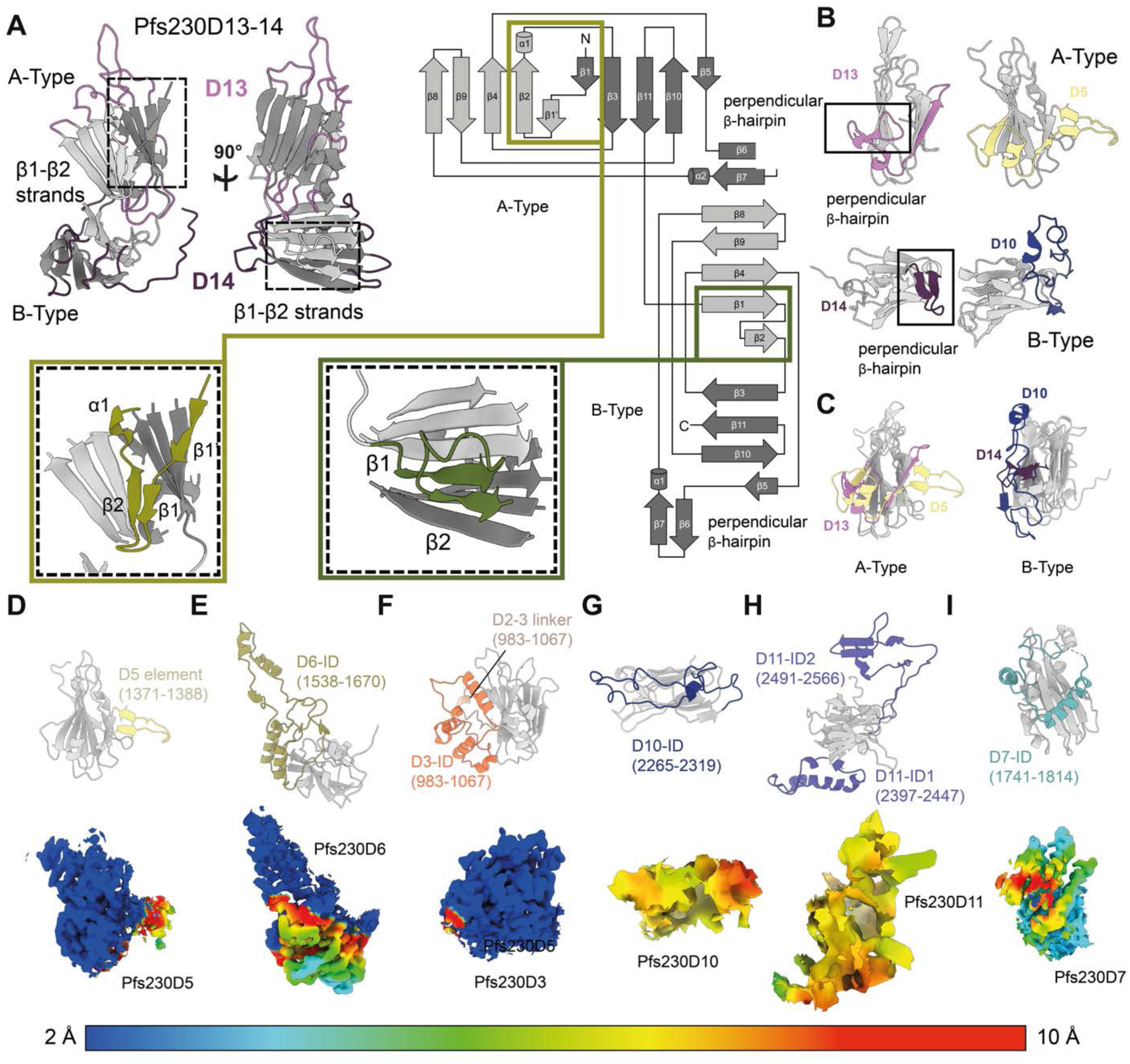
Structural features of Pfs230 6-Cys domains and Pfs230 domain clustering. **(A)** Pfs230D13-14 is shown as prototypical tandem A-and B-type 6-Cys domains of Pfs230. Loops of D13 and D14 are coloured according to the colouring scheme in Figure 1. The first β-sheet of the A-type and B-type domains is coloured distinctly (light gray) from the second β-sheet (gray). The distinguishing features of the A-and B-type domains (β1 to β2 strands) are indicated through the dashed box. Secondary structure topology of the two 6-Cys domains is shown with distinguishing features of A-(olive) and B-type (green) domains (β1 to β2 strands) indicated (inset) with a 3D representation of these differences shown. **(B-C)** Comparison of prototypical tandem 6-Cys domains (D13 and D14) to D5 and D10 respectively, (B) side by side and (C) overlaid, highlighting the structural differences of D5 and D10 at the perpendicular β-hairpin region. The regions that are structurally distinct are coloured according to the scheme in Figure 1. **(D-I)** Pfs230 domains that harbour IDs (Pfs230D3, D6, D7, D10, and D11) or structural elements contributing to ID clustering (Pfs230D3 and D5) are shown as ribbon representations with the cryoEM map corresponding to that portion of the model being shown underneath (shown at 5 Å radius around the models) and coloured according to local resolution (scale bar below). Non-canonical elements and IDs are coloured according to the colouring scheme specified in Figure 2. *Related to* Figure 2.

**Figure S5:**
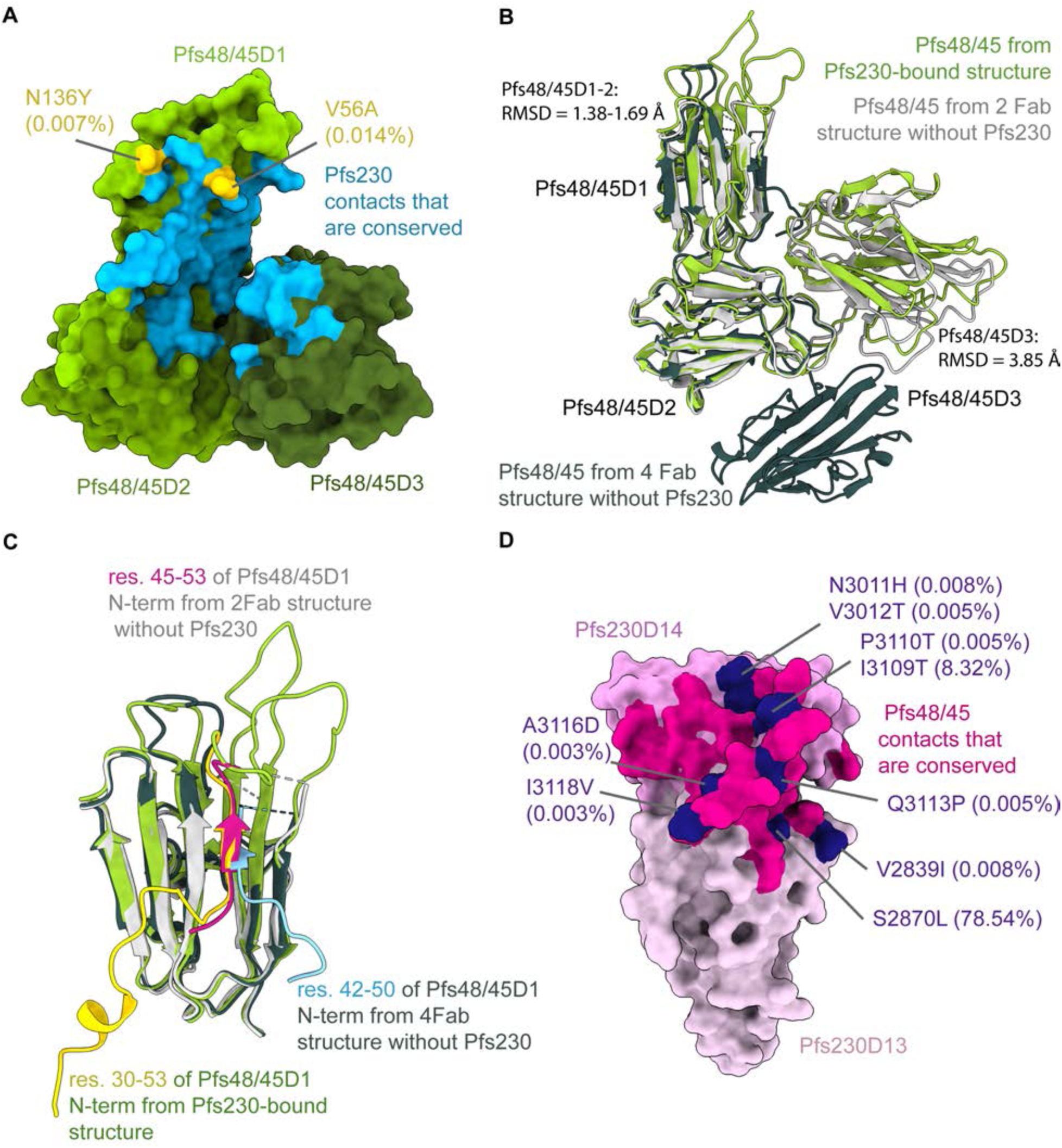
Comparison of Pfs48/45 conformation across determined structures and sequence conservation of Pfs230:Pfs48/45 binding site. **(A)** Structure of Pfs48/45 depicted as surface (yellow green, olive drab, and dark olive drab for Pfs48/45D1-3, respectively) with the Pfs230 binding site shown in sky blue and single nucleotide polymorphisms present within the epitope shown in gold. **(B)** Overlay of Pfs48/45 structures from Pfs48/45-Pfs230-6Fab complex (yellow green), Pfs48/45-4Fab complex (dark slate grey, PDB ID: 8U1P), and Pfs48/45-2 Fab complex (light grey, PDB ID: 7ZXF) aligned to Pfs48/45D1-2. **(C)** Overlay of N-terminal region of Pfs48/45-Pfs230-6Fab complex (green, yellow N-term), Pfs48/45-4Fab complex (dark slate grey, light blue N-term, PDB ID: 8U1P), and Pfs48/45-2 Fab complex (light grey, dark violet N-term, PDB ID: 7ZXF) aligned to Pfs48/45D1. **(D)** Pfs230D13 (thistle) and D14 (plum) shown as surface with the Pfs48/45 binding site in dark pink and SNPs in indigo. *Related to* Figure 3.

**Figure S6:**
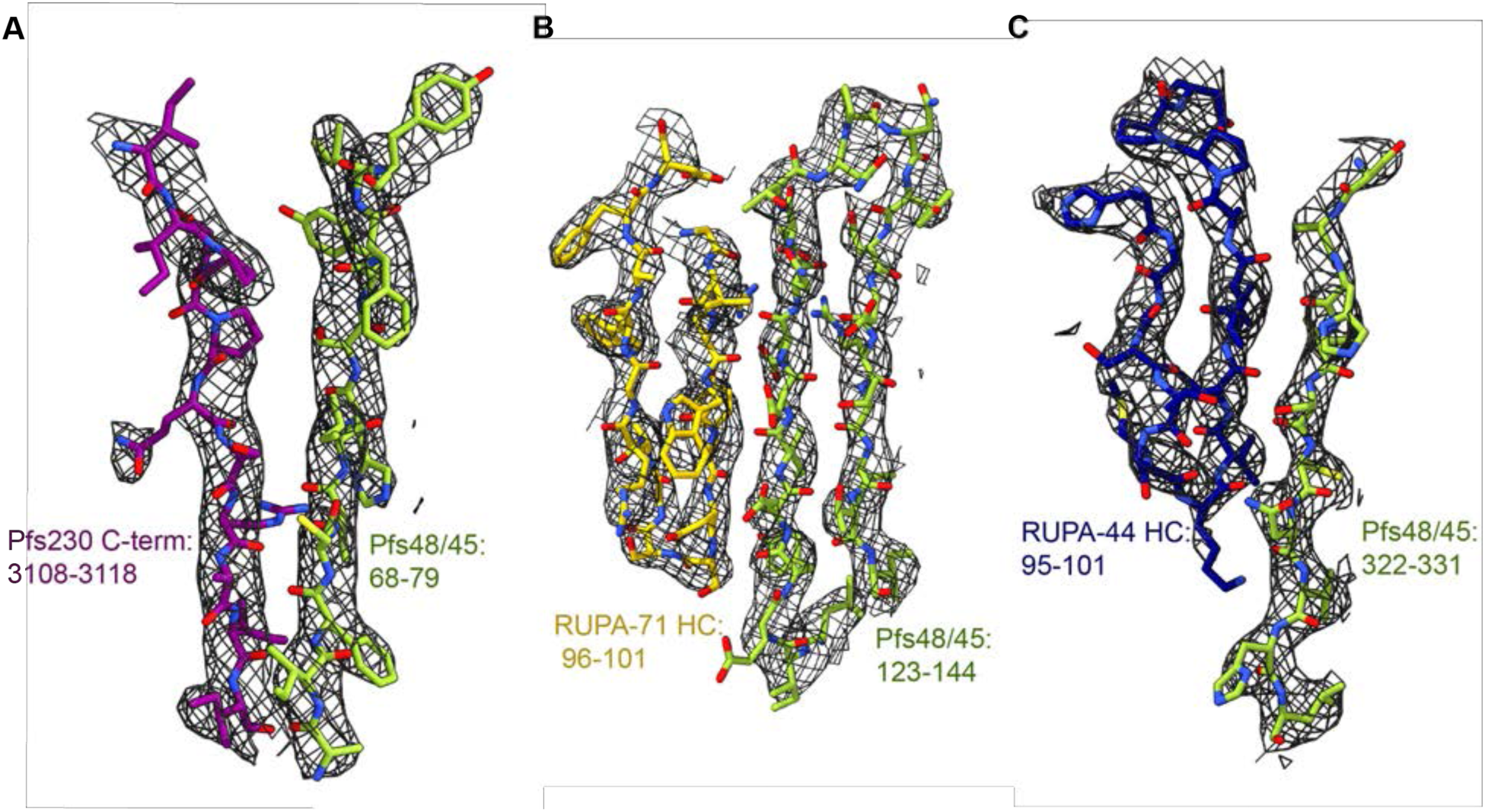
Cryo-EM map density around Pfs230:Pfs48/45:RUPA-71:RUPA-44 model. (A-D) Experimental map in grey mesh around key regions, including (A) the Pfs230 C-terminus and several of its Pfs48/45 contact residues, (B) residues at the Pfs48/45D1-RUPA-71 interface, and (C) residues at the Pfs48/45D3-RUPA-44 interface. Pfs48/45 (green), Pfs230D13-14 (purple), RUPA-71 (gold), and RUPA-44 (dark blue) are depicted as sticks. *Related to* Figure 3*-4*.

**Figure S7:**
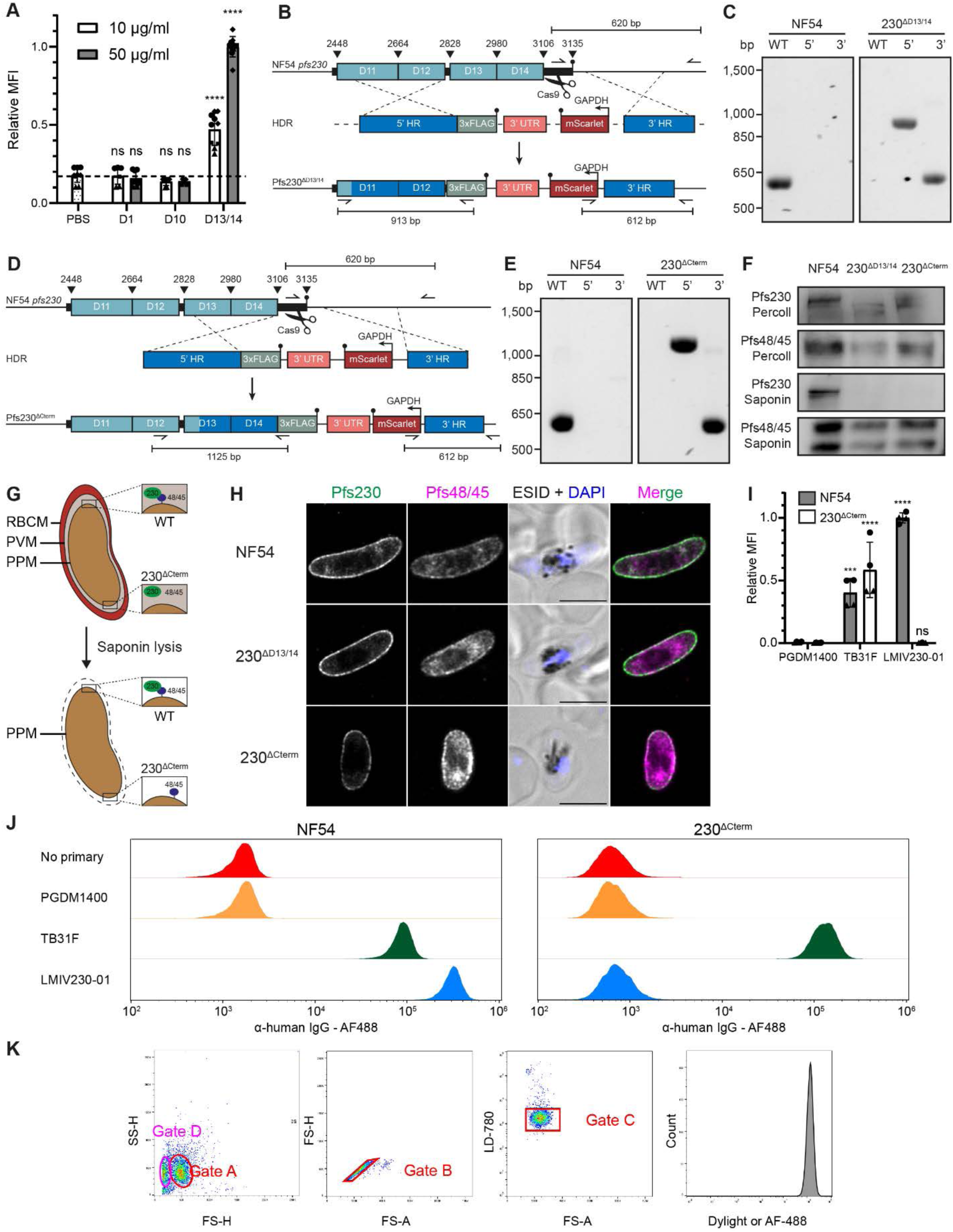
Pfs230:Pfs48/45 interactions are mediated by the C-terminal region Pfs230D13-14-Cterm *in vivo*. **(A)** Female gamete binding assay with recombinantly purified Pfs230D1, Pfs230D10 and Pfs230D13/14-Cterm at 10 µg/ml (white bar) or 50 µg/ml (gray bar). Bars show mean MFI (Mean Fluorescence Intensity) of two to four independent experiments with two to three technical replicates each. Error bars show standard deviation. Individual data points are shown, with different symbols for each independent experiment. Data was normalized against the mean of the highest value (50 µg/ml D13-14) in each independent experiment to allow for comparison across multiple experiments. Statistical analysis was done by comparing each group to the PBS control using an ordinary one-way ANOVA with Dunnett’s multiple comparisons test with a single pooled variance. **(B-E)** Schematic overview of genomic integration of Pfs230^ΔD13/14^ (B) and Pfs230^ΔCterm^ (D), and the corresponding diagnostic integration PCR (C, E). HDR = Homology Directed Repair, HR = homology region, 3xFLAG = triple FLAG tag, 3’UTR = bidirectional 3’untranslated region of PBANKA_142660, GAPDH = promoter of Pf3D7_1462800. **(F)** Western blot analysis of Pfs230 (18F25) and Pfs48/45 (32F3) expression in wildtype, Pfs230^ΔD13/14^ and Pfs230^ΔCterm^ late-stage gametocytes, isolated with Percoll (intact RBC and PVM membrane) or saponin (permeabilized RBC and PVM membrane). **(G)** Schematic illustration of localization of the Pfs230:Pfs48/45 complex on the parasite plasma membrane (PPM), and the effect of saponin on the red blood cell membrane (RBCM) and parasite vacuole membrane (PVM). **(H)** Representative immunofluorescence microscopy images of paraformaldehyde/glutaraldehyde-fixed NF54 wildtype, Pfs230^ΔD13/14^ and Pfs230^ΔCterm^ stage V gametocytes. Parasites were stained for Pfs230 (green, RUPA-55), Pfs48/45 (magenta, 45.1), and DNA (DAPI, blue). Scale bar is 5 µm. ESID = Electronically Switchable Illumination and Detection brightfield image. **(I)** Binding assay with wildtype (white bar) and Pfs230^ΔCterm^ (gray bar) female gametes, testing the binding of PDGM1400 (α-HIV-1 envelope glycoprotein), TB31F (α-Pfs48/45), and LMIV230-01 (α-Pfs230) antibodies at 1 µg/ml. Bars show the mean relative MFI of two independent experiments with two technical replicates each (±2000 live gametes per condition), error bars show standard deviation. MFI was normalized against the average of LMIV230-01 in the independent experiments. MFI values were compared within each parasite line to the PGDM1400 control using an ordinary two-way ANOVA and Šídák’s multiple comparison test with a single pooled variance. ns = not significant; ***=p<0.001, ****=p<0.0001. **(J)** Representative histogram of female gamete binding assay as shown in Figure S6I, comparing NF54 wildtype (top) and Pfs230^ΔCterm^ (bottom) gametes. **(K)** Exemplary plots to provide an overview for the gating strategy of macrogamete flow cytometry experiments. Live gametes (gate A) were gated based on side-scatter (SS) and forward scatter (FS). Single cells (gate B) were selected based on FS area (FS-A) versus height (FS-H), after which live cells (gate C) were selected based on absence of LD efluor780 staining. The geometric mean fluorescence intensity of the Dylight/AlexaFluor-488 channel of the resulting population was then used as the mean fluorescence intensity. Note that both gate A and D (containing dead gametes) were used in the case of the complement deposition assays, when no live/dead stain was used. *Related to* Figure 3.

**Figure S8:**
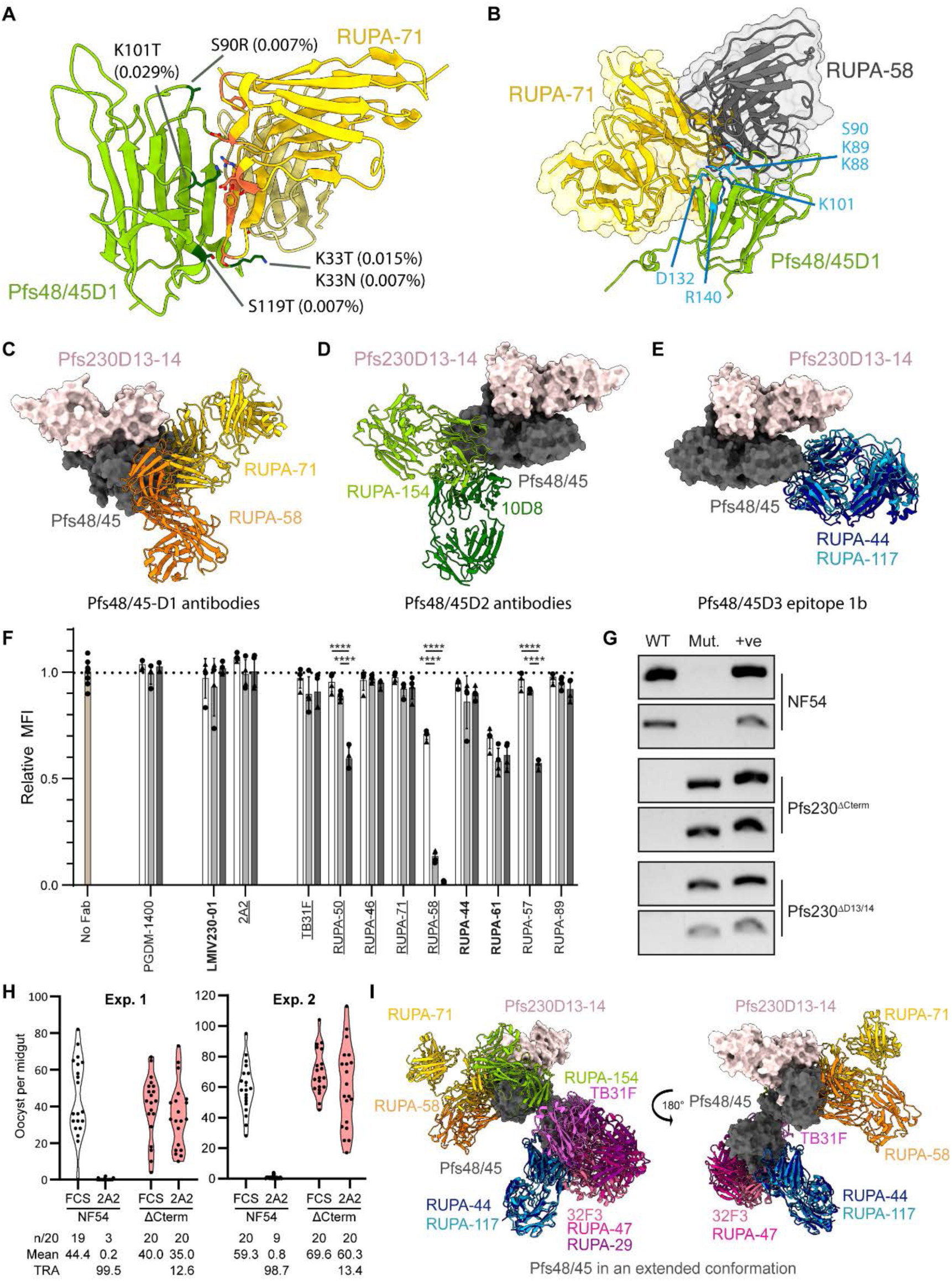
RUPA-71 epitope comparison and Pfs48/45-targeted Fab-induced dissociation of the Pfs230:Pfs48/45 complex *in vivo*. **(A)** RUPA-71 (yellow) bound to Pfs48/45 (yellow green) with Pfs48/45 residues with single nucleotide polymorphism in dark green and RUPA-71 residues that contact them in coral. **(B)** Overlay of Pfs48/45D1 (yellow green, cartoon) bound to antibodies RUPA-71 (gold) and RUPA-58 (dim gray; PDB ID: 8U1P). Pfs48/45 residues that contact both antibodies are indicated in blue. **(C-E)** Model of Pfs48/45 (dark gray, depicted in surface) bound to Pfs230D13-14 (misty rose, depicted in surface) in the disc-like conformation bound to (C) Pfs48/45D1 binders RUPA-71 (gold), RUPA-58 (orange, PDB ID: 8U1P)); (D) Pfs48/45D2 binders 10D8 (dark green, PDB ID: 7ZXF), RUPA-154 (yellow green, PDB ID: 8U1P)); (E) Pfs48/45D3-1B binders (RUPA-44 (dark blue), RUPA-117 (sky blue, PDB ID: 7UNB)). **(F)** Live female *P. falciparum* NF54 macrogametes were incubated with increasing concentrations of Fab fragments (white bar: 1 µg/ml; light grey: 10 µg/ml; dark grey: 100 µg/ml), after which Pfs230 surface retention was measured by determining the binding of 18F25-DyLight488 by flow cytometry. MFI was normalized against the no Fab control to allow for averaging across experiments (two biological replicates with two technical replicates each). Bars depict mean ± standard deviation, different symbols depict different biological replicates. Anti-Pfs48/45 Fabs are sorted on potency as determined by the IC_80_ value (Underlined: IC_80_ < 10 µg/ml; Bold: IC_80_ = 10-100 µg/ml, Others: IC_80_ > 100 µg/ml, see **Table S**1). Statistical analysis to test for a dose-dependent reduction in 18F25-488 binding by comparing all concentrations per individual Fab using an ordinary two-way ANOVA with a Šídák’s multiple comparisons test with a single pooled variance as one family. Only significant comparisons are shown. **** = p<0.0001. **(G)** PCR analysis of genomic DNA isolated from midgut oocysts from standard membrane feeding assay experiments of Pfs230 wildtype and truncation parasite lines. Semi-nested PCRs were used to detect: WT = wildtype Pfs230 genomic DNA; Mut = Pfs230 truncation genomic DNA; +ve = positive control for *P. falciparum* genomic DNA, amplifying the Pfs25 gene (PF3D7_1031000). **(H)** Raw standard membrane feeding assay data of NF54 wildtype (white) and Pfs230^ΔCterm^ (pink) with and without the addition of 10 µg/ml 2A2 and active human complement. TRA was calculated as percentage of reduction in mean oocyst per midgut compared to the FCS control. n/20 represents the number of mosquitoes that had at least 1 oocyst, “mean” is the average number of oocyst per mosquito midgut. **(I)** Model of Pfs48/45 (dark gray, depicted in surface) bound to Pfs230D13-14 (misty rose, depicted in surface) in the extended conformation overlayed with RUPA-71 (gold), RUPA-58 (orange, PDB ID: 8U1P), RUPA-154 (green, PDB ID: 8U1P)), RUPA-44 (dark blue), RUPA-117 (sky blue, PDB ID: 7UNB), TB31F (orchid, PDB ID: 6E63), RUPA-29 (magenta, PDB ID: 7UXL), RUPA-47 (violet red, PDB ID: 7UNB), and 32F3 (pale violet red, PDB ID: 7ZWI)). *Related to* Figure 4.

**Figure S9.**
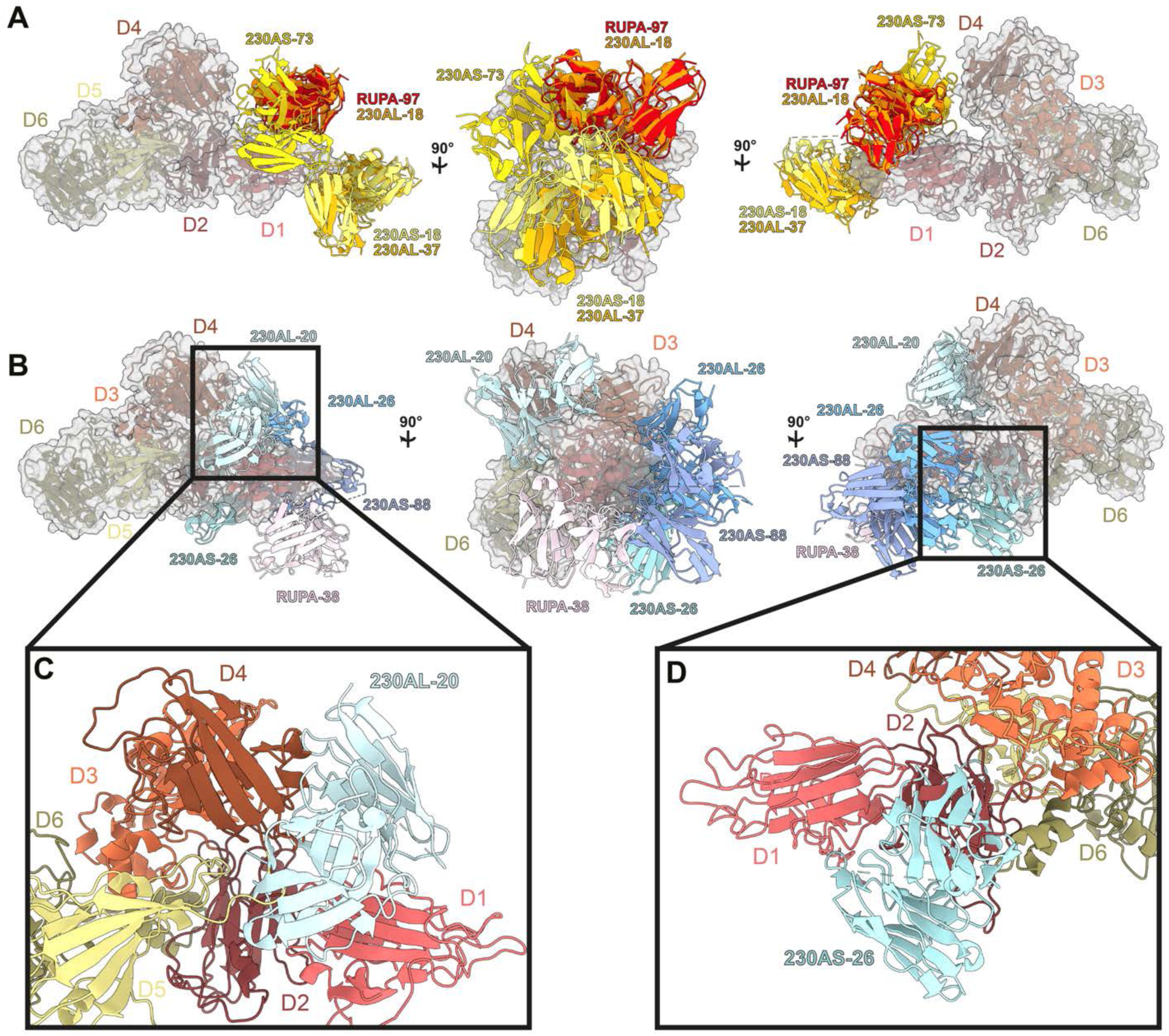
Overlay of Pfs230D1-EPA-elicted human mAbs. (*17*). **(A)** Potent mAbs (TRA > 80% at 100 μg/mL in SMFA; shades of yellow and orange) are shown overlaid on Pfs230D1-6 (surface: gray, ribbons: coloured as done in Figure 1) compared to RUPA-97 (red). (**B-D**) Non-potent mAbs (TRA < 80% at 100 μg/mL in SMFA; shades of blue) are shown overlaid on Pfs230D1-6 compared to RUPA-38 (light pink) (B). Zoom in of (C) 230AL-20 and (D) 230AS-26 showcasing steric clashes that occur with Pfs230D4 and Pfs230D2, respectively – as examples of mAbs elicited via a single Pfs230 domain subunit vaccine exposing non-native epitopes that would be incompatible with full-length Pfs230 binding. *Related to* Figure 6.

## SUPPLEMENTARY TABLES

**Table S1.** Overview of antibodies used or described in these studies. Shown are the targeted epitopes, structures in the RSCB/PDB database (structures in bold: this work), and estimates of the IC_80_ value (antibody concentration required to achieve 80% TRA in Standard Membrane Feeding Assay). ^a^ Estimate value from limited number of observations. ^b^ No blocking in High-Throughput membrane feeding assay at 100 µg/ml (32). *Related to Figures 1, 4, and 6*.

| Target | Antibody | Epitope | IC<sub>80</sub> (µg/ml) | Reference | Structure |
| --- | --- | --- | --- | --- | --- |
| Pfs230 | RUPA-97 | D1 – i | 0.7 | Ivanochko<sup>(20)</sup> | 7UVQ/**9N5H** |
|  | 4F12 | D1 – i | 55 | MacDonald<sup>(34)</sup>, Ivanochko<sup>(20)</sup> | 6OHG |
|  | LMIV230-01 | D1 – ii | 48 | Coelho<sup>(19)</sup>, Ivanochko<sup>(20)</sup> | 7UFW/7JUM/**9N5H** |
|  | RUPA-38 | D1 – ii | >>100 | Ivanochko<sup>(20)</sup> | 7UVI |
|  | LMIV230-02 | D1 – iii | >>100 | Coelho<sup>(19)</sup>, Ivanochko<sup>(20)</sup> | 7UVS |
|  | 15C5 | D1 – iii | >>100 | Lee<sup>(75)</sup>, Ivanochko<sup>(20)</sup> | 7UVQ |
|  | 2A2 | D4 | 1.9 | Roeffen<sup>(49)</sup>, De Jong<sup>(38)</sup> | **9N5H/9N89** |
|  | 18F25 | D7 | 10 to 30 <sup>a</sup> | Roeffen<sup>(49)</sup>, Inklaar<sup>(37)</sup> | **9N5O/9N6U** |
| Pfs48/45 | RUPA-58 | D1-i | 7.5 | Kucharska<sup>(15)</sup> | 8U1P |
|  | RUPA-94 | D1-i | <100 <sup>a</sup> | Fabra-García<sup>(32)</sup> | - |
|  | RUPA-46 | D1-ii | 2 to 10 <sup>a</sup> | Fabra-García<sup>(32)</sup> | - |
|  | RUPA-71 | D1-ii | 2 to 10 <sup>a</sup> | Fabra-García<sup>(32)</sup> | **9N5I/9N7K** |
|  | RUPA-154 | D2 | 103.1 | Kucharska<sup>(15)</sup> | 8U1P |
|  | RUPA-160 | D2 | ± 10 <sup>a</sup> | Fabra-García<sup>(32)</sup> | - |
|  | 10D8 | D2 | >350 <sup>a</sup> | Ko<sup>(14)</sup> | 7ZXF/7ZXG/7ZWM |
|  | TB31F | D3 - 1a | 0.86 | Kundu<sup>(41)</sup>, Kucharska<sup>(15)</sup>, Ko<sup>(14)</sup> | 6E63/7ZXF/6H5N |
|  | RUPA-29 | D3 - 1a | 0.4 to 2 <sup>a</sup> | Fabra-García<sup>(32)</sup> | 7UXL |
|  | RUPA-50 | D3 - 1a | 0.4 to 2 <sup>a</sup> | Fabra-García<sup>(32)</sup> | - |
|  | RUPA-47 | D3 - 1a | 10 to 100 <sup>a</sup> | Fabra-García<sup>(32)</sup> | 7UNB |
|  | RUPA-57 | D3 - 1a | >100 <sup>b</sup> | Fabra-García<sup>(32)</sup> | - |
|  | RUPA-72 | D3 - 1a | >100 <sup>b</sup> | Fabra-García<sup>(32)</sup> | - |
|  | RUPA-89 | D3 - 1a | >100 <sup>b</sup> | Fabra-García<sup>(32)</sup> | - |
|  | 32F3 | D3 - 1ab | <25 <sup>a</sup> | Vermeulen<sup>(25)</sup>, Ko<sup>(14)</sup> | 7ZWI/7ZWF/7ZWM |
|  | RUPA-61 | D3 - 1b | ± 10 <sup>a</sup> | Fabra-García<sup>(32)</sup> | - |
|  | RUPA-44 | D3 - 1b | 11 | Fabra-García<sup>(32)</sup>; Kucharska<sup>(15)</sup> | 7UXL/**9N5I** |
|  | RUPA-117 | D3 - 1b | ± 10 <sup>a</sup> | Fabra-García<sup>(32)</sup> | 7UNB |
| Negative controls | PGDM1400 | HIV Env. | n/a | n/a | n/a |
|  | CIS43 | PfCSP | n/a | n/a | n/a |

**Table S2:** Pfs230:Pfs48/45 residue contact table. *Related to Figure 3*.

| Pfs48/45 Domain | Pfs48/45 Residue | BSA (Å<sup>2</sup>) | Pfs230 Residue contacts |
| --- | --- | --- | --- |
| Domain 1 | N38 | 18 | 3108 |
|  | E40 | 25 | 3011, 3013 |
|  | I41 | 60 | 3011, 3112, 3013 |
|  | S42 | 88 | 2986, 2997, 2998, 3011,3012 |
|  | G43 | 34 | 2997 |
|  | F44 | 15 | 2997 |
|  | I45 | 86 | 2984, 3108-3110 |
|  | G46 | 16 | 3110 |
|  | Y47 | 65 | 3110-3114 |
|  | K48 | 49 | 2982, 2997, 3114 |
|  | N50 | 17 | 2993 |
|  | E54 | 48 | 2990, 2993, 3001 |
|  | G55 | 9 | 2993 |
|  | V56 | 90 | 2942, 2991, 2993, 2994 |
|  | H57 | 78 | 2942, 3116, 3118 |
|  | E65 | 34 | 2943, 3118 |
|  | R67 | 30 | 3118 |
|  | S68 | 20 | 3117-3118 |
|  | I69 | 25 | 3116-3118 |
|  | F70 | 56 | 3115-3117 |
|  | C71 | 25 | 3114-3116 |
|  | T72 | 50 | 3113-3115 |
|  | I73 | 16 | 3114 |
|  | H74 | 24 | 3115 |
|  | S75 | 8 | 3112 |
|  | D80 | 17 | 3110 |
|  | N136 | 39 | 3001 |
|  | Y137 | 18 | 2999 |
| Domain 2 | N247 | 13 | 3111 |
|  | K248 | 71 | 3109-3111 |
|  | I249 | 11 | 3108 |
|  | I250 | 17 | 3108 |
| Domain 3 | I348 | 28 | 2839 |
|  | I349 | 11 | 2839 |
|  | F354 | 48 | 2871, 2980 |
|  | Q355 | 25 | 2870 |
|  | I369 | 53 | 2868, 2870, 2977 |
|  | V370 | 9 | 2977 |
|  | Y371 | 11 | 2870, 2871, 2978-2980 |
|  | D373 | 47 | 2980, 3113 |
|  | N377 | 31 | 3115 |
|  | G379 | 25 | 3111, 3112 |
|  | D380 | 26 | 3111, 3112 |
|  | E385 | 12 | 3079 |

**Table S3.** Data collection and refinement statistics for the RUPA-71, 2A2, and 18F25 Fab crystal structures. *Related to Figures 4 and 5*.

|  | RUPA-71 Fab<br>(PDB: 9N7K) | 2A2 Fab<br>(PDB: 9N89) | 18F25 Fab<br>(PDB: 9N6U) |
| --- | --- | --- | --- |
| **Data collection** |  |  |  |
| Space group | P 2<sub>1</sub> 2<sub>1</sub> 2<sub>1</sub> | P 2<sub>1</sub> 2<sub>1</sub> 2<sub>1</sub> | P 2<sub>1</sub> 2<sub>1</sub> 2<sub>1</sub> |
| Cell dimensions |  |  |  |
| a, b, c (Å) | 64.66, 74.55, 204.90 | 59.97, 76.68, 91.87 | 37.64, 81.80, 140.53 |
| a, b, g (°) | 90, 90, 90 | 90, 90, 90 | 90, 90, 90 |
| Resolution (Å)<sup>a</sup> | 48.85 - 2.28<br>(2.36 - 2.28) | 28.51 - 1.20<br>(1.21 - 1.20) | 34.19 - 1.60<br>(1.63 - 1.60) |
| R<sub>sym</sub> or R<sub>merge</sub><sup>b</sup> | 38.1 (214.6) | 6.9 (58.3) | 7.2 (62.2) |
| R<sub>pim</sub><sup>c</sup> | 9.9 (56.2) | 4.2 (35.6) | 4.6 (40.2) |
| I / σI | 7.5 (1.7) | 8.9 (2.1) | 9.8 (1.8) |
| Completeness (%) | 100 (100) | 99.5 (98.2) | 99.4 (98.6) |
| Redundancy | 15.4 (15.2) | 3.5 (3.6) | 3.3 (3.3) |
| **Refinement** |  |  |  |
| Resolution (Å) | 2.28 | 1.20 | 1.60 |
| No. reflections | 711,113 (67,539) | 891,870 (29,484) | 365,949 (17,109) |
| No. unique reflections | 46,105 (4434) | 254,175 (8,275) | 109,994 (5,231) |
| R<sub>work</sub><sup>d</sup> / R<sub>free</sub><sup>e</sup> | 19.1 / 24.1 | 18.6 / 19.4 | 18.8 / 21.3 |
| No. atoms | 7037 | 3708 | 4056 |
| Protein | 6776 | 3343 | 3432 |
| Ligand/ion | 52 | 0 | 20 |
| Water | 209 | 427 | 604 |
| B-factors |  |  |  |
| Protein | 41 | 12 | 17 |
| Ligand/ion | 53<sup>f</sup> | 0 | 36<sup>g</sup> |
| Water | 39 | 20 | 26 |
| R.m.s. deviations |  |  |  |
| Bond lengths (Å) | 0.009 | 0.004 | 0.006 |
| Bond angles (°) | 0.99 | 0.78 | 0.83 |

<sup>a</sup>Values in brackets refer to the highest resolution bin.
<sup>b</sup> $R_{\text{merge}} = \sum_{\text{hkl}} \sum_i |I_{\text{hkl},i} - \langle I_{\text{hkl}} \rangle| / \sum_{\text{hkl}} \langle I_{\text{hkl}} \rangle$
<sup>c</sup> $R_{\text{pim}} = \sum_{\text{hkl}} [1/(N-1)]^{1/2} \sum_i |I_{\text{hkl},i} - \langle I_{\text{hkl}} \rangle| / \sum_{\text{hkl}} \langle I_{\text{hkl}} \rangle$
<sup>d</sup> $R_{\text{work}} = (\sum ||F_o| - |F_c||) / (\sum ||F_o|) - \text{for all data except } R_{\text{free}}$ (see footnote e).
<sup>e</sup>5% of data were used for the R<sub>free</sub> calculation.
<sup>f</sup>Four glycerol and seven 1,2-ethanediol molecules
<sup>g</sup>Four SO<sub>4</sub> ions

**Table S4:** 2A2, 18F25, and RUPA-71 contact table in Kabat numbering. *Related to Figures 4 and 5*.

| Pfs230 Residue | Structural Element | BSA (Å<sup>2</sup>) | 2A2 HC Residue | Structural Element | BSA (Å<sup>2</sup>) | 2A2 KC Residue | Structural Element | BSA (Å<sup>2</sup>) |
| --- | --- | --- | --- | --- | --- | --- | --- | --- |
| H1159 | D4: β2-β3 loop | 37 | N33 | HCDR1 | 16 | S1 | N-term / FR1-IMGT | 27 |
| E1160 | D4: β3 | 14 | N35 | FR2-IMGT | 2 | I2 | N-term / FR1-IMGT | 8 |
| Y1194 | D4: β5 | 27 | W47 | FR2-IMGT | 22 | Q27 | KCDR1 | 73 |
| Q1196 | D4: β5 | 68 | N50 | FR2-IMGT | 19 | S28 | KCDR1 | 8 |
| E1198 | D4: β5-β6 loop | 20 | D52 | HCDR2 | 21 | N30 | KCDR1 | 6 |
| Y1205 | D4: β6 | 34 | H55 | HCDR2 | 58 | D91 | KCDR3 | 8 |
| K1206 | D4: β6-β7 loop | 51 | G57 | HCDR2 | 23 | Y92 | KCDR3 | 80 |
| G1207 | D4: β6-β7 loop | 21 | T58 | HCDR2 | 11 | S93 | KCDR3 | 42 |
| L1208 | D4: β6-β7 loop | 170 | T59 | FR3-IMGT | 60 | S94 | KCDR3 | 74 |
| N1209 | D4: β6-β7 loop | 107 | Y60 | FR3-IMGT | 8 | P95 | KCDR3 | 12 |
| S1210 | D4: β7 | 54 | N61 | FR3-IMGT | 1 | L96 | KCDR3 | 26 |
| V1211 | D4: β7 | 44 | Q62 | FR3-IMGT | 51 |  |  |  |
| T1215 | D4: α1 | 31 | K65 | FR3-IMGT | 14 |  |  |  |
| Q1250 | D4: β10 | 33 | T100 | HCDR3 | 1 |  |  |  |
| T1252 | D4: β10 | 9 | L101 | HCDR3 | 20 |  |  |  |
| Q1255 | D4: β10-β11 loop | 4 | Y102 | HCDR3 | 70 |  |  |  |
| V1256 | D4: β11 | 8 | G103 | HCDR3 | 17 |  |  |  |
| V1257 | D4: β11 | 70 | S105 | HCDR3 | 32 |  |  |  |
| K1259 | D4: β11 | 41 |  |  |  |  |  |  |
| K1261 | D4: β11 | 15 |  |  |  |  |  |  |
| Pfs230 Residue | Structural Element | BSA (Å<sup>2</sup>) | 18F25 HC Residue | Structural Element | BSA (Å<sup>2</sup>) | 18F25 KC Residue | Structural Element | BSA (Å<sup>2</sup>) |
| K1751 | D7: β4-β5 loop (D7-ID) | 36 | Y32 | HCDR1 | 33 | T30 | KCDR1 | 16 |
| V1752 | D7: β4-β5 loop (D7-ID) | 47 | W33 | HCDR1 | 10 | N31 | KCDR1 | 10 |
| I1754 | D7: β4-β5 loop (D7-ID) | 27 | Y52 | HCDR2 | 23 | D32 | KCDR1 | 32 |
| E1756 | D7: β4-β5 loop (D7-ID) | 35 | D54 | HCDR2 | 9 | Y49 | FR2-IMGT | 24 |
| V1801 | D7: β4-β5 loop (D7-ID) | 20 | D56 | HCDR2 | 12 | S50 | KCDR2 | 4 |
| L1802 | D7: β4-β5 loop (D7-ID) | 79 | R58 | HCDR2 | 11 | Y53 | KCDR2 | 32 |
| D1803 | D7: β4-β5 loop (D7-ID) | 52 | R94 | HCDR3 | 2 | Y55 | FR3-IMGT | 7 |
| N1804 | D7: β4-β5 loop (D7-ID) | 7 | L96 | HCDR3 | 27 | H91 | KCDR3 | 28 |
| T1806 | D7: β4-β5 loop (D7-ID) | 32 | Y97 | HCDR3 | 42 | Y92 | KCDR3 | 85 |
| F1807 | D7: β4-β5 loop (D7-ID) | 115 | L98 | HCDR3 | 70 | S93 | KCDR3 | 1 |
| E1808 | D7: β4-β5 loop (D7-ID) | 40 |  |  |  |  |  |  |
| K1809 | D7: β4-β5 loop (D7-ID) | 2 |  |  |  |  |  |  |
| K1856 | D7: β8-β9 loop | 25 |  |  |  |  |  |  |
| D1857 | D7: β8-β9 loop | 1 |  |  |  |  |  |  |
| Pfs48/45 Residue | Structural Element | BSA (Å<sup>2</sup>) | RUPA-71 HC Res. | Structural Element | BSA (Å<sup>2</sup>) | RUPA-71 KC Res. | Structural Element | BSA (Å<sup>2</sup>) |
| K33 | D1: N-term | 28 | E2 | N-term / FR1-IMGT | 47 | K32 | KCDR1 | 40 |
| S35 | D1: N-term | 14 | G26 | HCDR1 | 19 | L46 | FR2-IMGT | 20 |
| S36 | D1: N-term | 11 | F27 | HCDR1 | 14 | Y49 | FR2-IMGT | 57 |
| K88 | D1: β3-β4 loop | 35 | T28 | HCDR1 | 26 | T56 | FR3-IMGT | 79 |
| K89 | D1: β3-β4 loop | 40 | D31 | HCDR1 | 24 | S67 | FR3-IMGT | 39 |
| S90 | D1: β3-β4 loop | 45 | Y32 | HCDR1 | 25 | | | |
| K101 | D1: β4-β5 loop | 81 | D52C | HCDR2 | 48 | | | |
| Q104 | D1: β4-β5 loop | 61 | E53 | HCDR2 | 18 | | | |
| S119 | D1: α1 | 21 | R94 | HCDR3 | 14 | | | |
| Y126 | D1: α1-β7 loop | 47 | G96 | HCDR3 | 24 | | | |
| E127 | D1: β7 | 42 | L97 | HCDR3 | 60 | | | |
| I128 | D1: β7 | 42 | R98 | HCDR3 | 86 | | | |
| E129 | D1: β7 | 61 | W99 | HCDR3 | 59 | | | |
| E130 | D1: β7 | 76 | Y100 | HCDR3 | 96 | | | |
| N131 | D1: β7 | 51 | D100A | HCDR3 | 35 | | | |
| D132 | D1: β7-β8 loop | 71 | S100B | HCDR3 | 69 | | | |
| T133 | D1: β7-β8 loop | 113 | D101 | HCDR3 | 30 | | | |
| N134 | D1: β7-β8 loop | 24 | | | | | | |
| P135 | D1: β7-β8 loop | 26 | | | | | | |
| R140 | D1: β8 | 6 | | | | | | |

**Table S5:**
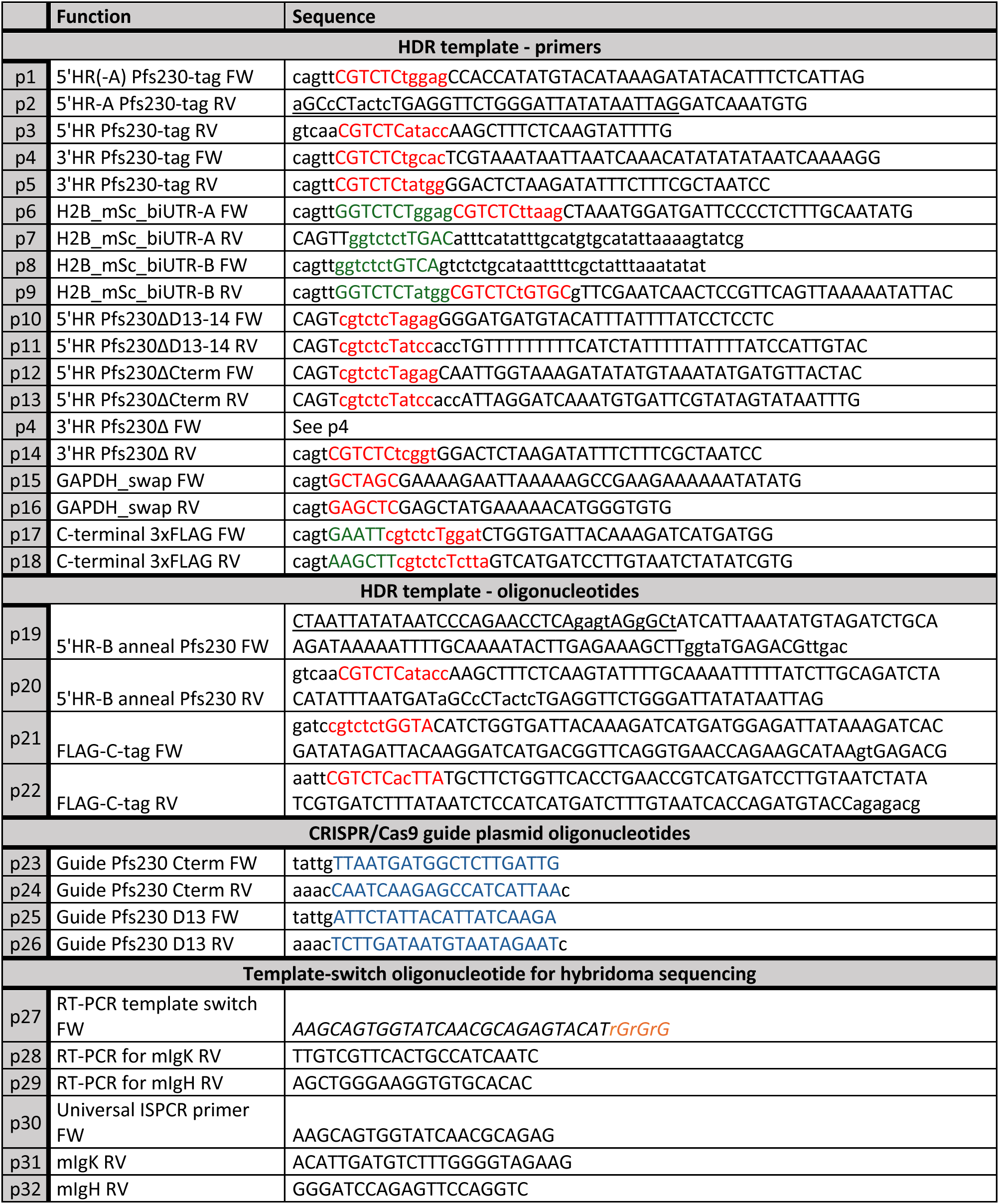
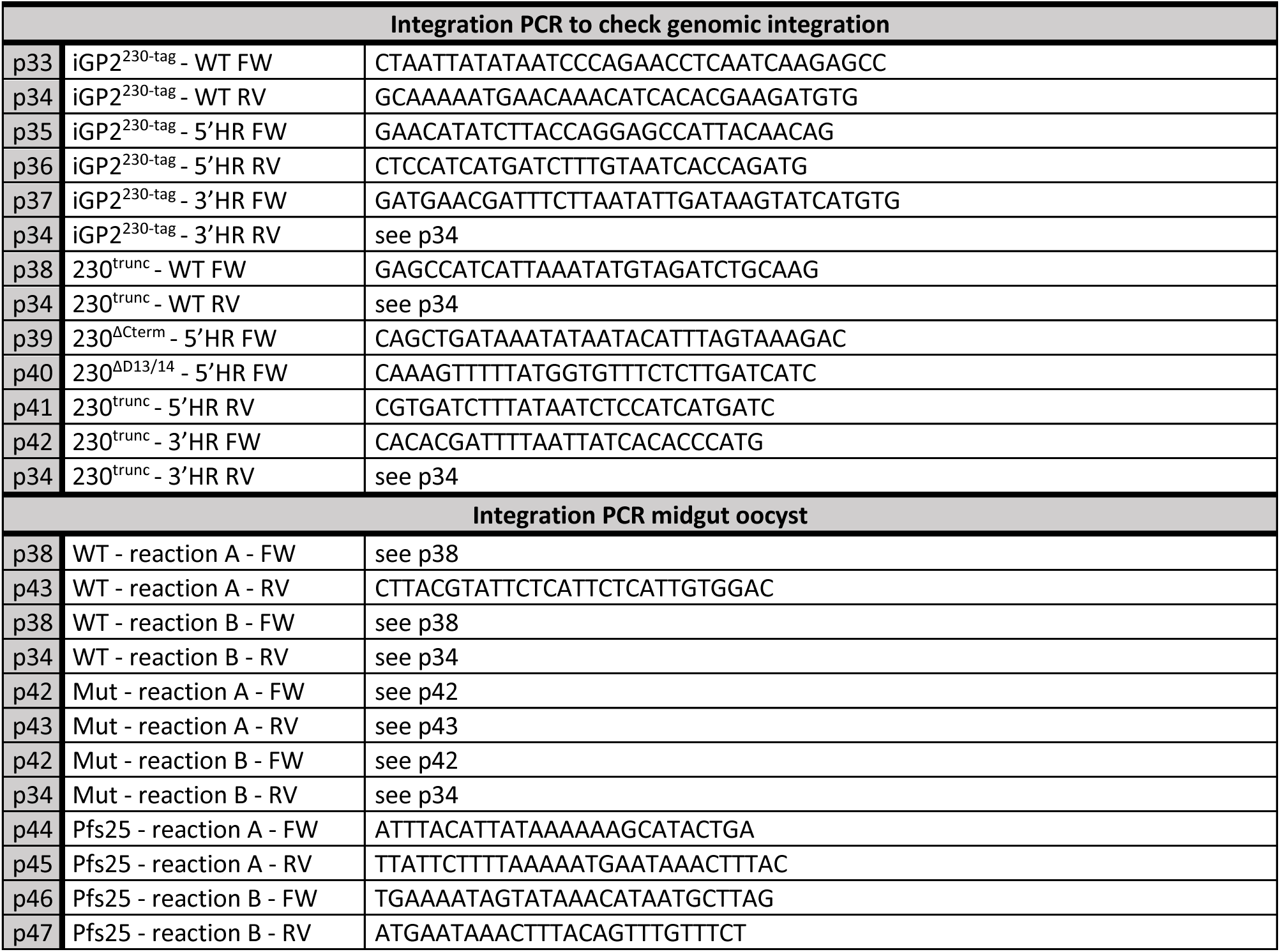
Oligonucleotides used in this study. Red or green nucleotides depict restriction sites used for cloning. Underlined nucleotides were used for overlap-PCR. Blue nucleotides highlight the CRISPR/Cas9 guide RNA sequence. Orange nucleotides are three sequential guanosine nucleotides used for template switching, as described in (*85*). *Related to Methods*.

**Table S6:**
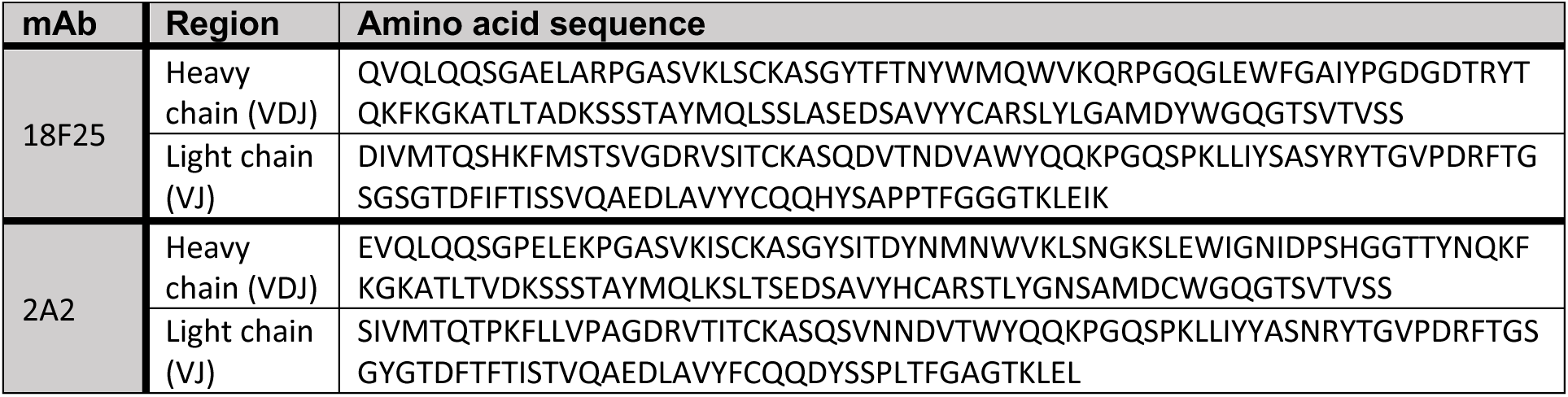
Amino acid sequence of variable chains of mAbs 18F25 and 2A2. *Related to Methods*.

## REFERENCES

1. World malaria report 2024: addressing inequity in the global malaria response. Geneva: World Health Organization; 2024. Contract No.: CC BY-NC-SA 3.0 IGO.

2. Smith DL, McKenzie FE, Snow RW, Hay SI. Revisiting the basic reproductive number for malaria and its implications for malaria control. PLoS Biol. 2007;5(3):e42.

3. Cowman AF, Healer J, Marapana D, Marsh K. Malaria: Biology and Disease. Cell. 2016;167(3):610–24.

4. Aly AS, Vaughan AM, Kappe SH. Malaria parasite development in the mosquito and infection of the mammalian host. Annu Rev Microbiol. 2009;63:195–221.

5. Duffy PE. The Virtues and Vices of Pfs230: From Vaccine Concept to Vaccine Candidate. Am J Trop Med Hyg. 2022;107:17–21.

6. Sauerwein RW, Plieskatt J, Theisen M. 40 Years of Pfs48/45 Research as a Transmission-Blocking Vaccine Target of Plasmodium falciparum Malaria. Am J Trop Med Hyg. 2022;107:22–6.

7. Eksi S, Czesny B, van Gemert GJ, Sauerwein RW, Eling W, Williamson KC. Malaria transmission-blocking antigen, Pfs230, mediates human red blood cell binding to exflagellating male parasites and oocyst production. Mol Microbiol. 2006;61(4):991–8.

8. van Dijk MR, Janse CJ, Thompson J, Waters AP, Braks JA, Dodemont HJ, et al. A central role for P48/45 in malaria parasite male gamete fertility. Cell. 2001;104(1):153–64.

9. Rener J, Graves PM, Carter R, Williams JL, Burkot TR. Target antigens of transmission-blocking immunity on gametes of plasmodium falciparum. J Exp Med. 1983;158(3):976–81.

10. Kumar N. Target antigens of malaria transmission blocking immunity exist as a stable membrane bound complex. Parasite Immunol. 1987;9(3):321–35.

11. Quakyi IA, Carter R, Rener J, Kumar N, Good MF, Miller LH. The 230-kDa gamete surface protein of Plasmodium falciparum is also a target for transmission-blocking antibodies. J Immunol. 1987;139(12):4213–7.

12. Templeton TJ, Kaslow DC. Identification of additional members define a Plasmodium falciparum gene superfamily which includes Pfs48/45 and Pfs230. Mol Biochem Parasitol. 1999;101(1-2):223–7.

13. Gerloff DL, Creasey A, Maslau S, Carter R. Structural models for the protein family characterized by gamete surface protein Pfs230 of Plasmodium falciparum. Proc Natl Acad Sci U S A. 2005;102(38):13598–603.

14. Ko KT, Lennartz F, Mekhaiel D, Guloglu B, Marini A, Deuker DJ, et al. Structure of the malaria vaccine candidate Pfs48/45 and its recognition by transmission blocking antibodies. Nat Commun. 2022;13(1):5603.

15. Kucharska I, Ivanochko D, Hailemariam S, Inklaar MR, Kim HR, Teelen K, et al. Structural elucidation of full-length Pfs48/45 in complex with potent monoclonal antibodies isolated from a naturally exposed individual. Nat Struct Mol Biol. 2025.

16. Dietrich MH, Gabriela M, Reaksudsan K, Dixon MWA, Chan LJ, Adair A, et al. Nanobodies against Pfs230 block Plasmodium falciparum transmission. Biochem J. 2022;479(24):2529–46.

17. Tang WK, Coelho CH, Miura K, Nguemwo Tentokam BC, Salinas ND, Narum DL, et al. A human antibody epitope map of Pfs230D1 derived from analysis of individuals vaccinated with a malaria transmission-blocking vaccine. Immunity. 2023;56(2):433–43.e5.

18. Singh K, Burkhardt M, Nakuchima S, Herrera R, Muratova O, Giås AG, et al. Structure and function of a malaria transmission blocking vaccine targeting Pfs230 and Pfs230-Pfs48/45 proteins. Commun Biol. 2020;3(1):395.

19. Coelho CH, Tang WK, Burkhardt M, Galson JD, Muratova O, Salinas ND, et al. A human monoclonal antibody blocks malaria transmission and defines a highly conserved neutralizing epitope on gametes. Nat Commun. 2021;12(1):1750.

20. Ivanochko D, Fabra-García A, Teelen K, van de Vegte-Bolmer M, van Gemert GJ, Newton J, et al. Potent transmission-blocking monoclonal antibodies from naturally exposed individuals target a conserved epitope on Plasmodium falciparum Pfs230. Immunity. 2023;56(2):420–32.e7.

21. Duffy PE. Transmission-Blocking Vaccines: Harnessing Herd Immunity for Malaria Elimination. Expert Rev Vaccines. 2021;20(2):185–98.

22. Tediosi F, Maire N, Penny M, Studer A, Smith TA. Simulation of the cost-effectiveness of malaria vaccines. Malar J. 2009;8:127.

23. Sherrard-Smith E, Sala KA, Betancourt M, Upton LM, Angrisano F, Morin MJ, et al. Synergy in anti-malarial pre-erythrocytic and transmission-blocking antibodies is achieved by reducing parasite density. Elife. 2018;7.

24. Challenger JD, Olivera Mesa D, Da DF, Yerbanga RS, Lefèvre T, Cohuet A, Churcher TS. Predicting the public health impact of a malaria transmission-blocking vaccine. Nat Commun. 2021;12(1):1494.

25. Vermeulen AN, Roeffen WF, Henderik JB, Ponnudurai T, Beckers PJ, Meuwissen JH. Plasmodium falciparum transmission blocking monoclonal antibodies recognize monovalently expressed epitopes. Dev Biol Stand. 1985;62:91–7.

26. Stone WJR, Campo JJ, Ouédraogo AL, Meerstein-Kessel L, Morlais I, Da D, et al. Unravelling the immune signature of Plasmodium falciparum transmission-reducing immunity. Nat Commun. 2018;9(1):558.

27. van der Kolk M, de Vlas SJ, Sauerwein RW. Reduction and enhancement of Plasmodium falciparum transmission by endemic human sera. Int J Parasitol. 2006;36(10-11):1091–5.

28. Bousema T, Roeffen W, Meijerink H, Mwerinde H, Mwakalinga S, van Gemert GJ, et al. The dynamics of naturally acquired immune responses to Plasmodium falciparum sexual stage antigens Pfs230 & Pfs48/45 in a low endemic area in Tanzania. PLoS One. 2010;5(11):e14114.

29. Graves PM, Carter R, Burkot TR, Quakyi IA, Kumar N. Antibodies to Plasmodium falciparum gamete surface antigens in Papua New Guinea sera. Parasite Immunol. 1988;10(2):209–18.

30. Mulder B, Lensen T, Tchuinkam T, Roeffen W, Verhave JP, Boudin C, Sauerwein R. Plasmodium falciparum: membrane feeding assays and competition ELISAs for the measurement of transmission reduction in sera from Cameroon. Exp Parasitol. 1999;92(1):81–6.

31. Roeffen W, Mulder B, Teelen K, Bolmer M, Eling W, Targett GA, et al. Association between anÜ-Pfs48/45 reactivity and P. falciparum transmission-blocking activity in sera from Cameroon. Parasite Immunol. 1996;18(2):103–9.

32. Fabra-García A, Hailemariam S, de Jong RM, Janssen K, Teelen K, van de Vegte-Bolmer M, et al. Highly potent, naturally acquired human monoclonal antibodies against Pfs48/45 block Plasmodium falciparum transmission to mosquitoes. Immunity. 2023;56(2):406–19.e7.

33. Lennartz F, Brod F, Dabbs R, Miura K, Mekhaiel D, Marini A, et al. Structural basis for recognition of the malaria vaccine candidate Pfs48/45 by a transmission blocking antibody. Nat Commun. 2018;9(1):3822.

34. MacDonald NJ, Nguyen V, Shimp R, Reiter K, Herrera R, Burkhardt M, et al. Structural and Immunological Characterization of Recombinant 6-Cysteine Domains of the Plasmodium falciparum Sexual Stage Protein Pfs230. J Biol Chem. 2016;291(38):19913–22.

35. Tachibana M, Miura K, Takashima E, Morita M, Nagaoka H, Zhou L, et al. Identification of domains within Pfs230 that elicit transmission blocking antibody responses. Vaccine. 2019;37(13):1799–806.

36. Inklaar MR, de Jong RM, Da DF, Hubregtse LL, Meijer M, Teelen K, et al. Pfs230 Domain 12 is a potent malaria transmission-blocking vaccine candidate. bioRxiv. 2024 (preprint):2024.11.09.622785.

37. Inklaar MR, de Jong RM, Bekkering ET, Nagaoka H, Fennemann FL, Teelen K, et al. Pfs230 Domain 7 is targeted by a potent malaria transmission-blocking monoclonal antibody. NPJ Vaccines. 2023;8(1):186.

38. de Jong RM, Meerstein-Kessel L, Da DF, Nsango S, Challenger JD, van de Vegte-Bolmer M, et al. Monoclonal antibodies block transmission of genetically diverse Plasmodium falciparum strains to mosquitoes. NPJ Vaccines. 2021;6(1):101.

39. Simons LM, Ferrer P, Gombakomba N, Underwood K, Herrera R, Narum DL, et al. Extending the range of Plasmodium falciparum transmission blocking antibodies. Vaccine. 2023;41(21):3367–79.

40. Roeffen W, Geeraedts F, Eling W, Beckers P, Wizel B, Kumar N, et al. Transmission blockade of Plasmodium falciparum malaria by anti-Pfs230-specific antibodies is isotype dependent. Infect Immun. 1995;63(2):467–71.

41. Kundu P, Semesi A, Jore MM, Morin MJ, Price VL, Liang A, et al. Structural delineation of potent transmission-blocking epitope I on malaria antigen Pfs48/45. Nat Commun. 2018;9(1):4458.

42. Miura K, Flores-Garcia Y, Long CA, Zavala F. Vaccines and monoclonal antibodies: new tools for malaria control. Clin Microbiol Rev. 2024;37(2):e0007123.

43. Yoo R, Jore MM, Julien JP. Targeting Bottlenecks in Malaria Transmission: Antibody-Epitope Descriptions Guide the Design of Next-Generation Biomedical Interventions. Immunol Rev. 2025;330(1):e70001.

44. Plieskatt J, Ofori EA, Naghizadeh M, Miura K, Flores-Garcia Y, Borbye-Lorenzen N, et al. ProC6C, a novel multi-stage malaria vaccine, elicits functional antibodies against the minor and central repeats of the Circumsporozoite Protein in human adults. Front Immunol. 2024;15:1481829.

45. Tiono AB, Plieskatt JL, Ouedraogo A, Soulama BI, Miura K, Bougouma EC, et al. A randomized first-in-human phase I trial of differentially adjuvanted Pfs48/45 malaria vaccines in Burkinabé adults. J Clin Invest. 2024;134(7):e175707.

46. Healy SA, Anderson C, Swihart BJ, Mwakingwe A, Gabriel EE, Decederfelt H, et al. Pfs230 yields higher malaria transmission-blocking vaccine activity than Pfs25 in humans but not mice. J Clin Invest. 2021;131(7):e146221.

47. Sagara I, Healy SA, Assadou MH, Kone M, Swihart BJ, Kwan JL, et al. Malaria transmission-blocking vaccines Pfs230D1-EPA and Pfs25-EPA in Alhydrogel in healthy Malian adults; a phase 1, randomised, controlled trial. Lancet Infect Dis. 2023;23(11):1266–79.

48. Boltryk SD, Passecker A, Alder A, Carrington E, van de Vegte-Bolmer M, van Gemert GJ, et al. CRISPR/Cas9-engineered inducible gametocyte producer lines as a valuable tool for Plasmodium falciparum malaria transmission research. Nat Commun. 2021;12(1):4806.

49. Roeffen W, Beckers PJ, Teelen K, Lensen T, Sauerwein RW, Meuwissen JH, Eling W. Plasmodium falciparum: a comparison of the activity of Pfs230-specific antibodies in an assay of transmission-blocking immunity and specific competition ELISAs. Exp Parasitol. 1995;80(1):15–26.

50. Dietrich MH, Chan LJ, Adair A, Boulet C, O’Neill MT, Tan LL, et al. Structure of the Pf12 and Pf41 heterodimeric complex of Plasmodium falciparum 6-cysteine proteins. FEMS Microbes. 2022;3:xtac005.

51. Parker ML, Peng F, Boulanger MJ. The Structure of Plasmodium falciparum Blood-Stage 6-Cys Protein Pf41 Reveals an Unexpected Intra-Domain Insertion Required for Pf12 Coordination. PLoS One. 2015;10(9):e0139407.

52. Churcher TS, Blagborough AM, Delves M, Ramakrishnan C, Kapulu MC, Williams AR, et al. Measuring the blockade of malaria transmission--an analysis of the Standard Membrane Feeding Assay. Int J Parasitol. 2012;42(11):1037–44.

53. Abdel Hamid MM, Abdelraheem MH, Acheampong DO, Ahouidi A, Ali M, Almagro-Garcia J, et al. Pf7: an open dataset of Plasmodium falciparum genome variation in 20,000 worldwide samples. Wellcome Open Res. 2023;8:22.

54. Kanoi BN, Nagaoka H, Morita M, Tsuboi T, Takashima E. Leveraging the wheat germ cell-free protein synthesis system to accelerate malaria vaccine development. Parasitol Int. 2021;80:102224.

55. Diebolder CA, Beurskens FJ, de Jong RN, Koning RI, Strumane K, Lindorfer MA, et al. Complement is activated by IgG hexamers assembled at the cell surface. Science. 2014;343(6176):1260-3.

56. Strasser J, de Jong RN, Beurskens FJ, Wang G, Heck AJR, Schuurman J, et al. Unraveling the Macromolecular Pathways of IgG Oligomerization and Complement Activation on Antigenic Surfaces. Nano Leà. 2019;19(7):4787–96.

57. Ho CM, Beck JR, Lai M, Cui Y, Goldberg DE, Egea PF, Zhou ZH. Malaria parasite translocon structure and mechanism of effector export. Nature. 2018;561(7721):70-5.

58. Ho CM, Jih J, Lai M, Li X, Goldberg DE, Beck JR, Zhou ZH. Native structure of the RhopH complex, a key determinant of malaria parasite nutrient acquisition. Proc Natl Acad Sci U S A. 2021;118(35):e2100514118.

59. Schureck MA, Darling JE, Merk A, Shao J, Daggupati G, Srinivasan P, et al. Malaria parasites use a soluble RhopH complex for erythrocyte invasion and an integral form for nutrient uptake. Elife. 2021;10.

60. Zhao J, Makhija S, Zhou C, Zhang H, Wang Y, Muralidharan M, et al. Structural insights into the human PA28-20S proteasome enabled by efficient tagging and purification of endogenous proteins. Proc Natl Acad Sci U S A. 2022;119(33):e2207200119.

61. Di Trani JM, Gheorghita AA, Turner M, Brzezinski P, Ädelroth P, Vahidi S, et al. Structure of the bc(1)-cbb(3) respiratory supercomplex from Pseudomonas aeruginosa. Proc Natl Acad Sci U S A. 2023;120(40):e2307093120.

62. Liang Y, Plourde A, Bueler SA, Liu J, Brzezinski P, Vahidi S, Rubinstein JL. Structure of mycobacterial respiratory complex I. Proc Natl Acad Sci U S A. 2023;120(13):e2214949120.

63. Ho CM, Li X, Lai M, Terwilliger TC, Beck JR, Wohlschlegel J, et al. Bottom-up structural proteomics: cryoEM of protein complexes enriched from the cellular milieu. Nat Methods. 2020;17(1):79–85.

64. Abramson J, Adler J, Dunger J, Evans R, Green T, Pritzel A, et al. Accurate structure prediction of biomolecular interactions with AlphaFold 3. Nature. 2024;630(8016):493-500.

65. van Dijk MR, van Schaijk BC, Khan SM, van Dooren MW, Ramesar J, Kaczanowski S, et al. Three members of the 6-cys protein family of Plasmodium play a role in gamete fertility. PLoS Pathog. 2010;6(4):e1000853.

66. Sanders RW, Moore JP. Virus vaccines: proteins prefer prolines. Cell Host Microbe. 2021;29(3):327–33.

67. Crank MC, Ruckwardt TJ, Chen M, Morabito KM, Phung E, Costner PJ, et al. A proof of concept for structure-based vaccine design targeting RSV in humans. Science. 2019;365(6452):505-9.

68. McLeod B, Mabrouk MT, Miura K, Ravichandran R, Kephart S, Hailemariam S, et al. Vaccination with a structure-based stabilized version of malarial antigen Pfs48/45 elicits ultra-potent transmission-blocking antibody responses. Immunity. 2022;55(9):1680–92.e8.

69. Dickey TH, Gupta R, McAleese H, Ouahes T, Orr-Gonzalez S, Ma R, et al. Design of a stabilized non-glycosylated Pfs48/45 antigen enables a potent malaria transmission-blocking nanoparticle vaccine. NPJ Vaccines. 2023;8(1):20.

70. Correia BE, Bates JT, Loomis RJ, Baneyx G, Carrico C, Jardine JG, et al. Proof of principle for epitope-focused vaccine design. Nature. 2014;507(7491):201-6.

71. Bale JB, Gonen S, Liu Y, Sheffler W, Ellis D, Thomas C, et al. Accurate design of megadalton-scale two-component icosahedral protein complexes. Science. 2016;353(6297):389-94.

72. Dietrich MH, Chmielewski J, Chan L-J, Tan LL, Adair A, Lyons FMT, et al. Cryo-EM structure of endogenous *Plasmodium falciparum* Pfs230 and Pfs48/45 fertilization complex. bioRxiv. 2025:2025.02.13.638205.

73. Brooks SR, Williamson KC. Proteolysis of Plasmodium falciparum surface antigen, Pfs230, during gametogenesis. Mol Biochem Parasitol. 2000;106(1):77–82.

74. Deutsch EW, Bandeira N, Perez-Riverol Y, Sharma V, Carver JJ, Mendoza L, et al. The ProteomeXchange consortium at 10 years: 2023 update. Nucleic Acids Res. 2023;51(D1):D1539–d48.

75. Lee SM, Plieskatt J, Krishnan S, Raina M, Harishchandra R, King CR. Expression and purification optimization of an N-terminal Pfs230 transmission-blocking vaccine candidate. Protein Expr Purif. 2019;160:56–65.

76. Lim MY, LaMonte G, Lee MCS, Reimer C, Tan BH, Corey V, et al. UDP-galactose and acetyl-CoA transporters as Plasmodium multidrug resistance genes. Nat Microbiol. 2016;1:16166.

77. Verhoef JMJ, Bekkering ET, Boshoven C, Hannon M, Proellochs NI, Spruijt CG, Kooij TWA. The role of stomatin-like protein (STOML) in Plasmodium falciparum. bioRxiv. 2024:2024.07.18.604071.

78. Trager W, Jensen JB. Human malaria parasites in continuous culture. Science. 1976;193(4254):673-5.

79. Lambros C, Vanderberg JP. Synchronization of Plasmodium falciparum erythrocytic stages in culture. J Parasitol. 1979;65(3):418–20.

80. Wu Y, Sifri CD, Lei HH, Su XZ, Wellems TE. Transfection of Plasmodium falciparum within human red blood cells. Proc Natl Acad Sci U S A. 1995;92(4):973–7.

81. Crabb BS, Cowman AF. Characterization of promoters and stable transfection by homologous and nonhomologous recombination in Plasmodium falciparum. Proc Natl Acad Sci U S A. 1996;93(14):7289–94.

82. Graumans W, van der Starre A, Stoter R, van Gemert GJ, Andolina C, Ramjith J, et al. AlbuMAX supplemented media induces the formation of transmission-competent P. falciparum gametocytes. Mol Biochem Parasitol. 2024;259:111634.

83. van de Vegte-Bolmer M, Graumans W, Stoter R, van Gemert GJ, Sauerwein R, Collins KA, Bousema T. A porìolio of geographically distinct laboratory-adapted Plasmodium falciparum clones with consistent infection rates in Anopheles mosquitoes. Malar J. 2021;20(1):381.

84. Schindelin J, Arganda-Carreras I, Frise E, Kaynig V, Longair M, Pietzsch T, et al. Fiji: an open-source plaìorm for biological-image analysis. Nat Methods. 2012;9(7):676–82.

85. Stone WJ, Eldering M, van Gemert GJ, Lanke KH, Grignard L, van de Vegte-Bolmer MG, et al. The relevance and applicability of oocyst prevalence as a read-out for mosquito feeding assays. Sci Rep. 2013;3:3418.

86. Ramjith J, Alkema M, Bradley J, Dicko A, Drakeley C, Stone W, Bousema T. Quantifying Reductions in Plasmodium falciparum Infectivity to Mosquitos: A Sample Size Calculator to Inform Clinical Trials on Transmission-Reducing Interventions. Front Immunol. 2022;13:899615.

87. Meyer L, López T, Espinosa R, Arias CF, Vollmers C, DuBois RM. A simplified workflow for monoclonal antibody sequencing. PLoS One. 2019;14(6):e0218717.

88. Rappsilber J, Ishihama Y, Mann M. Stop and go extraction Üps for matrix-assisted laser desorption/ionization, nanoelectrospray, and LC/MS sample pretreatment in proteomics. Anal Chem. 2003;75(3):663–70.

89. Marr CR, Benlekbir S, Rubinstein JL. Fabrication of carbon films with ∼ 500nm holes for cryo-EM with a direct detector device. J Struct Biol. 2014;185(1):42–7.

90. Guo H, Franken E, Deng Y, Benlekbir S, Singla Lezcano G, Janssen B, et al. Electron-event representation data enable efficient cryoEM file storage with full preservation of spatial and temporal resolution. IUCrJ. 2020;7(Pt 5):860-9.

91. Punjani A, Rubinstein JL, Fleet DJ, Brubaker MA. cryoSPARC: algorithms for rapid unsupervised cryo-EM structure determination. Nat Methods. 2017;14(3):290–6.

92. Bepler T, Morin A, Rapp M, Brasch J, Shapiro L, Noble AJ, Berger B. Positive-unlabeled convolutional neural networks for particle picking in cryo-electron micrographs. Nat Methods. 2019;16(11):1153–60.

93. Punjani A, Zhang H, Fleet DJ. Non-uniform refinement: adaptive regularization improves single-particle cryo-EM reconstruction. Nat Methods. 2020;17(12):1214–21.

94. Sanchez-Garcia R, Gomez-Blanco J, Cuervo A, Carazo JM, Sorzano COS, Vargas J. DeepEMhancer: a deep learning solution for cryo-EM volume post-processing. Commun Biol. 2021;4(1):874.

95. Vonrhein C, Flensburg C, Keller P, Sharff A, Smart O, Paciorek W, et al. Data processing and analysis with the autoPROC toolbox. Acta Crystallogr D Biol Crystallogr. 2011;67(Pt 4):293–302.

96. Kabsch W. XDS. Acta Crystallogr D Biol Crystallogr. 2010;66(Pt 2):125–32.

97. McCoy AJ, Grosse-Kunstleve RW, Adams PD, Winn MD, Storoni LC, Read RJ. Phaser crystallographic soïware. J Appl Crystallogr. 2007;40(Pt 4):658–74.

98. Emsley P, Lohkamp B, Scott WG, Cowtan K. Features and development of Coot. Acta Crystallogr D Biol Crystallogr. 2010;66(Pt 4):486–501.

99. Liebschner D, Afonine PV, Baker ML, Bunkóczi G, Chen VB, Croll TI, et al. Macromolecular structure determination using X-rays, neutrons and electrons: recent developments in Phenix. Acta Crystallogr D Struct Biol. 2019;75(Pt 10):861–77.

100. Krissinel E, Henrick K. Inference of macromolecular assemblies from crystalline state. J Mol Biol. 2007;372(3):774–97.

101. Pettersen EF, Goddard TD, Huang CC, Meng EC, Couch GS, Croll TI, et al. UCSF ChimeraX: Structure visualization for researchers, educators, and developers. Protein Sci. 2021;30(1):70–82.

102. Jumper J, Evans R, Pritzel A, Green T, Figurnov M, Ronneberger O, et al. Highly accurate protein structure prediction with AlphaFold. Nature. 2021;596(7873):583–9.

103. Croll TI. ISOLDE: a physically realistic environment for model building into low-resolution electron-density maps. Acta Crystallogr D Struct Biol. 2018;74(Pt 6):519–30.

104. Chaudhury S, Lyskov S, Gray JJ. PyRosetta: a script-based interface for implementing molecular modeling algorithms using Rosetta. Bioinformatics. 2010;26(5):689–91.

105. Adams PD, Afonine PV, Bunkóczi G, Chen VB, Davis IW, Echols N, et al. PHENIX: a comprehensive Python-based system for macromolecular structure solution. Acta Crystallogr D Biol Crystallogr. 2010;66(Pt 2):213–21.

106 . Morin A, Eisenbraun B, Key J, Sanschagrin PC, Timony MA, Ottaviano M, Sliz P. Collaboration gets the most out of soïware. Elife. 2013;2:e01456.

107. Li H. A statistical framework for SNP calling, mutation discovery, association mapping and population genetical parameter estimation from sequencing data. Bioinformatics. 2011;27(21):2987–93.

108. Ranford-Cartwright LC, Balfe P, Carter R, Walliker D. Genetic hybrids of Plasmodium falciparum identified by amplification of genomic DNA from single oocysts. Mol Biochem Parasitol. 1991;49(2):239–43.

